# Uridine as a potentiator of aminoglycosides through activation of carbohydrate transporters

**DOI:** 10.1101/2023.07.31.551273

**Authors:** Manon Lang, Stéphane Renard, Imane El-Meouche, Ariane Amoura, Erick Denamur, Tara Brosschot, Molly Ingersoll, Eric Bacqué, Didier Mazel, Zeynep Baharoglu

**Affiliations:** Institut Pasteur, Université Paris Cité, CNRS UMR3525, Unité Plasticité du Génome Bactérien, 75015 Paris, France; Sorbonne Université, Collège Doctoral, F-75005 Paris, France; Evotec ID (Lyon) SAS, 69000 Lyon, France; Université Paris Cité, INSERM UMR 1137 IAME, F-75018 Paris, France; Université Paris Cité, Institut Cochin, INSERM U1016, CNRS UMR 8104, Paris, 75014, France; Mucosal Inflammation and Immunity, Department of Immunology, Institut Pasteur, 75015 Paris, France and Inserm U1223, Paris, France

## Abstract

Aminoglycosides (AGs) are broad-spectrum antibiotics effective against Gram-negative bacteria. AG uptake depends on membrane potential, but the precise mechanisms are incompletely understood. We report here a new mechanism of active AG uptake in Gram-negative bacteria. In *E. coli*, overexpression of various carbohydrate transporters increases susceptibility to AGs. Conversely, deletion of a single transporter has little impact. We propose a new uptake model where AGs act as substrates for redundant carbohydrate transporters. This mechanism appears to be shared among Gram-negative ESKAPE pathogens. We screened for molecules that induce transporters’ expression and identified uridine. When uridine is co-administered with AGs under conditions mimicking urinary tract infections, the efficacy of AG therapies is significantly improved against *E. coli*, including resistant strains, due to enhanced bacterial uptake. Based on previous knowledge on the use of uridine in humans, we propose that uridine can be a potentiating adjuvant to AG treatment of infectious diseases in the hospital.

## Introduction

Antibiotics have played a crucial role in modern medicine. Their use is jeopardized by the emergence of resistance in bacterial populations ^1^, with a prevalence of Gram-negative bacteria as highlighted by the World Health Organization ^2^. This is primarily due the Gram-negative double membrane barrier ^3^. Bacterial antimicrobial resistance (AMR) was directly linked with 1.27 million deaths in 2019, with only six pathogens accounting for 95% of these deaths, namely *Escherichia coli*, *Staphylococcus aureus*, *Klebsiella pneumoniae*, *Acinetobacter baumannii*, and *Pseudomonas aeruginosa* ^4^. Among these, all except *S. aureus* are Gram-negative bacteria. AMR-related fatalities thus surpass the combined death toll of malaria and HIV ^4^, underscoring the urgent need for the development of novel therapies.

Aminoglycosides (AGs) are a class of broad-spectrum antibiotics that effectively cross the double membrane barriers of Gram-negative bacteria. They are commonly used in clinical settings to treat infections caused by various Gram-negative pathogens, such as pneumonia, sepsis, and urinary tract infections (UTIs) ^5^. However, AG treatment is associated with side effects like ototoxicity ^6^ and nephrotoxicity ^7,8^ in case of long-term treatments.

AGs entry into Gram-negative cells is believed to occur in several steps ^9,10^. First, electrostatic interactions between AGs and the outer membrane ^9,11,12^ leads to membrane disruption and AG entry into the periplasm ^13^. Outer membrane porins have also been observed to be involved in this initial step ^14^. The second step allows AG diffusion from the periplasm into the cytoplasm, and relies on the proton motive force (PMF), which is directly linked with electron transport and respiration ^15,16^. Finally, AGs reach their target, the ribosome, and impair translation^17,18^. Mistranslated membrane proteins further compromise membrane integrity, leading to additional AG uptake in a third step.

In a previous study on factors influencing the response of *Vibrio cholerae* to AGs, we identified a non-coding RNA called *ctrR* (<u>c</u>arbohydrates transport regulating RNA). Deletion of *ctrR* increased resistance to AGs and reduced their uptake^19^. *ctrR* and homologous RNAs in other *Vibrio* spp target and stabilize the mRNAs of several carbohydrate transporters ^20^, including the *V. cholerae* phosphotransferase system (PTS) *VC1826-27*, which is involved in fructose uptake, the maltose ABC transporter *malEFG*, and the maltoporin *lamB*. Deletion of these carbohydrate transporters resulted in decreased susceptibility to tobramycin and reduced uptake. This led us to envisage a general connection between sugar transporters and AG uptake. Consistent with this hypothesis, carbon sources also impact the minimum inhibitory concentration (MIC) of tobramycin in *E. coli*, *P. aeruginosa*, *K. pneumoniae* and *A. baumannii* ^19^.

In the present study, we identify the specific carbohydrate transporters involved in AG uptake in *E. coli* and demonstrate that this uptake is not dependent on changes in the PMF. Thus, we provide evidence for a previously unreported mechanism of AG uptake in Gram-negative bacteria other than *V. cholerae*. We propose that the sugar moiety inherent to AG structure, allows them to be recognized as substrates by several carbohydrate transporters in Gram-negative pathogens, enabling AGs to exploit bacterial sugar transport systems. We screen for molecules capable of increasing the number of these transporters both in rich medium and synthetic human urine, and identify uridine as a potentiator of AGs. Uridine supplementation increases the number of AG transporters and offers a means to reduce effective AG doses and to re-sensitize resistant strains in clinics.

## Results

### *cmtA* deletion decreases susceptibility to AGs

We addressed whether like *V. cholerae, E. coli* exhibits phenotypes connecting AG uptake and carbohydrate transporters. Firstly, we generated single deletion mutants of carbohydrate transporters in *E. coli* and examined the impact on tobramycin susceptibility (Table S1).

Among 23 mutants tested, Δ*cmtA* displayed a 4-fold increase in the Minimal Inhibitory Concentration (MIC) of tobramycin compared to the wild-type (WT) strain (Figure 1A), and better growth in the presence of tobramycin (Figure 1B). CmtA is involved in mannitol transport ^21^ and shares 52% similarity with the mannitol transporter MtlA^22^. *ΔcmtA* also showed an increased MIC for other AGs (kanamycin, gentamicin and neomycin); but not for other antibiotic families (trimethoprim, ciprofloxacin, amoxicillin, or chloramphenicol) (Table 1), including spectinomycin. Spectinomycin is an aminocyclitol antibiotic closely related to AGs but lacking the amino-sugar moiety. Hence, the decreased susceptibility of *ΔcmtA* was AG-specific. The other transporter deletion mutants showed no impact or a slight increase in the MIC of tobramycin (Table S1).

**Figure 1:**
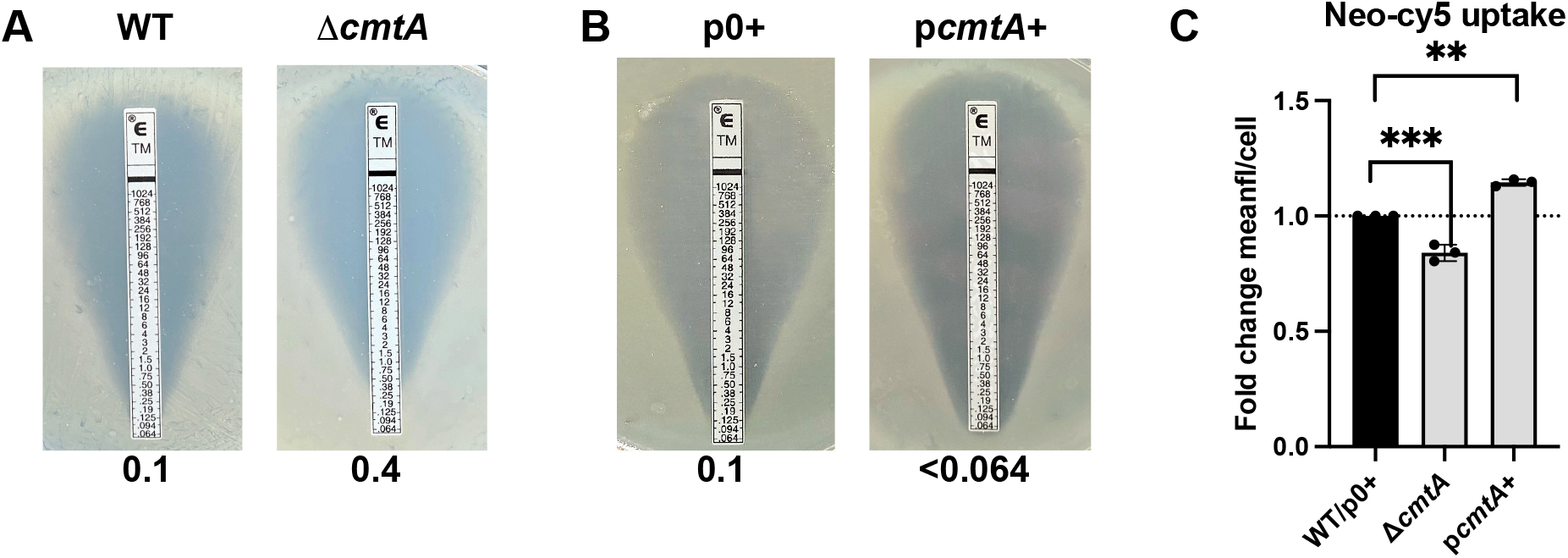
CmtA is involved in tobramycin tolerance. **A**. MIC of tobramycin, indicated in µg/ml, measured using Etests in *E. coli* WT and *ΔcmtA.* **B**. MIC of tobramycin, indicated in µg/ml, measured using Etests on *E. coli* carrying the empty vector (p0+) compared to the vector overexpressing CmtA (p*cmtA*+). Sodium benzoate 1 mM as inducer was added on the medium. **C.** Uptake of Neo-cy5 evaluated by flow cytometry on *E. coli* Δ*cmtA* compared to the WT strain; and *E. coli* carrying a plasmid overexpressing CmtA compared to the strain carrying the empty vector (p0), expressed as fold change of mean fluorescence per cell (compared either to the WT for mutants or to the empty vector for overexpression). For statistical significance calculations, we used one-way ANOVA. **** means p<0.0001, ** means p<0.01. Number of replicates for each experiment: n=3

**Table 1:** MIC (µg/ml) of different antibiotics. determined by Etest in MH medium for all antibiotics except for neomycin (microtiter broth dilution method). (Tob: Tobramycin; Kan: kanamycin; Gen: gentamicin; Neo: neomycin; Spec: spectinomycin; Cip: ciprofloxacin; Trim: trimethoprim; Amox: amoxicillin; Chlo: chloramphenicol). R: plasmid resistance.

| MIC | Tob | Kan | Gen | Neo | Spec | Cip | Trim | Amox | Chlo |
| --- | --- | --- | --- | --- | --- | --- | --- | --- | --- |
| WT | 0.1 | 1 | 0.1 | 1 | 4 | 0.008 | 0.4 | 6 | 3 |
| $\Delta cmtA$ | 0.4 | 2 | 0.4 | 2 | 4 | 0.008 | 0.4 | 6 | 4 |
| p0+ | 0.1 | R | 0.064 | R | 6 | 0.008 | 0.5 | 6 | 1.5 |
| pcmtA+ | >0.064 | R | 0.02 | R | 6 | 0.008 | 0.5 | 6 | 1.5 |

The lack of a pronounced tobramycin resistance phenotype in the single deletion strains might be attributed to compensatory effects among these transporters, some of which share predicted substrates (Table S1). Consequently, we decided to investigate the effect of each transport system by overexpressing them in trans.

### Overexpression of various carbohydrate transporters increases susceptibility to AGs

We expressed the *cmtAB* PTS transporter gene from an inducible promoter to evaluate its response to AGs and non-AG antibiotics as above. Overexpression of the transporter enhanced susceptibility to tobramycin when compared to the empty vector (Figure 1B and Table 1). This response was again specific to AGs (Table 1). Subsequently, we cloned 23 identified carbohydrate transporters on the same inducible plasmid. Among these, 16 demonstrated increased susceptibility to tobramycin compared to the empty vector (Figure S1A and Table S1), and this was AG-specific for 11 transporters: *cmtAB*, *chbCB* (chitobiose PTS), *srlEAB* (glucitol PTS), *ascF* (cellobiose/arbutine PTS), *malEFG (*maltose ABC transporter), *fruBKA (fructose PTS)*, *frwBC* (fructose-like PTS), *mngA* (fructose-like PTS), *ypdGH* (fructose-like PTS), *bglF* (β-glucoside PTS), and *malX* (maltose PTS) (Figure S1B). Five overexpression strains became susceptible AGs, but also to ciprofloxacin or carbenicillin (Figure S1B). These observations suggest the involvement of multiple carbohydrate transporters in the susceptibility to AGs under our experimental conditions.

### Carbohydrate transporters promote AG uptake

To address whether differential expression of carbohydrate transporters modulates AG uptake, we used Neo-cy5, a fluorescent AG specifically designed for bacterial uptake studies^23^. Neo-cy5 mimics the properties of AGs in terms of uptake, mode of action, and activity against Gram-negative bacteria^24^. Deletion of *cmtA* reduced intracellular fluorescence following treatment, indicating a decrease in AG uptake (Figure 1C - Figure S2A and S2C), while CmtAB overexpression led to increased Neo-cy5 uptake (Figure 1C), confirming that CmtAB is involved in AG uptake.

We next addressed whether AG uptake through carbohydrate transporters occurs in other Gram-negative pathogens, namely *P. aeruginosa* and *A. baumannii*. In *P. aeruginosa*, we overexpressed five of its transporters involved in glucose, mannose, maltose, and fructose uptake. This resulted in up to 10-fold increased susceptibility to AGs (tobramycin and gentamicin, Figure S3A and Table S1), but not to other antibiotic families (Table S1). To confirm increased AG uptake, we measured Neo-cy5 fluorescence on the strain overexpressing *mtlFGK*, demonstrating a 1.3-fold higher uptake of AGs compared to the empty vector after 15 minutes of sub-MIC treatment (Figure S3B). For *A. baumannii*, overexpression of the PTS *fruA* also decreased the MIC of tobramycin to <0.064 compared to 0.1 with the empty vector (Table S1).

Taken together, these observations support the hypothesis that redundant transporters play a role in AG uptake and suggest a strategy where therapeutic efficacy of AGs would be enhanced through increased expression of selected transporters.

### Uridine activates the expression of *ctmA*

We set out to identify conditions that could strongly induce the expression of selected carbohydrate transporters and subsequently enhance AG uptake. As a proof-of-concept, we chose to use the CmtAB transporter. The regulation of sugar transporters is known to involve CRP (cyclic AMP Repressor Protein), the primary regulator of carbon catabolic repression ^25^. We constructed a plasmid carrying a transcriptional fusion between the *cmtAB* operon promoter and GFP and to confirmed that that *cmtAB* is controlled by CRP (Figure S4A). To explore potential carbohydrate sources that could activate the expression of the *cmtA* promoter, we used this fluorescent tool to screen a total of 198 molecules in plates from the Biolog Phenotype Microarray system. We inoculated these plates with WT *E. coli* carrying the P*cmtA*-GFP plasmid and monitored both growth at OD_600nm_ and GFP fluorescence over an 8-hour period (Table S2). By calculating the fluorescence to growth ratio, we identified uridine as the most potent activator of P*cmtA*-GFP (ratio of of 9959) compared to glucose which had one of the lowest levels of *cmtA* expression (ratio of 698) (Table S2). Uridine was followed by bromo-succinic acid and inosine. We also screened a plate with nucleosides and nucleotides, uridine once again showed the highest activation ratio (7601), followed by inosine (6235), guanosine (5669), cytidine (5177), adenosine (5154), thymidine (4152), xanthosine (3907), and finally glucose as the control (ratio of 839) (Table S2).

To validate the findings from high throughput Biolog screens, we measured the effect of uridine on fluorescence in standard media, using both a plate reader (Figure 2A) and flow cytometry (Figure 2B and Figure S2B). Consistently, a six-fold increase in fluorescence production was observed when the media was supplemented with uridine compared to the condition without additional carbohydrates. Uridine consists of a sugar moiety, ribose, and a nucleotide moiety, uracil. The induction of the *cmtA* promoter was observed neither with ribose alone, nor with uracil alone (Figure 2B).

**Figure 2:**
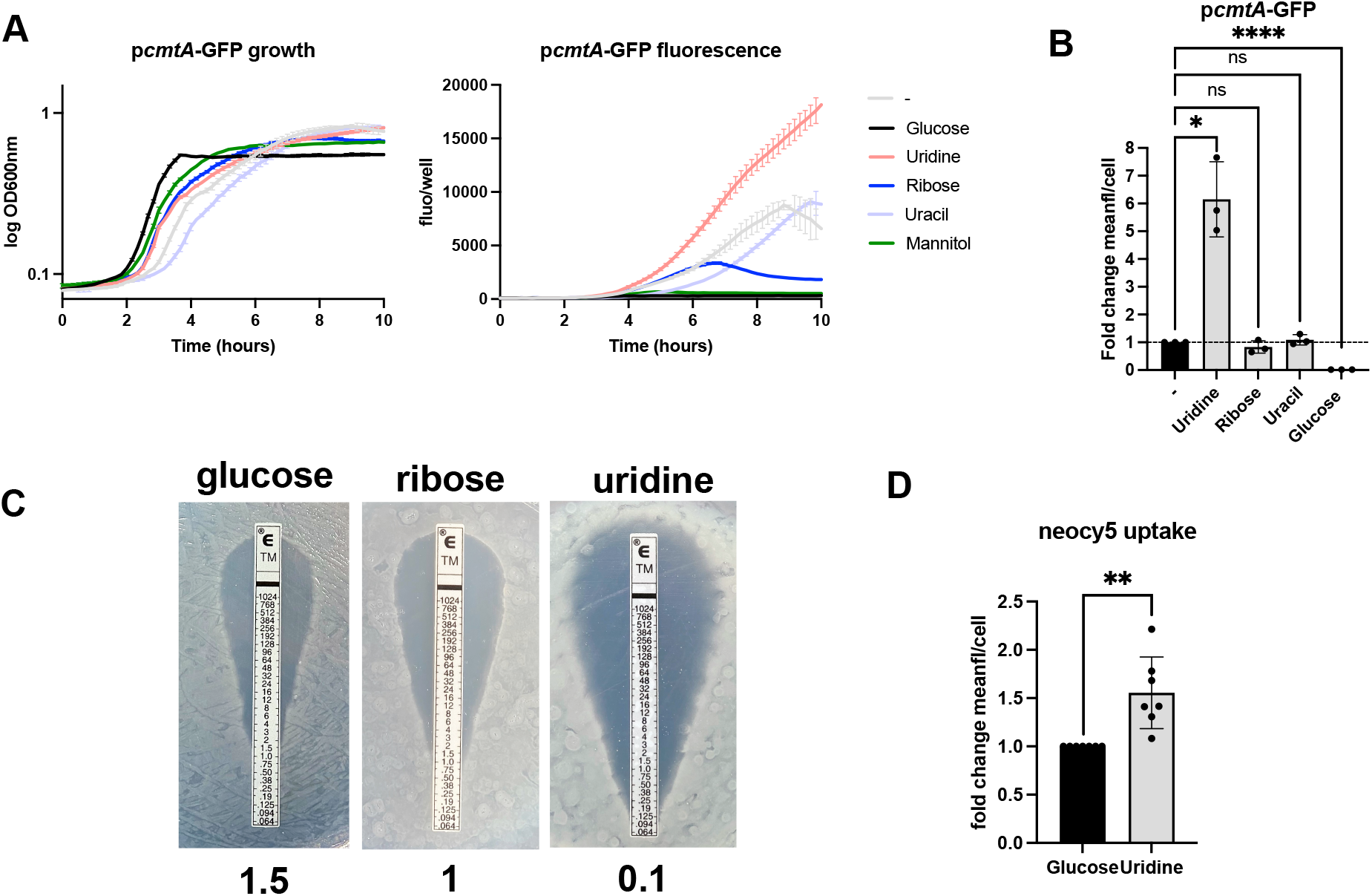
Uridine increases *cmtA* expression and decreases the MIC of tobramycin by enhancing its uptake. **A**. GFP expression from *cmtA* promoter depending on the condition (supplemented or not with glucose, uridine, ribose, uracile or mannitol 0.5%). Growth depending on the molecule is presented on the right by following OD 600 nm. Mean of three biological replicate and standard deviation are represented. **B.** Quantification of GFP fluorescence using flow cytometry depending on the substrate added in the medium (supplemented or not with glucose, uridine, ribose 0.5%, or uracil 0.1%) expressed as fold change of mean fluorescence per cell compared with the no molecule condition. For statistical significance calculations, we used one-way ANOVA. **** means p<0.0001, * means p<0.05. ns: non-significant. Number of replicates for each experiment: n=3. **C**. Tobramycin MIC of *E. coli* WT determined by Etest, expressed in µg/ml, depending on the molecule added in the medium (glucose, ribose or uridine 0.5%). **D.** Uptake of Neo-cy5 evaluated by flow cytometry on *E. coli* WT grown either with a supplementation of glucose (MIC 1.5 µg/ml) or uridine (MIC 0.1 µg/ml) 0.5%, expressed as fold change of mean fluorescence per cell compared to glucose. For statistical significance calculations, we used t-test. ** means p<0.01.

We next asked whether uridine can activate the promoter of another carbohydrate transporter involved in AG uptake, specifically *fruA*. FruA is the only conserved PTS system present in both enterobacteria and *Pseudomonadales*, annotated as a fructose-specific transporter. The p*fruA*-GFP construct also exhibited increased fluorescence when uridine was added to the medium (Figure S5A). To rule out any potential pleiotropic effects of uridine on gene expression under our experimental conditions, we tested the effect of uridine on BtuB, the transporter for vitamin B12. The P*btuB*-GFP construct did not show increased GFP expression when the medium was supplemented with uridine (Figure S5B). These results suggest that uridine has the ability to specifically activate transcription from multiple sugar transporters.

### Uridine supplementation decreases the MIC of aminoglycosides by increasing their uptake

Since uridine enhances the expression of carbohydrate/AG transporters, its addition in growth media should increase bacterial AG uptake, resulting in a lower dose of AGs to kill bacteria, i.e. reduced MIC. We measured MIC values of various antibiotics in *E. coli*, in the presence of different carbohydrate sources (Table S1). In the case of AGs, supplementation with uridine decreased the MIC of tobramycin and gentamicin by 10-fold compared to glucose as a baseline. Glucose is the preferential carbon source in *E. coli* and its presence decreases the expression of non-glucose sugar transporters in a process called carbon catabolic repression, involving CRP as mentioned above ^26^. No increase of susceptibility was observed to amoxicillin or chloramphenicol (Table S1). Unlike with uridine, no significant decrease in MIC was observed upon supplementation with uridine’s sugar moiety, ribose (Figure 2C and Table S1).

Supplementation with uridine also decreased the MIC of tobramycin compared to glucose for *V. cholerae* (10-fold) and ESKAPE pathogens *K. pneumoniae* (4-fold), *P.* aeruginosa (4-fold) and A. *baumannii* (8-fold) (Table S1). Thus, the involvement of carbohydrate transporters in AG uptake is a shared characteristic among Gram-negative bacteria, extending beyond a single genus. The potentiating effects of uridine appear to be present in the pathogens we examined.

Similar to uridine, supplementation with cytidine, adenosine, and thymidine also resulted in increased susceptibility to AGs (MIC of 0.1-0.4 µg/ml) compared to glucose supplementation, whereas inosine did not (MIC of 1 µg/ml) (Table S1). Guanosine was poorly soluble.

We confirmed that uridine increases susceptibility to AGs through increasing AG uptake, using Neo-cy5 fluorescence per cell, which varied from 1.92 with glucose to 2.82 with uridine after a 15-minute sub-MIC tobramycin treatment (Figure 2D and Figure S2C). In addition, the effect of uridine on AG uptake was not due to changes in membrane potential (Figure S6A) although it has to be maintained to observe a potentiating effect (Figure S6B). We also tested and confirmed that uridine mediated AG susceptibility is not related to uridine catabolism (Table S1, Figure S7A) or stress responses (Figure S7B). Altogether, our data supports that uridine supplementation leads to a higher AG uptake through carbohydrate transporters.

### Uridine induces rapid death and prevents the selection of resistant mutants

In order to evaluate the impact of different carbon sources on the effectiveness of tobramycin, we quantified the kinetics of cell death by measuring survival rates during several times points after lethal tobramycin treatment. When glucose (MIC 1 µg/ml, Table S1) was supplemented in the medium, the bactericidal effect of tobramycin was abolished, despite administering the drug at a concentration 10 times higher than the MIC (Figure S6). In contrast, supplementation with maltose (MIC 0.4 µg/ml, Table S1) or uridine (MIC 0.1 µg/ml, Table S1) enhanced the bactericidal effect of tobramycin to varying degrees (Figure S8A), with maltose showing the least effectiveness. Notably, in the maltose-treated bacteria, regrowth was observed after 20 hours of treatment, due to the selection of *fusA* or *rplL* resistant mutants (Figure S8B), already known to be involved in AG resistance mechanisms ^27,28^.

On the other hand, tobramycin showed the highest efficacy on uridine-treated bacteria. The rapid bactericidal effect induced by uridine supplementation may have prevented the selection of these mutations in the bacterial population.

### Uridine potentiates AG efficiency *on E. coli* in synthetic urine and plasma

Urinary tract infections (UTIs) are a commonly caused by Gram-negative bacteria, and treated using AGs in hospital settings. We thus tested the effects of uridine on AG susceptibility in synthetic human urine medium ^29^. The MIC of tobramycin in this medium was enhanced 10-fold with addition of 0.5% of uridine (1 µg/mL with uridine vs. 10 µg/mL without). Next, we compared the effect of different concentrations of uridine, ranging from 1% to 0% in 2-fold dilutions, on bacterial survival after 20 hours of tobramycin treatment (Figure 3A). Bacterial cultures with a density of 10^7^ CFU/mL were treated with a tobramycin concentration at 50% of the MIC with uridine, i.e. 0.5 µg/mL. We observed the most significant AG potentiation effect with 0.031% uridine. Interestingly, the potentiating effect increased between uridine concentrations of 0.0009% and 0.031%, but decreased from 0.031% to 1%. This observation suggests a possible competition between uridine and tobramycin for the transporter responsible for the entry of AGs. Note that a concentration of 0.031% uridine can be achieved in the human body, indicating its promising potential as an adjuvant to enhance AG treatment ^30^.

**Figure 3:**
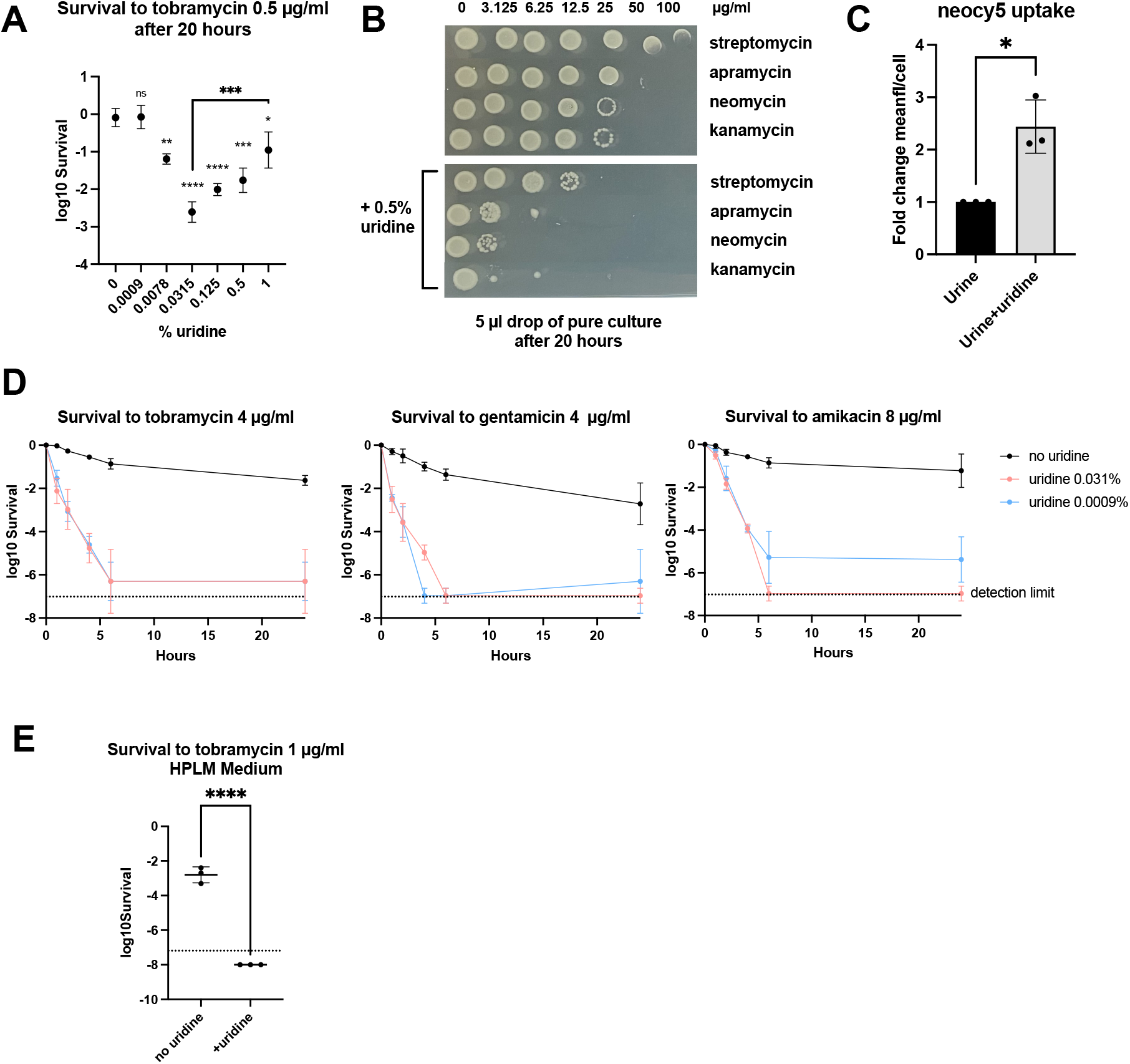
Uridine potentiates AGs in urine and plasma synthetic media by enhancing uptake. **A**. Survival of *E. coli* after 20 hours of treatment with low-dose tobramycin (0.5 µg/ml) according to the addition of different concentrations of uridine (shown in %). Significance above each point represents the significance of the decrease in survival induced by the addition of the uridine percentage mentioned below compared with the no uridine condition (0%), except for the comparison between 0.0315% and 1% as indicated. For statistical significance calculations, we used one-way ANOVA. **** means p<0.0001, *** means p<0.001, ** means p<0.01, * means p<0.05. ns: non-significant. Number of replicates: n=3. **B.** Liquid MIC test performed on *E. coli*, in synthetic urine medium by drop test as indicated in the method section, using concentration of the AG from 0 to 100 µg/ml, with or without addition of 0.5% uridine in the medium. **C.** Neocy5 uptake evaluated by flow cytometry on *E. coli* growing in synthetic urine medium supplemented or not with uridine 0.5%, expressed as fold change of mean fluorescence per cell compared with no supplementation. For statistical significance calculations, we used t-test. * means p<0.05. ns: non-significant. Number of: replicates n=3. **D**. Survival of *E. coli* in urine synthetic medium, after addition of 4 µg/ml of tobramycin, 4 µg/ml of gentamicin or 8 µg/ml of amikacin in the medium supplemented or not with 0.031% or 0.0009% of uridine. Survival was assessed by plating and counting CFU/ml at time 0, 1, 2, 4, 6 and 24 hours after treatment. The dotted line indicates the limit of detection (no colony on the pure culture plate, counted as 1 CFU). Geometric mean and geometric standard deviation on three biological replicates are represented. **E.** Survival of *E. coli* K12 after 20 hours of treatment with 1 µg/ml of tobramycin supplemented or not with 0.031% of uridine. The dotted line indicates the limit of detection (no colony on the pure culture plate, counted as 1 CFU). For statistical significance calculations, we used t-test. **** means p<0.0001. Number of replicates: n=3

Liquid MIC assays were performed with other AGs, such as streptomycin, neomycin, apramycin and kanamycin. The addition of uridine was found to decrease the MIC for all four AGs (Figure 3B). Neo-cy5 assay demonstrated approximately 2.5-fold higher uptake at the single-cell level in uridine-supplemented synthetic urine medium, confirming enhanced AG uptake (Figure 3C and Figure S2C). For cell death kinetics experiments, uridine concentrations of 0.0009% (0.036 mM) or 0.031% (1.27 mM) were used in combination with tobramycin and gentamicin (4 µg/ml), as well as amikacin (8 µg/ml), three commonly used clinical AGs. The addition of uridine resulted in rapid eradication of bacteria, as evidenced by a significant decrease in survival after 24 hours, compared to only a one to two log decrease when uridine was absent from the medium (Figure 3D).

Note that the concentrations of AGs used in these experiments were lower than those typically achieved in the bladder during therapy. In humans, the administration of 1 mg/kg of gentamicin leads to urinary concentrations ranging from 113 to 423 µg/ml after 1 hour of treatment, and 12 to 271 µg/ml after 2 hours ^31^. Despite this, the addition of uridine still demonstrated a substantial effect, suggesting that the use of uridine for urinary tract infections could theoretically be feasible.

To investigate the potential use of uridine in treating blood infections, e.g. sepsis, we conducted experiments using a plasma-like medium (HPLM), with or without the addition of 0.031% uridine. Tobramycin was highly efficient in this medium since no survival was monitored at 2 µg/ml of tobramycin after 20 hours of treatment. Remarkably, uridine supplementation decreased survival to 1 µg/ml of tobramycin (Figure 3E). Note that this medium also contains other carbohydrate sources, but they did not interfere with the potentiating action of uridine in this context.

These findings provide further support for the hypothesis that uridine has the potential to enhance the effectiveness of AG therapy in the context of blood infections.

### Uridine potentiation of tobramycin is effective on clinical *E. coli* strains

Next, we used synthetic urine medium to assess the AG potentiating action of uridine on clinical strains of *E. coli,* isolated from a long-stay hospital (Table S3), and for which the complete genome sequences are available. We first tested four strains that do not carry any known tobramycin resistance gene: strain 886 with no resistance genes; strains Ec019 and Ec068 with the MdfA efflux pump associated with a 2-3 fold increase in AG MICs ^32^; and strain 1236 which harbors resistance genes against various other antibiotics (ß-lactams, sulfonamides, spectinomycin, streptomycin). In all four strains, the addition of uridine (at 0.031 %) resulted in strongly decreased survival to 20 hours lethal tobramycin treatment (10 µg/ml) (Figure 4A).

**Figure 4:**
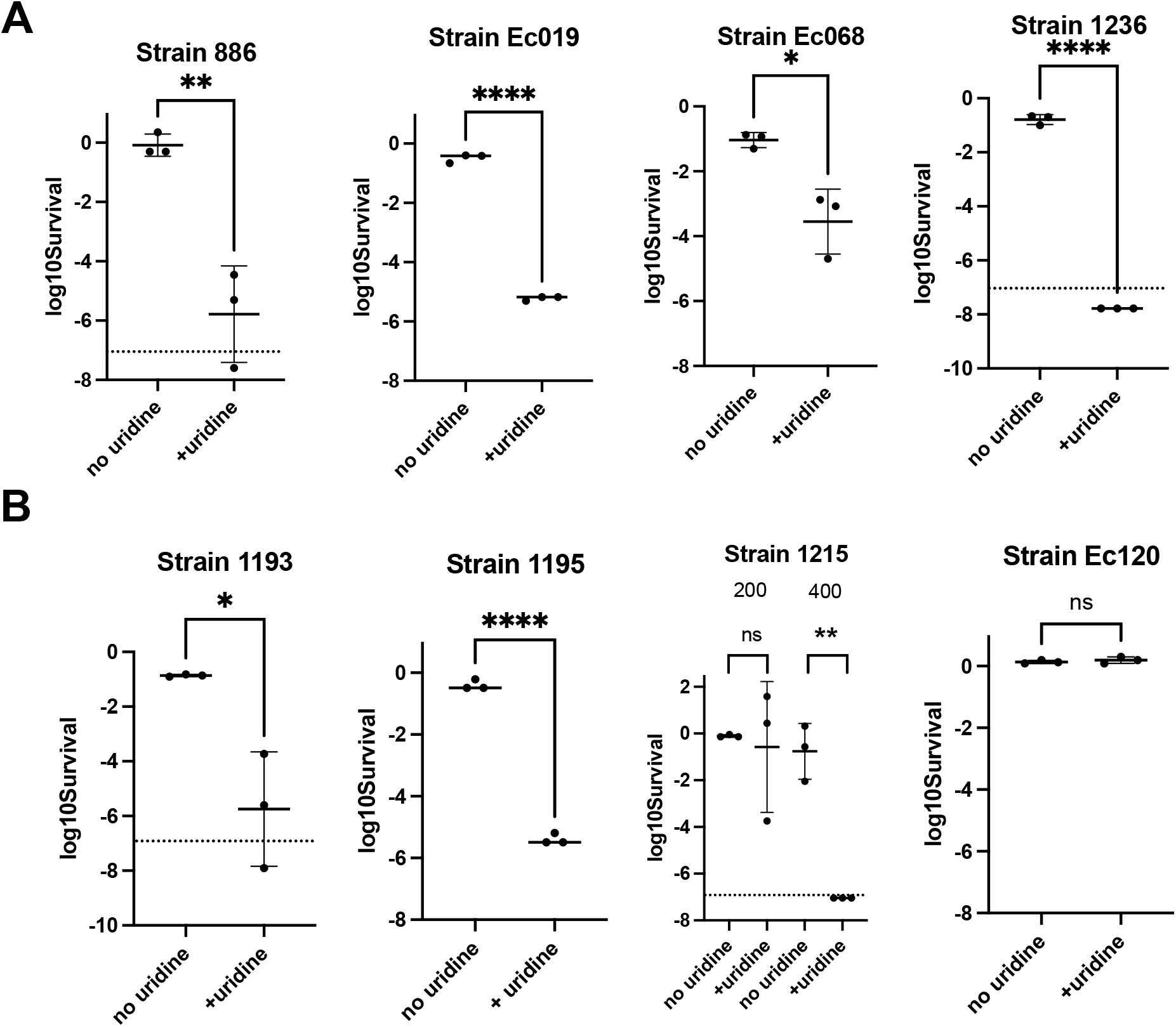
Uridine potentiates AGs on clinical *E. coli* strains in synthetic urine. **A.**Survival of susceptible *E. coli* strains after 20 hours of treatment with 10 µg/ml of tobramycin, supplemented or not with 0.031% of uridine. The dotted lines indicate the limit of detection (no colony on the pure culture plate, counted as 1 CFU). For statistical significance calculations, we used t-test. **** means p<0.0001, ** means p<0.01, * means p<0.05. Number of replicates for each experiment: n=3. **B.** Survival of resistant clinical *E. coli* strains after 20 hours of treatment with 200 µg/ml of tobramycin (Strains 1193 and 1195), 200 and 400 µg/ml of tobramycin (Strain 1215) or 400 µg/ml of tobramycin (Strain Ec120), supplemented or not with 0.031% of uridine. The dotted lines indicate the limit of detection (no colony on the pure culture plate, counted as 1 CFU). For statistical significance calculations, we used t-test (strains 1193, 1195 and Ec120) and one way-ANOVA (strain 1215). **** means p<0.0001, ** means p<0.01, * means p<0.05. ns: non-significant. Number of replicates for each experiment: n=3.

We also evaluated AG resistant clinical strains with AG modifying enzymes (Figure 4B). With tobramycin concentrations of 200 and 400 µg/ml, which are typically ineffective against these strains, the addition of uridine showed promising results. For strains 1193 and 1195, uridine supplementation reduced survival at 200 µg/ml, a concentration achievable in urine. In the case of strain 1215, the potentiating effect of uridine was observed at 400 µg/ml instead of 200 µg/ml. However, for strain Ec120, which carried both efflux pumps and AG modification enzymes, uridine did not demonstrate any effect, even at 400 µg/ml, potentially because of a very high level of resistance.

These findings indicate that uridine could be used in combination with AGs to effectively treat specific infections caused by resistant *E. coli* strains in clinics.

## Discussion

Until now, modulation of AG uptake into the Gram-negative bacterial cells was primarily associated with changes in proton motive force (PMF) ^9,15,33^, generated by bacterial respiration. While this remains true, our research has identified carbohydrate transporters as players of a new mechanism which can tune AG uptake in various Gram-negative bacteria. Previous reports have indicated that the efficiency of AG uptake can be influenced by carbon sources and metabolites, but this was proposed to be through modulation of PMF (e.g.: ^34–37^). Importantly, this means that sugars, which are capable of regulating the expression of their own transporters, can consequently regulate AG entry through upregulation of these transporters. Note that although AG entry through sugar transporters does not involve a change in the PMF, the presence of membrane potential is necessary, as respiration contributes to ATP production, and ATP powers transporters ^38^. Therefore, the requirement for the presence of PMF in AG uptake could be associated, at least partly, with the activity of the transporters.

The chemical structure of AGs consists of amino sugars linked to an aminocyclitol core structure ^10^. The sugar moiety potentially allows these molecules to be recognized as substrates by bacterial carbohydrate transporters. Consistently, overexpression or deletion of the CmtAB transporter does not impact susceptibility to spectinomycin, which lacks the sugar moiety. This is not the first report of an antibiotic molecule hijacking a bacterial transporter for uptake. For instance, in *P. aeruginosa*, imipenem which has structural similarities to amino acids was shown to be transported through the amino acid porin OprD2 ^39^.

AG resistance is usually associated either with genomic AG modifying factors which inactivate the AG molecules, or genetic mutations. These mutations can either impact the target of AGs (the ribosome), or more frequently, their uptake via decreased PMF ^27,40,41^. The reason why mutations in carbohydrate transporters have not been previously linked with AG resistance may be explained by the diverse array of transporters capable of transporting AGs. Through systematic overexpression of sugar transporters, we have identified at least 11 transport systems in *E. coli* that are specifically involved in AG uptake. These systems include phosphotransferase systems (PTS) for various sugars (chitobiose, glucitol, cellobiose/arbutine, fructose and fructose-like, β-glucoside and maltose), and the ABC transporter MalEFG for maltose. Such redundancy for AG transport is also consistent with the relatively mild phenotypic effects observed in single transporter deletions.

The individual impact of sugar transporters on AG susceptibility may appear modest when compared to genetic resistance mechanisms. However, simultaneous activation of multiple transporters can have a significant effect on sensitization to AGs. We showed that such upregulation can be achieved by supplementing uridine, the most potent activator of *cmtA* transcription among 198 substrates tested. Uridine potentiates the killing effect of the clinical AGs tobramycin, gentamicin, and amikacin in synthetic urine, a standardized medium that mimics the bacterial growth conditions in urinary tract infections (UTIs). This enhancement was due to an increased uptake of AGs by the bacteria.

Considering the well-established regulation of sugar transport by carbon catabolite control, we investigated the potential involvement of the CRP regulator in AG entry. As predicted ^42^, we show that CRP serves as a regulator of *cmtA* and contributes to AG susceptibility in *E. coli* (Figure S4). This does not exclude regulation of AG uptake through other transcription factors. Deletion of *crp* causes growth impairment even in laboratory conditions (Figure S4 and Figure S2A), we thus did not consider it for further study it in the context of urinary infections. Note that *Pseudomonadales* and enterobacteria like *E. coli* ^43^ show differences in catabolic repression and carbon source utilization. Nevertheless, the uptake of AGs through carbohydrate transporters appears to be a shared property among *Pseudomonadales* ^19^, and overexpression of carbohydrate transporters also enhances the killing of *P. aeruginosa* and *A. baumannii* by AGs. Therefore, improving treatments by exploiting carbohydrate/AGs transport represents a promising strategy to combat antibiotic resistance, considering the potential synergistic effects of AG and sugars.

In clinical settings, the emergence of AG resistance has been detected in 5.5% of the treated cases ^44^, and inadequate antibiotic dosing has been identified as a primary cause ^45^. Uridine supplementation, by accelerating the kinetics of the bacterial death, may be able to reduce the selection of AG resistance, as we observe in synthetic urine, and potentially limit the occurrence of recurrent infections. Co-treatment of uridine with AGs may be particularly beneficial in the case of UTIs such as cystitis or pyelonephritis but may also be applicable to otitis, or eye infections, where the administered doses can be easily adjusted due to the mode of drug delivery. For pulmonary infections, combining uridine with inhaled tobramycin, for example, could prove effective ^46^.

The World Health Organization has classified AGs as critically important antimicrobials for human medicine ^47^, underscoring the significance of seeking improvements in AG treatments ^48^. The use of uridine as adjuvant to AGs could be a potent approach to enhance treatment outcomes either by reducing the required AG dosage and mitigating the associated adverse effects, or by limiting the appearance of AG resistance and even re-sensitizing AG resistant bacteria. Treatment with high doses of uridine (up to 10 g/m²) induces no side effects ^30^. AGs exhibit a concentration-dependent killing ^49^, and are thus more effective in treating bacterial infections if higher doses are administered ^50^. Uridine offers a solution to enhance AG uptake, allowing for increased effective doses in bacteria without escalating toxicity for the patient.

## Acknowledgments

We thank Dominique Fourmy for the generous gift of the Neo-cy5. We thank Jean-Marc Ghigo’s lab for sharing equipments, Philippe Glaser for sharing *E. coli* clinical strains and helpful discussions. We are thankful to André Carvalho and Laurence Mulard for helpful discussions, and Evelyne Krin for technical advice. We thank Andreas Liaropoulos for his valuable help with urine experiments. This work was funded by Evotec ID Lyon, the Association Nationale pour la Recherche Technologique (Convention CIFRE n° 2019/0539), the Centre National de la Recherche Scientifique (CNRS-UMR 3525), the Fondation pour la Recherche Médicale (équipe FRM 202103012569), and Institut Pasteur Grant PTR 245-19.

## Authors Contributions

Conceptualization: M.L., S.R., D.M. and Z.B.

Methodology: M.L., S.R., I. E., M. I., E. D., and Z.B.

Experiments: M.L., A. A., T. B., M. I.

Investigation: M.L, S.R., and Z.B.

Supervision: S.R., E.B., D.M. and Z.B.

First draft: M.L.

Review and editing: M.L., S.R., Z.B, E.B. and D.M.

Fundings acquisition: Z.B., S.R. and D.M.

## Conflict of interest

The patent application EP23305045.9 covers the aspects of potentiation of aminoglycosides through activation of carbohydrate transporters described in the manuscript. This study is the result of a partnership between Institut Pasteur and Evotec ID Lyon. M.L., S. R. and E. B. were employed by Evotec ID Lyon during this study.

## Material and Methods

### Strains, plasmids, and primers

The strains, plasmids and primers used in the study are presented in Table S3.

### Growth conditions

MH medium was used to determine the MIC of the deletion or overexpression strains and for all the overnight cultures. MOPS Rich (Teknova EZ rich defined medium) was used for the screening assay to avoid intrinsic fluorescence of other media, and for the neocy5 assay. A medium containing 1% bactotryptone and 0.5% NaCl supplemented with the sugar was used for specific-substrate susceptibility tests. To mimic urinary tract infection, the urine synthetic medium was prepared according to ^29^. To mimic the human plasma, the HPLM medium (Human Plasma Like Medium, Thermofischer Scientific, Cat#A4899102) was used. The five species were grown at 37°C with shaking (150 rotations per minutes).

### Genes deletions

All engineered *E. coli* strains used in this work are derivatives of *E. coli* MG1655 and were constructed by transduction using Keio knockouts strains. Kanamycin resistance cassette *aph* was removed using FLP/FRT system ^51^. Regarding deletion of carbohydrate transporters, for multi-component PTS systems, we deleted the gene for the membrane-associated component.

### Overexpression of transporters

Overexpressions were performed by cloning the genes of interest into the pSEVA-238 vector ^52^ under the dependence of the *Pm* promoter, using 1 mM of sodium benzoate as inducer ^53^. The primers and restriction sites associated used for cloning are listed in Table S3.

### MIC evaluation

Etest: Overnight cultures in MH were diluted 20X in PBS (except for the Δ*crp* strain and in urine synthetic medium, culture were not diluted). Then, 300 µl were plated on the appropriate medium: MH for genes deletions; MH supplemented with kanamycin and sodium benzoate for maintenance and induction of the plasmid in overexpression strains; 1% amino acid, 0.5% NaCl and 0.5% of substrate (eg. Glucose, ribose, uridine …) to assess the impact of carbon sources, or in urine synthetic medium with agar. Plates were then dried for 10 minutes. Etests (Biomerieux) were placed on the plates and incubated overnight at 37°C, or until visible growth for urine synthetic medium.

Liquid cultures: MICs were determined by the microtiter broth dilution method with an initial inoculum size of 10^6^ CFUs/ml. In MH medium, the MIC was interpreted as the lowest antibiotic concentration preventing visible growth (used for neomycin). In urine synthetic medium, growth being difficult to interpret with naked eye, 5 µl of pure culture of each antibiotic dilution were plated and MIC was interpreted as the lowest concentration preventing growth.

### Neo-cy5 uptake assay

Quantification of fluorescent neomycin (Neo-cy5) uptake was performed as described ^23^. Neo-cy5 is an aminoglycoside (neomycin) coupled to the fluorophore Cy5 that retained activity and mode of uptake in Gram-negative bacteria ^24^. Overnight cultures were diluted 100X in rich MOPS (Teknova EZ rich defined medium). When the bacterial cultures reached an OD_600_ of 0.25, they were treated with 0.4 μM of Cy5 labeled neomycin for 15 minutes at 37°C under aluminum foil. For the assays with different substrates, cultures were washed once with PBS before treatment. 20 μl of each treated culture were then used for flow cytometry, diluted in 200 μl of PBS before reading fluorescence. Flow cytometry experiments were performed as described ^54^. For each experiment, 50000 events were counted on the Miltenyi MACSquant device with the Y3 laser.

### Evaluation of PMF

Quantification of PMF was performed using the Mitotracker Red CMXRos dye (Invitrogen)^55^, in parallel with the Neo-cy5 uptake assay, using the same bacterial cultures. 50 µl of each culture were mixed with 60 µl of PBS. Tetrachlorosalicylanilide TCS (Thermofischer), a protonophore, was used as a negative control with a 500 µM treatment applied for 10 minutes at room temperature. Then, 25 nM of Mitotracker Red were added to each sample and let at room temperature for 15 minutes under aluminium foil. 20 μl of the treated culture were then used for flow cytometry, diluted in 200 μl of PBS before reading fluorescence. Flow cytometry was performed as described above using the Y2 laser.

### Carbon sources screening assay

The GFPmut3 was fused to promoter of interest ^56^ and cloned into a plasmid pSC101. For the first screening, overnight cultures of the strain carrying the screening system were diluted 200X in MOPS Rich (Teknova EZ rich defined medium) supplemented with carbenicillin for plasmid maintenance. The phenotype Microarray (Biolog) plates PM1 and PM2B (carbon sources), and PM3B, for a total of 198 molecules including 7 nucleosides and 6 nucleotides were used for molecules screening. Each well was filled with 100 µl of inoculated media and mixed by pipetting. Media were transferred to 96 well dark-bottom plates (Thermo Scientific). GFP fluorescence was followed on the Tecan Infinite 200 PRO (Life Science) at 37°C during 8 hours. Fluorescence induction by the substrate was calculated using the ratio fluorescence (t8h-t0h) over growth (t8h-t0h OD_600nm_).

For flow cytometry quantification, overnight cultures in MH of strain carrying the screening system were diluted 200X in rich MOPS (Teknova EZ rich defined medium) supplemented with carbenicillin for plasmid maintenance and grown overnight, and the molecule tested at 0.5% (except uracil: 0.1%, limit of solubility). Fluorescence was read on 5 µl of overnight cultures diluted in 200 µl of PBS with the B1 laser.

### Stringent response

The P1*rrnB*-GFP fusion was constructed using GFP ASV, and cloned into a plasmid pSC101 ^57^. Cultures were grown overnight (positive control) or diluted 100X until reaching an OD_600nm_ of 0.4, in MH medium supplemented with carbenicillin for plasmid maintenance and 0.5% of the mentioned substrate or 0.1 µg/ml of tobramycin. Fluorescence was read by flow cytometry, on 20 µl of cultures diluted in 200 µl of PBS.

### Killing assay

On bactotryptone medium: Overnight cultures were diluted 1000X in 25 ml of medium containing 1% bactotryptone and 0.5% NaCl, supplemented or not with glucose, maltose or uridine 0.5%. Cultures were grown to an OD_600nm_ of 0.3-0.4, and a 5 ml aliquot was treated with a lethal concentration of tobramycin (10 µg/ml). Cultures were plated at 0 h (t0), 1, 2, 4, 6 and 20 hours after treatment, and survival was calculated by counting CFUs/ml after treatment divided by the initial number of CFUs/ml (t0).

On synthetic urine medium: Overnight cultures in MH were diluted 10-fold in PBS. Approximately 10^7^ CFUs/ml were diluted in 200 µl of urine synthetic medium containing or not uridine and treated with AGs, in a 96-well plate, then incubated at 37°C with 100 rpm. Cultures were plated at 0 h (t0), 1, 2, 4, 6 and 24 hours after treatment, and survival was calculated by counting CFUs/ml after treatment divided by the initial number of CFUs/ml (t0).

### Dose-response assay

Two-fold dilutions of uridine in urine synthetic medium were prepared into column of a 96 wells plate, from 2% to 0.0018%. The first column was only filled with synthetic urine medium without uridine. Then, 100 µl of a solution containing 2X concentration of the tested antibiotic was added, thus diluted uridine and antibiotics by two and increasing the total volume per well at 200 µl. The last line was only filled with urine synthetic medium without antibiotic. Each well was then inoculated with approximately 10^7^ CFUs from MH stationary phase culture and incubated 20 hours at 37°C with agitation (110 rpm). CFUs were count by plating before and after treatment on MH medium.

### Whole genome sequencing

gDNA was extracted from 500 µl of overnight cultures in MH, using the Blood and Tissue Extraction Kit (Qiagen) according to the manufacturer instructions. The presence of variants (Single Nucleotide Polymorphism) was analyzed with SnpEff 5.0 ^58^.

### Statistical analysis

Means and standard deviations for growth curves and survival rate, means and geometric means for logarithmic values were calculated using GraphPad Prism. All analysis were realized using GraphPad Prism. F-test was first performed in order to determine whether the variances were equal or different between conditions. For conditions with significantly different variances, Welch correction was applied. For two groups comparison, Student’s t-test was used. One-way ANOVA or two-way ANOVA were used for multiple comparisons. ∗∗∗∗ means p<0.0001, ∗∗∗ means p<0.001, ∗∗ means p<0.01, ∗ means p<0.05. Number of replicates for each experiment was 3<n<7.

## References

1. Clatworthy, A. E., Pierson, E. & Hung, D. T. Targeting virulence: a new paradigm for antimicrobial therapy. Nat. Chem. Biol. 3, 8 (2007).

2. Cassini, A. et al. Attributable deaths and disability-adjusted life-years caused by infections with antibiotic-resistant bacteria in the EU and the European Economic Area in 2015: a population-level modelling analysis. Lancet Infect. Dis. 19, 56–66 (2019).

3. Miller, S. I. Antibiotic Resistance and Regulation of the Gram-Negative Bacterial Outer Membrane Barrier by Host Innate Immune Molecules. mBio 7, e01541–16 (2016).

4. Murray, C. J. et al. Global burden of bacterial antimicrobial resistance in 2019: a systematic analysis. The Lancet 399, 629–655 (2022).

5. Avent, M. L., Rogers, B. A., Cheng, A. C. & Paterson, D. L. Current use of aminoglycosides: indications, pharmacokinetics and monitoring for toxicity: Aminoglycosides: review and monitoring. Intern. Med. J. 41, 441–449 (2011).

6. Selimoglu, E. Aminoglycoside-induced ototoxicity. Curr. Pharm. Des. 13, 119–126 (2007).

7. Martínez-Salgado, C., López-Hernández, F. J. & López-Novoa, J. M. Glomerular nephrotoxicity of aminoglycosides. Toxicol. Appl. Pharmacol. 223, 86–98 (2007).

8. Lopez-Novoa, J. M., Quiros, Y., Vicente, L., Morales, A. I. & Lopez-Hernandez, F. J. New insights into the mechanism of aminoglycoside nephrotoxicity: an integrative point of view. Kidney Int. 79, 33– 45 (2011).

9. Taber, H. W., Mueller, J. P., Miller, P. F. & Arrow, A. S. Bacterial uptake of aminoglycoside antibiotics. Microbiol. Rev. 51, 439–457 (1987).

10. Krause, K. M., Serio, A. W., Kane, T. R. & Connolly, L. E. Aminoglycosides: An Overview. Cold Spring Harb. Perspect. Med. 6, a027029 (2016).

11. Martin, N. L. & Beveridge, T. J. Gentamicin interaction with Pseudomonas aeruginosa cell envelope. Antimicrob. Agents Chemother. 29, 1079–1087 (1986).

12. Bryan, L. E. & Van DenElzen, H. M. Streptomycin Accumulation in Susceptible and Resistant Strains of *Escherichia coli* and *Pseudomonas aeruginosa*. Antimicrob. Agents Chemother. 9, 928–938 (1976).

13. Vaara, M. Agents That Increase the Permeability of the Outer Membrane. MICROBIOL REV 56, 17 (1992).

14. Bafna, J. A. et al. Kanamycin Uptake into *Escherichia coli* Is Facilitated by OmpF and OmpC Porin Channels Located in the Outer Membrane. ACS Infect. Dis. 6, 1855–1865 (2020).

15. Fraimow, H. S., Greenman, J. B., Leviton, I. M., Dougherty, T. J. & Miller, M. H. Tobramycin uptake in Escherichia coli is driven by either electrical potential or ATP. J. Bacteriol. 173, 2800–2808 (1991).

16. Muir, M. E., Van Heeswyck, R. S. & Wallace, B. J. Effect of Growth Rate on Streptomycin Accumulation by Escherichia coli and Bacillus megaterium. Microbiology 130, 2015–2022 (1984).

17. Davies, J., Gilbert, W. & Gorini, L. Streptomycin, suppression, and the code. Proc. Natl. Acad. Sci. U. S. A. 51, 883–890 (1964).

18. Davis, B. D., Chen, L. L. & Tai, P. C. Misread protein creates membrane channels: an essential step in the bactericidal action of aminoglycosides. Proc. Natl. Acad. Sci. 83, 6164–6168 (1986).

19. Pierlé, S. A. et al. Identification of the active mechanism of aminoglycoside entry in V. cholerae through characterization of sRNA ctrR, regulating carbohydrate utilization and transport. Preprint: https://doi.org/10.1101/2023.07.19.549712 (2023).

20. Luo, X., Esberard, M., Bouloc, P. & Jacq, A. A Small Regulatory RNA Generated from the malK 5’ Untranslated Region Targets Gluconeogenesis in Vibrio Species. mSphere e0013421 (2021) doi:10.1128/mSphere.00134-21.

21. Sprenger, G. A. Two open reading frames adjacent to the Escherichia coli K-12 transketolase (tkt) gene show high similarity to the mannitol phosphotransferase system enzymes from Escherichia coli and various Gram-positive bacteria. Biochim. Biophys. Acta BBA - Gen. Subj. 1158, 103–106 (1993).

22. Grisafi, P. L. et al. Deletion mutants of the Escherichia coli K-12 mannitol permease: dissection of transport-phosphorylation, phospho-exchange, and mannitol-binding activities. J. Bacteriol. 171, 2719–2727 (1989).

23. Lang, M. et al. Sleeping ribosomes: Bacterial signaling triggers RaiA mediated persistence to aminoglycosides. iScience 24, 103128 (2021).

24. Sabeti Azad, M., et al. Fluorescent Aminoglycoside Antibiotics and Methods for Accurately Monitoring Uptake by Bacteria. ACS Infect. Dis. 6, 1008–1017 (2020).

25. Brückner, R. & Titgemeyer, F. Carbon catabolite repression in bacteria: choice of the carbon source and autoregulatory limitation of sugar utilization. FEMS Microbiol. Lett. 209, 141–148 (2002).

26. Görke, B. & Stülke, J. Carbon catabolite repression in bacteria: many ways to make the most out of nutrients. Nat. Rev. Microbiol. 6, 613–624 (2008).

27. Bolard, A., Plésiat, P. & Jeannot, K. Mutations in Gene fusA1 as a Novel Mechanism of Aminoglycoside Resistance in Clinical Strains of Pseudomonas aeruginosa. Antimicrob. Agents Chemother. 62, e01835–17 (2018).

28. Maisnier-Patin, S., Paulander, W., Pennhag, A. & Andersson, D. I. Compensatory evolution reveals functional interactions between ribosomal proteins S12, L14 and L19. J. Mol. Biol. 366, 207–215 (2007).

29. Sarigul, N., Korkmaz, F. & Kurultak, İ. A New Artificial Urine Protocol to Better Imitate Human Urine. Sci. Rep. 9, 20159 (2019).

30. Albert Leyva et al. Phase I and Pharmacokinetic Studies of High-Dose Uridine Intended for Rescue from 5-Fluorouracil Toxicity. Cancer Res. 44, 5928–5933 (1984).

31. Wood, M. J. & Farrell, W. Comparison of Urinary Excretion of Tobramycin and Gentamicin in Adults. J. Infect. Dis. 134, S133–S138 (1976).

32. Edgar, R. & Bibi, E. MdfA, an Escherichia coli multidrug resistance protein with an extraordinarily broad spectrum of drug recognition. J. Bacteriol. 179, 2274–2280 (1997).

33. Webster, C. M. et al. Proton motive force underpins respiration-mediated potentiation of aminoglycoside lethality in pathogenic Escherichia coli. Arch. Microbiol. 204, 120 (2022).

34. Martins, D. & Nguyen, D. Stimulating Central Carbon Metabolism to Re-sensitize Pseudomonas aeruginosa to Aminoglycosides. Cell Chem. Biol. 24, 122–124 (2017).

35. Crabbé, A. et al. Host metabolites stimulate the bacterial proton motive force to enhance the activity of aminoglycoside antibiotics. PLOS Pathog. 15, e1007697 (2019).

36. Allison, K. R., Brynildsen, M. P. & Collins, J. J. Metabolite-enabled eradication of bacterial persisters by aminoglycosides. Nature 473, 216–220 (2011).

37. Rosenberg, C. R., Fang, X. & Allison, K. R. Potentiating aminoglycoside antibiotics to reduce their toxic side effects. PLoS ONE 15, e0237948 (2020).

38. Schneider, E. ABC transporters catalyzing carbohydrate uptake. Res. Microbiol. 152, 303–310 (2001).

39. Trias, J. & Nikaido, H. Protein D2 channel of the Pseudomonas aeruginosa outer membrane has a binding site for basic amino acids and peptides. J. Biol. Chem. 265, 15680–15684 (1990).

40. Thorbjarnardóttir, S. H., Magnúsdóttir, R. Á., Eggertsson, G., Kagan, S. A. & Andrésson, Ó. S. Mutations determining generalized resistance to aminoglycoside antibiotics in Escherichia coli. Mol. Gen. Genet. MGG 161, 89–98 (1978).

41. Hancock, R. E. Aminoglycoside uptake and mode of action--with special reference to streptomycin and gentamicin. I. Antagonists and mutants. J. Antimicrob. Chemother. 8, 249–276 (1981).

42. Shimada, T., Fujita, N., Yamamoto, K. & Ishihama, A. Novel Roles of cAMP Receptor Protein (CRP) in Regulation of Transport and Metabolism of Carbon Sources. PLoS ONE 6, e20081 (2011).

43. Sonnleitner, E., Abdou, L. & Haas, D. Small RNA as global regulator of carbon catabolite repression in Pseudomonas aeruginosa. Proc Natl Acad Sci USA 106, 21866–21871 (2009).

44. Fish, D. N., Piscitelli, S. C. & Danziger, L. H. Development of Resistance During Antimicrobial Therapy: A Review of Antibiotic Classes and Patient Characteristics in 173 Studies. Pharmacother. J. Hum. Pharmacol. Drug Ther. 15, 279–291 (1995).

45. Abdelraouf, K., Linder, K. E., Nailor, M. D. & Nicolau, D. P. Predicting and preventing antimicrobial resistance utilizing pharmacodynamics: part II Gram-negative bacteria. Expert Opin. Drug Metab. Toxicol. 13, 705–714 (2017).

46. Smith, S., Rowbotham, N. J. & Regan, K. H. Inhaled anti-pseudomonal antibiotics for long-term therapy in cystic fibrosis. Cochrane Database Syst. Rev. (2018) doi:10.1002/14651858.CD001021.pub3.

47. World Health Organization. Critically important antimicrobials for human medicine: 6th revision. https://www.who.int/publications-detail-redirect/9789241515528 (2018).

48. Serio, A. W., Keepers, T., Andrews, L. & Krause, K. M. Aminoglycoside Revival: Review of a Historically Important Class of Antimicrobials Undergoing Rejuvenation. EcoSal Plus 8, (2018).

49. Reed, M. D. Optimal antibiotic dosing. The pharmacokinetic-pharmacodynamic interface. Postgrad. Med. 108, 17–24 (2000).

50. Kashuba, A. D., Nafziger, A. N., Drusano, G. L. & Bertino, J. S. Optimizing aminoglycoside therapy for nosocomial pneumonia caused by gram-negative bacteria. Antimicrob. Agents Chemother. 43, 623–629 (1999).

51. Zhu, X.-D. & Sadowski, P. D. Cleavage-dependent Ligation by the FLP Recombinase: Characterization of a mutant Flp protein with an alteration in a a catalytic amino acid. J. Biol. Chem. 270, 23044–23054 (1995).

52. Silva-Rocha, R. et al. The Standard European Vector Architecture (SEVA): a coherent platform for the analysis and deployment of complex prokaryotic phenotypes. Nucleic Acids Res. 41, D666–D675 (2013).

53. Kessler, B., Timmis, K. N. & de Lorenzo, V. The organization of the Pm promoter of the TOL plasmid reflects the structure of its cognate activator protein XylS. Mol. Gen. Genet. MGG 244, 596– 605 (1994).

54. Baharoglu, Z., Bikard, D. & Mazel, D. Conjugative DNA Transfer Induces the Bacterial SOS Response and Promotes Antibiotic Resistance Development through Integron Activation. PLoS Genet. 6, e1001165 (2010).

55. El Mortaji, L., et al. A peptide of a type I toxin−antitoxin system induces *Helicobacter pylori* morphological transformation from spiral shape to coccoids. Proc. Natl. Acad. Sci. 117, 31398–31409 (2020).

56. Cormack, B. P., Valdivia, R. H. & Falkow, S. FACS-optimized mutants of the green fluorescent protein (GFP). Gene 173, 33–38 (1996).

57. Babosan, A. et al. Nonessential tRNA and rRNA modifications impact the bacterial response to sub-MIC antibiotic stress. microLife 3, 18 (2022).

58. Cingolani, P. et al. A program for annotating and predicting the effects of single nucleotide polymorphisms, SnpEff: SNPs in the genome of Drosophila melanogaster strain w ^1118^; iso-2; iso-3. Fly (Austin) 6, 80–92 (2012).

59. Castañeda-García, A., Rodríguez-Rojas, A., Guelfo, J. R. & Blázquez, J. The glycerol-3-phosphate permease GlpT is the only fosfomycin transporter in Pseudomonas aeruginosa. J. Bacteriol. 191, 6968– 6974 (2009).

60. Miller, M. H., Edberg, S. C., Mandel, L. J., Behar, C. F. & Steigbigel, N. H. Gentamicin Uptake in Wild-Type and Aminoglycoside-Resistant Small-Colony Mutants of *Staphylococcus aureus*. Antimicrob. Agents Chemother. 18, 722–729 (1980).

61. Patching, S. G. et al. The nucleoside transport proteins, NupC and NupG, from Escherichia coli: specific structural motifs necessary for the binding of ligands. Org. Biomol. Chem. 3, 462 (2005).

62. Sakai, T. T. & Cohen, S. S. Interrelation Between Guanosine Tetraphosphate Accumulation, Ribonucleic Acid Synthesis, and Streptomycin Lethality in Escherichia coli CP78. Antimicrob. Agents Chemother. 7, 730–735 (1975).

63. Koskiniemi, S., Pränting, M., Gullberg, E., Näsvall, J. & Andersson, D. I. Activation of cryptic aminoglycoside resistance in Salmonella enterica: Activation of cryptic resistance. Mol. Microbiol. 80, 1464–1478 (2011).

64. Kim, N. et al. DksA Modulates Antimicrobial Susceptibility of *Acinetobacter baumannii*. Antibiot. Basel Switz. 10, (2021).

65. Park, Y.-H., Lee, C.-R., Choe, M. & Seok, Y.-J. HPr antagonizes the anti-σ70 activity of Rsd in Escherichia coli. Proc. Natl. Acad. Sci. U. S. A. 110, 21142–21147 (2013).

66. Carvalho, A., Krin, E., Korlowski, C., Mazel, D. & Baharoglu, Z. Interplay between Sublethal Aminoglycosides and Quorum Sensing: Consequences on Survival in V. cholerae. Cells 10, 3227 (2021).

67. Fruchard, L. et al. Queuosine modification of tRNA-Tyrosine elicits translational reprogramming and enhances growth of Vibrio cholerae with aminoglycosides. Preprint: https://doi.org/10.1101/2022.09.26.509455 (2022)

68. Wu, K.-M. et al. Genome Sequencing and Comparative Analysis of Klebsiella pneumoniae NTUH-K2044, a Strain Causing Liver Abscess and Meningitis. J. Bacteriol. 191, 4492–4501 (2009).

