## Supplemental figures and tables for "Uridine as a potentiator of aminoglycosides through activation of carbohydrate transporters"

**Supplementary figures:**

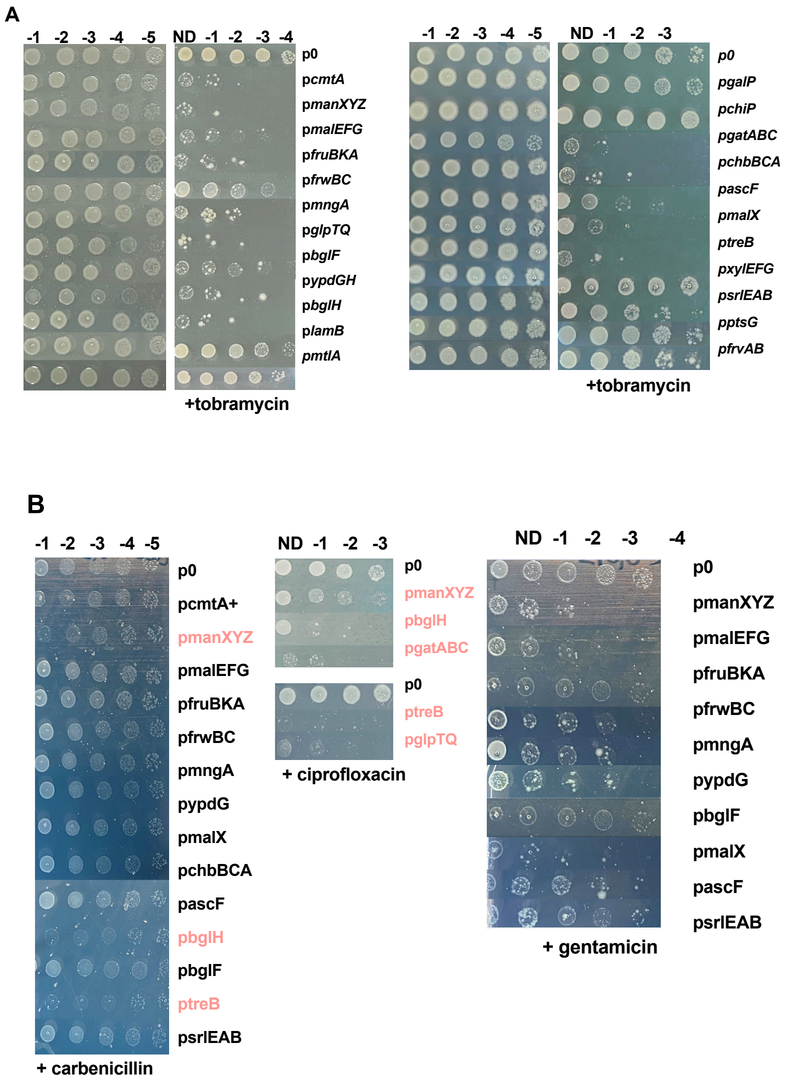

**Figure S1: Overexpression of 11 carbohydrates transporters sensitize to AGs specifically.** See also Table S1. **A.** Response to tobramycin (0.1 µg/ml), carbenicillin (3 µg/ml), ciprofloxacin (0.05 µg/ml) or gentamicin (0.08 µg/ml) of *E. coli* carrying the empty plasmid (p0) compared to the vector overexpressing carbohydrates transporters. Cultures of each strain were grown overnight. Susceptibility was assessed by serial dilutions (ND: non-diluted) and dropping 5 µl of each dilution on plates containing or not 0.1 µg/ml of tobramycin. Inducer was added in the media. Overexpression of 16 transporters sensitize to tobramycin. The *frwBC*+ strain displayed more pronounced growth impairment in media containing tobramycin than the control with an empty plasmid, but did not exhibit a lower MIC to tobramycin compared to the empty plasmid. Overexpression of the LamB porin at the outer membrane showed no response to AGs, nor did ChiP (chitoporin), PtsG (glucose PTS), XylEFG (Major Facilitator Superfamily (MFS) protein of xylose), GalP (MFS protein of galactose), FrvAB (fructose-like PTS) or MtlA (Mannitol PTS)**.** **B**. Overexpression of 5 transporters sensitize also to carbenicillin or ciprofloxacin. Strains overexpressing *manXYZ (*mannose PTS*)*, *treB* (trehalose PTS*)*, *bglH* (β-glucoside porin), *gatABC* (galacticol PTS) and *glpTQ* (glycerol-3-phosphate permease already known to favor uptake of fluoroquinolone[59]) showed a phenotype of susceptibility to AGs , but also to ciprofloxacin or carbenicillin. Overexpression of 11 transporters sensitize also to gentamicin (AG).

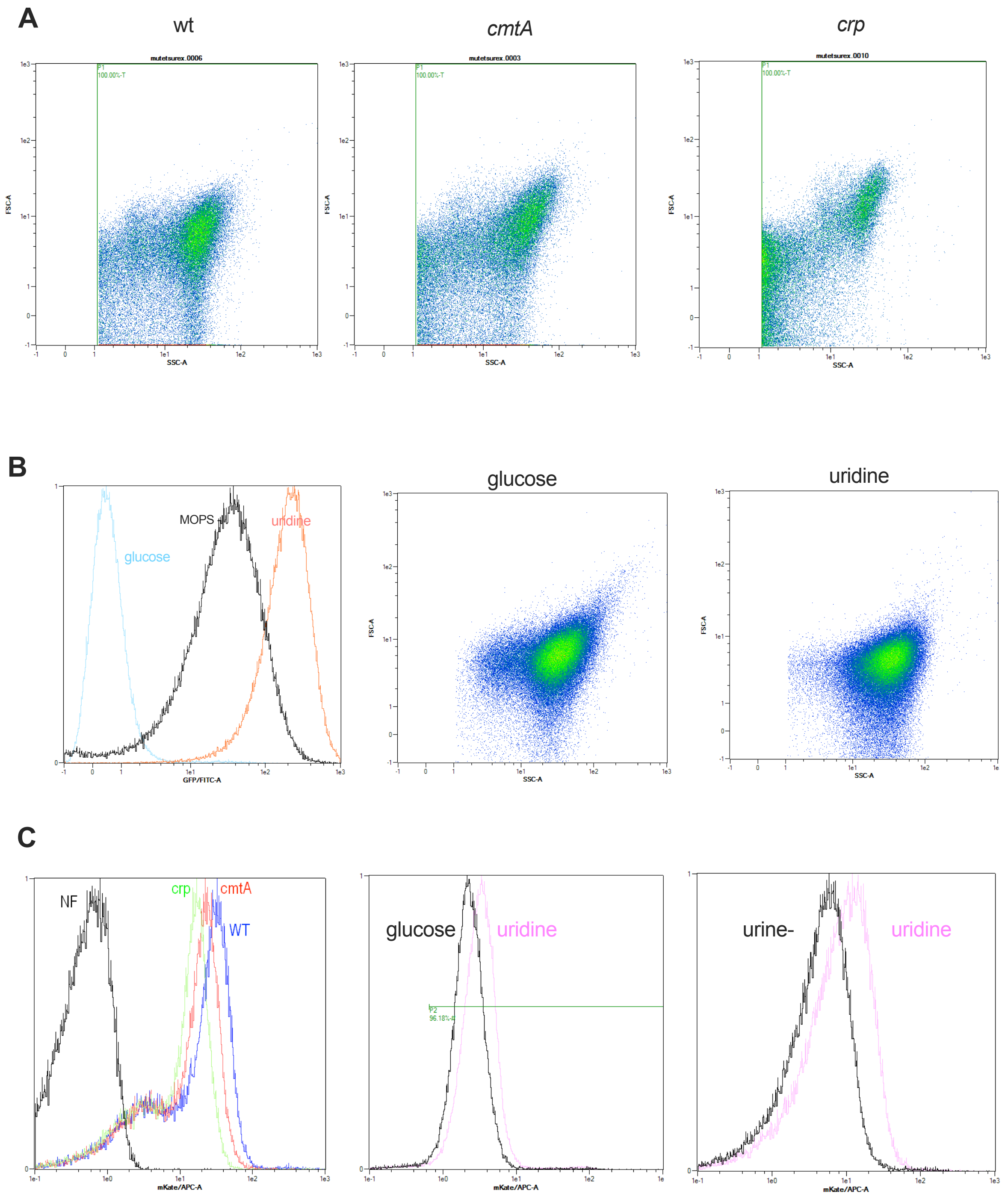

**Figure S2:** Flow cytometry data. **A.** Forward Scatter and Side Scatter (size and granularity) of *E. coli* WT, *cmtA* and *crp*. **B**. Forward Scatter and Side Scatter (size and granularity) of *E. coli* growing in MOPS Rich supplemented with glucose or uridine. Associated fluorescence (laser B2) of P*cmtA*-GFP with glucose, uridine or no supplementation. **C.** Fluorescence of Neocy5 detected by the Y3 laser in the mutant and with uridine compared to glucose or in synthetic urine.

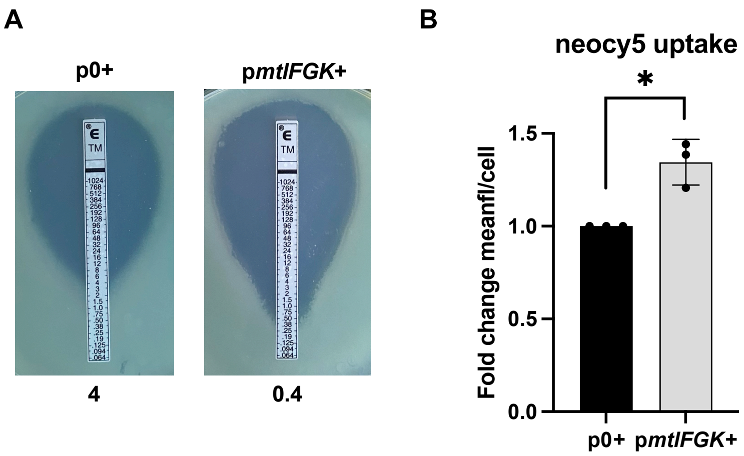

**Figure S3: Overexpression of the carbohydrate transporter *mtlFGK* in *P. aeruginosa* sensitizes to tobramycin by increasing the uptake**. **A**. Etest on *P. aeruginosa* carrying the empty vector (p0+) compared to the vector overexpressing MtlFGK (p*mtlFGK*+). Inducer was added on the medium. **B.** Neocy5 uptake evaluated by flow cytometry on *P. aeruginosa* carrying a plasmid overexpressing MtlFGK (p*mtlFGK*+) compared to the strain carrying the empty vector (p0+), expressed as fold change of mean fluorescence per cell. For statistical significance calculations, we used one-way t-test. * means $p<0.05$. Number of replicates n=3.

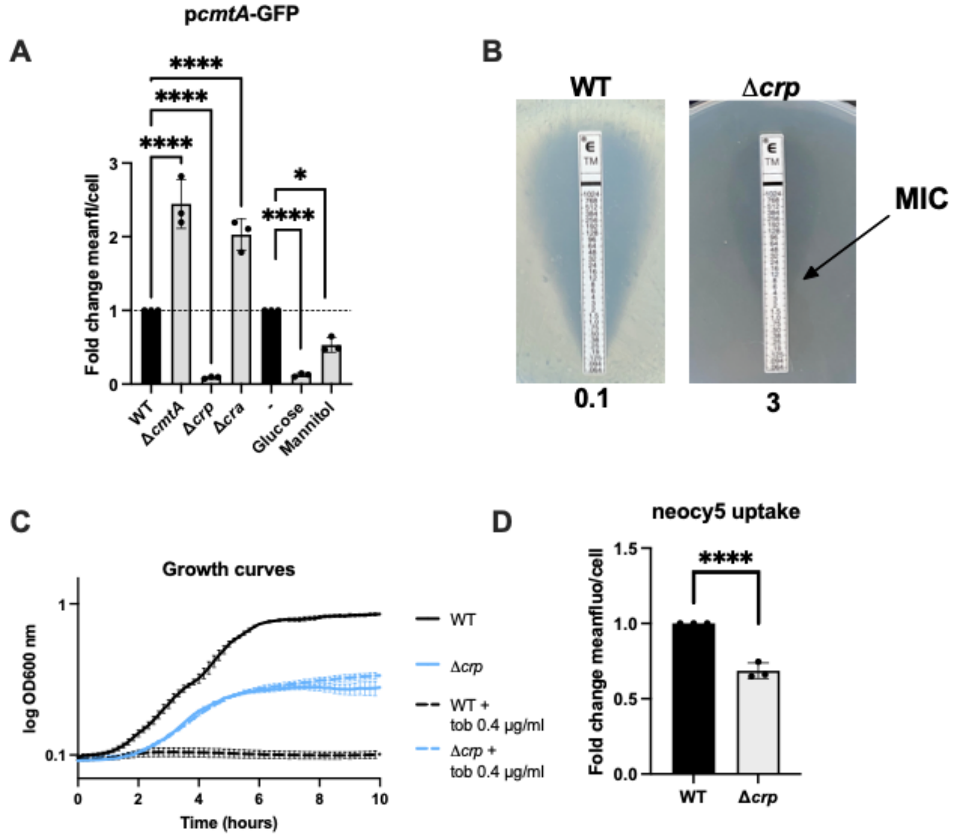

**Figure S4. CRP is involved in regulation of *cmtAB* and AG uptake**. CRP (cyclic AMP Repressor Protein), is the primary regulator of carbon catabolic repression and transcriptional activator of non-preferential sugar transporters. **A.** Quantification of GFP fluorescence using flow cytometry depending on the strain (WT Δ*cmtA*, Δ*crp* or Δ*cra*) or the molecule added in the medium (supplemented or not with glucose or mannitol 0.5%), expressed as fold change of mean fluorescence per cell (compared either to the WT for mutants, or to the no substrate condition for carbon sources). For statistical significance calculations, we used one-way ANOVA. **** means p<0.0001, * means p<0.05. Number of replicates for each experiment: n=3. *cmtAB* is controlled by CRP and its expression varies according to supplemented sugar**. B.** MIC of tobramycin, indicated in µg/ml, measured using Etests in *E. coli* WT and *Δcrp*. The Δ*crp* strain exhibited a 10-fold higher MIC compared to the WT strain, despite significant growth impairment. **C.** Growth curve of *E. coli* WT and Δ*crp*, in presence or not of tobramycin 0.4 µg/ml. Δ*crp* displayed no susceptibility to a 0.4 µg/ml tobramycin treatment, but CRP deletion is detrimental for the strain. Based on this, we hypothesized that transporters involved in AG uptake are non-preferential carbohydrates transporters repressed by glucose through CRP/cAMP regulation. **D**. Uptake of Neo-cy5 evaluated by flow cytometry on *E. coli* Δ*crp* compared to the WT strain, expressed as fold change of mean fluorescence per cell (compared either to the WT for mutants or to the empty vector for overexpression). For statistical significance calculations, we used one-way ANOVA (same experiment as Figure 1D). **** means p<0.0001. Number of replicates for each experiment: n=3. Neo-cy5 uptake is decreased in the *Δcrp* mutant, which aligns with the higher MIC value observed. To determine whether the potentiating effect of uridine was dependent on CRP, we tested the MIC in the presence of glucose or uridine in the Δ*crp* mutant strain. The MIC of the Δ*crp* increased to 6 µg/ml in the presence of glucose and remained unchanged in the presence of uridine. These results support the hypothesis that the increased expression of carbohydrate transporters stimulated by uridine is CRP-dependent (Table S1).

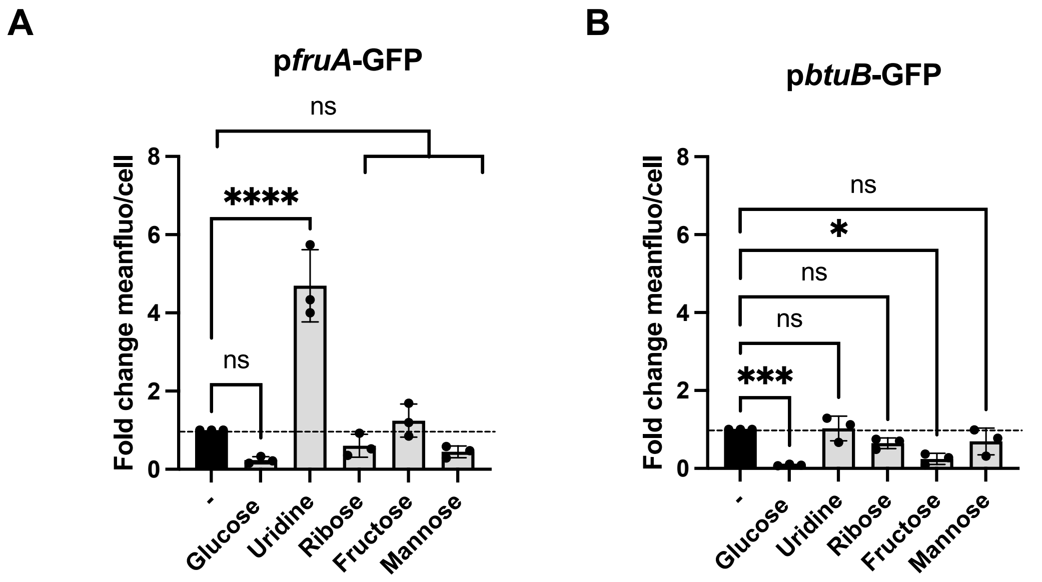

**Figure S5: GFP expression from *fruA* or *btuB* promoter depending on the molecule. A.** Quantification of GFP fluorescence from the *fruA* promoter using flow cytometry depending on the molecule added in the media (supplemented or not with glucose, uridine, ribose, fructose or mannose 0.5%) expressed as fold change of mean fluorescence per cell. **B.** Quantification of GFP fluorescence from the *btuB* promoter using flow cytometry depending on the molecule added in the medium (supplemented or not with glucose, uridine, ribose, fructose or mannose 0.5%) expressed as fold change of mean fluorescence per cell. For statistical significance calculations, we used one-way ANOVA. **** means p<0.0001, *** means p<0.001, * means p<0.05, ns means not significant. Number of replicates for each experiment: n=3.

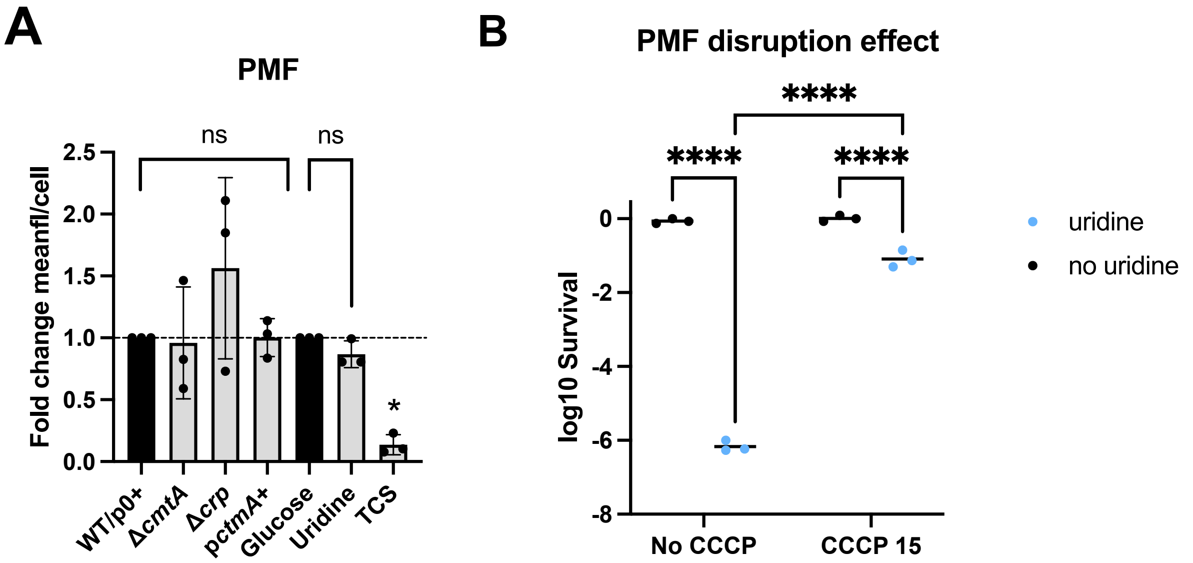

**Figure S6: Uridine effect is PMF-dependent, with no effect on PMF changes. A.** Previous studies have indicated a connection between changes in membrane potential and alterations in AG resistance [60]. To investigate whether differences in AG uptake are associated with variations in the PMF, we employed Mitotracker Red [23,55], a probe that accumulates based on membrane potential, in the WT, Δ*cmtA*, Δ*crp*, and p0/p*cmtA*+ strains. No changes in PMF were observed in the deletion strains or in the *cmtAB* overexpression strain, indicating that carbohydrate transporters facilitate AG uptake without affecting the PMF. Next, we investigated whether uridine supplementation influences the PMF. The addition of uridine or glucose to the medium did not result in any alteration of the PMF (Figure S5A), despite a 10-fold MIC difference. Therefore, the positive effect of uridine on AG uptake cannot be attributed to an increase in membrane potential.

On the other hand, the absence of an increase in PMF resulting from uridine supplementation does not rule out the possibility of a requirement for PMF in the effect of uridine on AG uptake. To address this question, we investigated the impact of uridine addition on the MIC of tobramycin in the presence or absence of carbonyl cyanide m-chlorophenyl hydrazine (CCCP), a protonophore which dissipates the ΔΨ. The addition of 15 µg/ml of CCCP substantially diminished the potentiating effect of uridine on AGs when the cells when treated with 4 µg/ml of tobramycin (Figure S5B). These results indicate that the effect of uridine on AG uptake relies on the presence of the PMF, likely because the activity of sugar transporters is dependent on the functionality of ATP synthesis. In **B.** Uptake of Mitotracker Red evaluated by flow cytometry on *E. coli* Δ*cmtA* and Δ*crp* compared to the WT strain ; *E. coli* carrying a plasmid overexpressing CmtA compared to the strain carrying the empty vector (p0); and *E. coli* WT grown either with a supplementation of glucose or uridine 0.5%, expressed as fold change of mean fluorescence per cell. TCS (tetrachlorosalicylanilide) treatment was used as negative control. For statistical significance calculations, we used one-way ANOVA. * means $p<0.05$, ns: non-significant, compared to WT/p0+ or Glucose. Number of replicates for each experiment: n=3. **C.** Survival to tobramycin 4 µg/ml treatment in urine synthetic media supplemented or not with uridine 0.5%, in the presence or not of 15 µM of the protonophore carbonyl cyanide m-chlorophenyl hydrazine (CCCP). Survival was assessed by plating and counting CFU after 16 hours of treatment. For statistical significance calculations, we used two-way ANOVA. **** means $p<0.0001$. Number of replicates for each experiment: n=3.

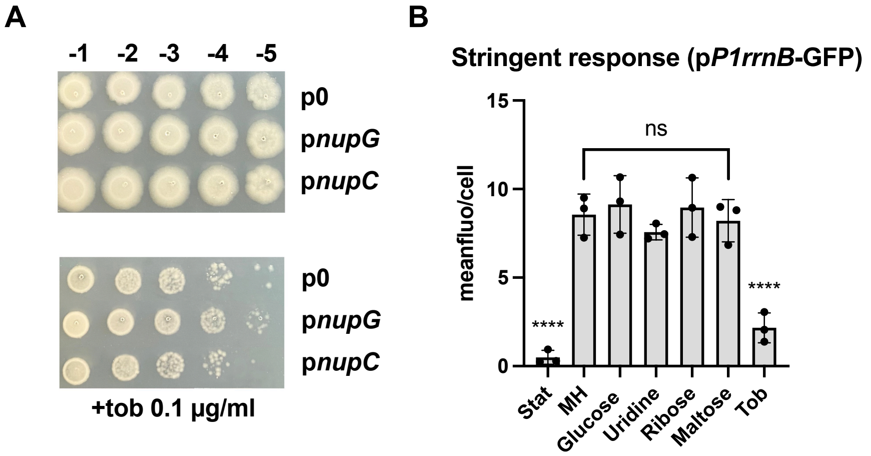

**Figure S7: Uridine mediated AG susceptibility is not related to uridine uptake, catabolism, PMF increasing or stress responses.** We have demonstrated that uridine induces the expression of at least two carbohydrate transporters, namely *cmtA* and *fruA*, resulting in enhanced uptake of AGs and increased susceptibility. We further explored the involvement of nucleoside transporters in AG uptake. Specifically, we overexpressed nup*G* and *nupC,* which are known to transport uridine and other nucleosides [61]. None of these overexpressions had any impact on the susceptibility to tobramycin (Table S1). This indicates that AGs cannot enter cells through nucleoside transporters. Furthermore, when uridine was added to the medium along with overexpressed nucleoside transporters, leading to increased intracellular uridine levels, we observed no increase in susceptibility to tobramycin **(A)**. These findings suggest that the level of uridine uptake is not responsible for the heightened susceptibility to tobramycin. However, a basal level of uridine uptake may still be necessary to increase the expression of the sugar transporters. To determine whether the catabolism of uridine is necessary for the potentiation of AGs, we deleted *udk* (responsible for the degradation of uridine into uracil and ribose-1-phosphate) and *udp* (involved in the degradation of uridine into uridine monophosphate through reduction of guanosine triphosphate). However, the deletion of these genes did not affect the MIC of tobramycin (Table S1). Along with the results from nucleoside transporter overexpression, these findings suggest that uridine utilization is not essential for the potentiating effect of uridine on AGs. Moreover, the observation that other nucleosides can also decrease the MIC of AGs implies that the phenotype is not specifically linked to uridine utilization pathways. **A.** Susceptibility to tobramycin of *E. coli* carrying the empty plasmid (p0) compared to the vector overexpressing NupG or NupC, two uridine transporters. Cultures of each strain were grown overnight. Susceptibility was assessed by serial dilutions and dropping 5 µl of each dilution on plates containing or not 0.1 µg/ml of tobramycin and uridine 0.5%. Inducer was added in the media. **B.** Additionally, we sought to investigate the possible association between the effects of uridine and the stringent response. Previous research has indicated that the stringent response may play a role in the uptake of streptomycin [62] and the susceptibility to AGs [63,64]. Furthermore, SpoT, an enzyme involved in ppGpp synthesis, has been found to be influenced by carbon sources [65]. To assess whether uridine supplementation could exert its potentiating effect on AGs via modulation of the stringent response, we employed a reporter plasmid containing the *rrnB* 16S ribosomal RNA promoter 1 fused to GFP [57]. This promoter is negatively regulated by ppGpp [64] decrease in fluorescence intensity thus indicates an induction of the stringent response. As controls, tobramycin sub-MIC [66,67] treatment and stationary phase cultures were used, as both are known to trigger the stringent response. In our experimental conditions, treatment with various carbon sources, including uridine, did not induce the stringent response Quantification of GFP fluorescence in exponential phase from the P1*rrnB* using flow cytometry depending on the substrate added in the media (supplemented or not with glucose, uridine, ribose or maltose 0.5 %) expressed as fold change of mean fluorescence per cell. Cultures in stationary phase (Stat) and cells treated with sub-MIC tobramycin (Tob, 0.06 µg/ml) were used as positive controls of stringent response activation. For statistical significance calculations, we used one-way ANOVA. **** means $p<0.0001$, ns: non-significant (Glucose, Uridine, Ribose, Maltose compared to MH). Number of replicates for each experiment: n=3. The significance corresponds to comparison with the MH condition.

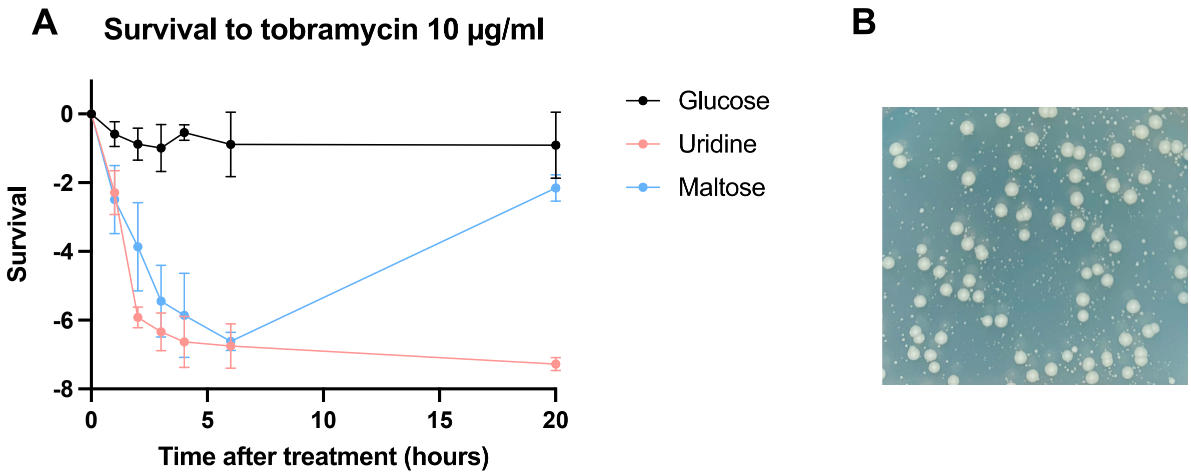

**Figure S8: Uridine induces fast killing and prevent the appearance of mutants**. Survival of *E. coli* WT, growing in mid-exponential phase in liquid cultures supplemented or not with glucose, maltose or uridine 0.5%. Lethal treatment of tobramycin (10 µg/ml) was applied, and survival was assessed after 1, 2, 3, 4, 6 and 20 hours by plating and counting CFUs/ml. Geometric mean and geometric standard deviation on three biological replicates are represented.

Regrowth was observed in the maltose-treated bacteria after 20 hours of treatment. We hypothesized that this regrowth could be attributed to the selection of resistant mutants growing after 6 hours of treatment, as the number of surviving bacteria after 20 hours in the presence of tobramycin was higher than at 6 hours of treatment. To investigate further, we sequenced the genomic DNA from 10 colonies which exhibited a MIC around 1 µg/ml. We identified mutations in the *fusA* gene, which encodes elongation factor G and the *rplL* gene, which encodes 50S ribosomal protein L7/L12. These genes are known to be involved in the translation process affected by AGs.

**Supplementary tables**

**Table S1** : MIC (µg/ml) determined by Etest. (Tob: Tobramycin; Kan: kanamycin; Gen: gentamicin; Cip: ciprofloxacin; Chlo: chloramphenicol)

| **Gene deleted in *E. coli* and associated substrate** | Tob |
|---|---|
| WT | 0.1 |
| *lamB* - Maltose | 0.1 |
| *malE* - Maltose | 0.12 |
| *manY* - Mannose | 0.2 |
| *fruA* - Fructose | 0.12 |
| *bglH* – ß-glucoside | 0.1 |
| *frwB* - Fructose | 0.1 |
| *mngA* – Fructose-like | 0.12 |
| *cmtA* - Mannitol (cryptic) | 0.4 |
| *ptsG* - Glucose | 0.1 |
| *mglA* - Galactose | 0.2 |
| *chbC* - Di-N-acetylchitobiose/Cellobiose | 0.15 |
| *nagE* - Acetylglucosamine | 0.12 |
| *ypdG* - Fructose-like | 0.1 |
| *ascF* - Cellobiose/arbutine | 0.1 |
| *agaW* - N-acetylgalactosamine | 0.1 |
| *sgcC* -Pentose | 0.1 |
| *malX* - Maltose | 0.12 |
| *mtlA* - Mannitol | 0.1 |
| *treB* - Trehalose | 0.1 |
| *frvB* - Fructose-like | 0.1 |
| *agaC* - Galactosamine | 0.1 |
| *srlE* - Glucitol | 0.12 |

| | | | | | | |
|---|---|---|---|---|---|---|
| *glvC* - Arbutine | 0.1 | | | | | |
| *crp* | 1.5 | | | | | |
| *udk* | 0.1 | | | | | |
| *udp* | 0.1 | | | | | |
| **Overexpressed gene in *E. coli*** | Tob | | | | | |
| p0 (empty vector) | 0.1 | | | | | |
| *lamB* | 0.1 | | | | | |
| *malE* | 0.08 | | | | | |
| *fruA* | 0.08 | | | | | |
| *frwB* | 0.1 | | | | | |
| *mngA* | 0.08 | | | | | |
| *cmtA* | <0.064 | | | | | |
| *chbC* | <0.064 | | | | | |
| *ascF* | 0.064 | | | | | |
| *malX* | 0.064 | | | | | |
| *frvAB* | 0.1 | | | | | |
| *srlE* | 0.064 | | | | | |
| *nupG* | 0.1 | | | | | |
| *nupC* | 0.1 | | | | | |
| **Substrate supplementation, E. coli WT** | Tob | Gen | Kan | Cip | Chlo | Amp |
| Glucose | 1.5 | 2 | 2 | 0.12 | 1.5 | 1.5 |
| Uridine | 0.1 | 0.1 | 0.75 | 0.06 | 1.5 | 1.5 |
| Ribose | 1 | 1 | 2 | 0.06 | 1.5 | 1 |
| Glycerol | 0.4 | | | | | |
| Fructose | 1 | | | | | |
| Galactose | 1 | | | | | |

| | | |
|---|---|---|
| Mannitol | 1 | |
| D-galactosamine | 0.3 | |
| Mannose | 1 | |
| Maltose | 0.4 | |
| Cytidine | 0.1 | 0.1 |
| Adenosine | 0.1 | 0.15 |
| Thymidine | 0.4 | |
| Inosine | 1 | 1.5 |
| **Substrate supplementation, E. coli Δcrp** | Tob | |
| Glucose | 6 | |
| Uridine | 6 | |

| **MIC to tobramycin in different species according to the substrate supplementation** | *V. cholerae* | *P. aeruginosa* | *K. pneumoniae* | *A. baumannii* |
|---|---|---|---|---|
| Glucose | 6 | 1 | 1.5 | 4 |
| Ribose | | | 0.50 | 1 |
| Uridine | 0.5 | 0.25 | 0.38 | 0.5 |
| **Overexpressed gene in *P. aeruginosa* and predicted substrate** | Tob | Gen | Chlo | Cip |
| p0 | 4 | 4 | 16 | 0.094 |
| *oprB* - Glucose/mannitol/fructose/ glycerol | 1.5 | 2 | 16 | 0.094 |
| *gtsB* - Mannose/glucose | 0.4 | 1.5 | 16 | 0.094 |
| *mtlFGK* - Maltose/mannitol | 0.4 | 1.5 | 16 | 0.094 |
| *oprB2* (PA2291) - *oprB* homolog | 1 | 1 | 16 | 0.094 |
| *FruA* - Fructose | 0.4 | 1.5 | 16 | 0.094 |

| **Overexpressed gene in *A. baumannii* and predicted substrate** | |
|---|---|
| p0 | 0.1 |
| *FruA* - Fructose | <0.064 |

**Table S2:** Screening data using PcmtA-GFP reporter on Biolog plates (PM1, PM2A and PM3B). Ratio as calculated in the method section are indicated on the last line.

**Plate PM1:**

| OD600nm | Uridine | Bromo-succin | Inosine | L-alanine | D-alanine | D-Threonine | Fumaric Acid | Adenosine | D-L malic acid | D-malic-acid | a-glycerol-pho | α-Hydroxy Bu | G-glucose-1-p | L-Alanyl-Glyci | D-methyl-D-gl | Glycyl-L-Aspar |
|---|---|---|---|---|---|---|---|---|---|---|---|---|---|---|---|---|
| 0s | 0,0557 | 0,0553 | 0,0558 | 0,0541 | 0,053 | 0,0502 | 0,0564 | 0,0539 | 0,0525 | 0,055 | 0,052 | 0,0562 | 0,0516 | 0,0571 | 0,0589 | 0,052 |
| 897s | 0,0524 | 0,0549 | 0,0661 | 0,0596 | 0,0528 | 0,0507 | 0,0565 | 0,0585 | 0,0538 | 0,0532 | 0,0525 | 0,06 | 0,0524 | 0,0593 | 0,0588 | 0,0526 |
| 1154s | 0,0541 | 0,055 | 0,0585 | 0,0569 | 0,0568 | 0,0512 | 0,0536 | 0,0566 | 0,0533 | 0,0564 | 0,0538 | 0,0609 | 0,0527 | 0,0576 | 0,0563 | 0,0534 |
| 1412s | 0,0544 | 0,0552 | 0,0612 | 0,0574 | 0,0544 | 0,0512 | 0,0533 | 0,0628 | 0,0539 | 0,0561 | 0,0544 | 0,056 | 0,0534 | 0,0567 | 0,061 | 0,0534 |
| 1669s | 0,0552 | 0,0562 | 0,0661 | 0,0612 | 0,0561 | 0,0528 | 0,055 | 0,0576 | 0,0543 | 0,0566 | 0,0556 | 0,0566 | 0,0539 | 0,062 | 0,0618 | 0,0537 |
| 1926s | 0,0558 | 0,0567 | 0,06 | 0,059 | 0,0541 | 0,0526 | 0,0605 | 0,055 | 0,0545 | 0,0583 | 0,0552 | 0,0588 | 0,0542 | 0,0609 | 0,0571 | 0,0545 |
| 2183s | 0,0558 | 0,057 | 0,0645 | 0,0615 | 0,0544 | 0,0537 | 0,0553 | 0,0554 | 0,0553 | 0,0625 | 0,0553 | 0,0608 | 0,0551 | 0,0644 | 0,0612 | 0,0561 |
| 2711s | 0,0586 | 0,0575 | 0,0653 | 0,0618 | 0,0551 | 0,0545 | 0,0572 | 0,0636 | 0,0573 | 0,0624 | 0,0575 | 0,0609 | 0,0574 | 0,0637 | 0,0564 | 0,0566 |
| 3611s | 0,0619 | 0,0592 | 0,0677 | 0,0624 | 0,0554 | 0,0559 | 0,0592 | 0,0589 | 0,0594 | 0,0565 | 0,0611 | 0,0598 | 0,0611 | 0,0623 | 0,0579 | 0,0577 |
| 4511s | 0,0689 | 0,0597 | 0,0685 | 0,0651 | 0,0593 | 0,0555 | 0,06 | 0,0634 | 0,0604 | 0,0584 | 0,0653 | 0,0646 | 0,0659 | 0,0661 | 0,0584 | 0,0585 |
| 5411s | 0,0681 | 0,062 | 0,0697 | 0,0639 | 0,0623 | 0,0568 | 0,0642 | 0,0688 | 0,0611 | 0,0638 | 0,07 | 0,0654 | 0,0699 | 0,0623 | 0,06 | 0,0597 |
| 6311s | 0,0701 | 0,0641 | 0,0649 | 0,0673 | 0,0576 | 0,0609 | 0,0716 | 0,0677 | 0,0634 | 0,0635 | 0,072 | 0,0679 | 0,0737 | 0,0666 | 0,0624 | 0,0631 |
| 7212s | 0,0919 | 0,0704 | 0,0733 | 0,0705 | 0,0606 | 0,0659 | 0,076 | 0,0777 | 0,0696 | 0,0661 | 0,0787 | 0,07 | 0,0842 | 0,0714 | 0,0657 | 0,0695 |
| 8112s | 0,1008 | 0,0767 | 0,0978 | 0,0759 | 0,0627 | 0,0736 | 0,0856 | 0,0897 | 0,0776 | 0,0773 | 0,0907 | 0,0784 | 0,0938 | 0,081 | 0,0745 | 0,078 |
| 9012s | 0,1125 | 0,0861 | 0,1187 | 0,0825 | 0,071 | 0,0812 | 0,0916 | 0,1061 | 0,0873 | 0,0846 | 0,1061 | 0,0928 | 0,1099 | 0,0902 | 0,085 | 0,0898 |
| 9912s | 0,1188 | 0,0992 | 0,1291 | 0,0906 | 0,0736 | 0,0914 | 0,1083 | 0,1214 | 0,0958 | 0,0955 | 0,1139 | 0,1037 | 0,1224 | 0,1014 | 0,0982 | 0,1021 |
| 10812s | 0,1308 | 0,1122 | 0,1367 | 0,1017 | 0,0872 | 0,0977 | 0,1262 | 0,137 | 0,1009 | 0,1091 | 0,1281 | 0,116 | 0,1332 | 0,1115 | 0,1072 | 0,1172 |
| 11712s | 0,1425 | 0,1272 | 0,1494 | 0,1153 | 0,0959 | 0,1089 | 0,1374 | 0,1474 | 0,1188 | 0,1253 | 0,1477 | 0,1293 | 0,1471 | 0,1311 | 0,1296 | 0,1264 |
| 12612s | 0,165 | 0,1392 | 0,1573 | 0,1269 | 0,1006 | 0,1211 | 0,147 | 0,1551 | 0,1319 | 0,143 | 0,1597 | 0,1419 | 0,1551 | 0,1392 | 0,1361 | 0,1329 |
| 13512s | 0,1575 | 0,1468 | 0,1651 | 0,139 | 0,0969 | 0,1305 | 0,1476 | 0,1649 | 0,1395 | 0,1455 | 0,1782 | 0,1557 | 0,1657 | 0,1521 | 0,1577 | 0,1473 |
| 14412s | 0,1676 | 0,1531 | 0,1714 | 0,1465 | 0,0972 | 0,1329 | 0,1606 | 0,1739 | 0,1433 | 0,154 | 0,1859 | 0,1549 | 0,1761 | 0,1523 | 0,1571 | 0,155 |
| 15312s | 0,1644 | 0,1525 | 0,1771 | 0,1473 | 0,1072 | 0,1329 | 0,1605 | 0,1816 | 0,1472 | 0,1604 | 0,1872 | 0,1539 | 0,1859 | 0,1494 | 0,1639 | 0,1609 |
| 16212s | 0,1902 | 0,1586 | 0,1874 | 0,1568 | 0,1129 | 0,1361 | 0,1706 | 0,1995 | 0,1759 | 0,1621 | 0,1968 | 0,1593 | 0,2012 | 0,156 | 0,1532 | 0,1699 |
| 17112s | 0,1973 | 0,1608 | 0,1986 | 0,1582 | 0,1211 | 0,1423 | 0,168 | 0,2036 | 0,1631 | 0,1701 | 0,1999 | 0,1682 | 0,2116 | 0,1599 | 0,1625 | 0,1673 |
| 18013s | 0,1927 | 0,1727 | 0,2001 | 0,1671 | 0,1193 | 0,1488 | 0,214 | 0,2096 | 0,183 | 0,1774 | 0,206 | 0,1794 | 0,2358 | 0,167 | 0,1688 | 0,1783 |
| 18913s | 0,2004 | 0,1733 | 0,2101 | 0,1652 | 0,1217 | 0,1501 | 0,2095 | 0,2161 | 0,1833 | 0,1834 | 0,2126 | 0,1814 | 0,2524 | 0,1657 | 0,1857 | 0,1799 |
| 19813s | 0,1935 | 0,1761 | 0,2063 | 0,1764 | 0,1169 | 0,1517 | 0,2271 | 0,2155 | 0,1939 | 0,1815 | 0,21 | 0,1837 | 0,2621 | 0,1669 | 0,1779 | 0,1751 |
| 20713s | 0,194 | 0,1803 | 0,2093 | 0,1748 | 0,1302 | 0,1559 | 0,2297 | 0,2182 | 0,1913 | 0,1837 | 0,2114 | 0,1923 | 0,2736 | 0,1722 | 0,1874 | 0,1795 |
| 21613s | 0,2191 | 0,2023 | 0,2188 | 0,1813 | 0,1273 | 0,1596 | 0,2604 | 0,224 | 0,1923 | 0,184 | 0,2169 | 0,1946 | 0,3041 | 0,1866 | 0,2073 | 0,1791 |
| 22513s | 0,2111 | 0,2158 | 0,229 | 0,1828 | 0,1286 | 0,1663 | 0,2369 | 0,2339 | 0,1819 | 0,1929 | 0,2185 | 0,1979 | 0,2897 | 0,1855 | 0,2076 | 0,1805 |
| 23413s | 0,2207 | 0,2262 | 0,2306 | 0,1861 | 0,1309 | 0,1702 | 0,2667 | 0,256 | 0,1862 | 0,1906 | 0,2177 | 0,1984 | 0,3002 | 0,1913 | 0,2115 | 0,1818 |
| 24313s | 0,2166 | 0,2187 | 0,2317 | 0,1859 | 0,141 | 0,1723 | 0,2339 | 0,2459 | 0,1912 | 0,1963 | 0,2223 | 0,197 | 0,2898 | 0,1893 | 0,2013 | 0,1819 |
| 25213s | 0,2177 | 0,211 | 0,2349 | 0,1903 | 0,1399 | 0,1713 | 0,2318 | 0,2453 | 0,1938 | 0,208 | 0,224 | 0,2002 | 0,2929 | 0,1879 | 0,2346 | 0,179 |
| 26114s | 0,2175 | 0,2185 | 0,2381 | 0,1944 | 0,148 | 0,1759 | 0,2424 | 0,242 | 0,2014 | 0,198 | 0,2261 | 0,2038 | 0,2967 | 0,1959 | 0,2415 | 0,1813 |
| 27014s | 0,2159 | 0,2201 | 0,2348 | 0,1977 | 0,147 | 0,1776 | 0,2396 | 0,2433 | 0,1979 | 0,1972 | 0,2232 | 0,2044 | 0,2958 | 0,1992 | 0,2373 | 0,1808 |
| 27914s | 0,2316 | 0,2322 | 0,2438 | 0,2025 | 0,1477 | 0,1793 | 0,2537 | 0,2508 | 0,2008 | 0,2055 | 0,2276 | 0,209 | 0,306 | 0,2043 | 0,2626 | 0,1815 |
| 28814s | 0,2227 | 0,2264 | 0,2454 | 0,2054 | 0,143 | 0,1818 | 0,2382 | 0,2392 | 0,2028 | 0,1998 | 0,2281 | 0,2082 | 0,2911 | 0,2046 | 0,258 | 0,1825 |
| 29714s | 0,2332 | 0,2282 | 0,2435 | 0,2051 | 0,1418 | 0,1812 | 0,2201 | 0,2488 | 0,206 | 0,2022 | 0,23 | 0,2173 | 0,2935 | 0,2081 | 0,2436 | 0,1839 |
| 30614s | 0,2362 | 0,232 | 0,2442 | 0,2081 | 0,1322 | 0,1819 | 0,2272 | 0,2453 | 0,2084 | 0,2135 | 0,2331 | 0,2126 | 0,2912 | 0,2162 | 0,2534 | 0,1849 |
| 31514s | 0,2232 | 0,2303 | 0,2527 | 0,2098 | 0,1299 | 0,1872 | 0,2269 | 0,24 | 0,2103 | 0,2153 | 0,2391 | 0,2142 | 0,2949 | 0,2166 | 0,2542 | 0,1853 |
| 32414s | 0,2051 | 0,2311 | 0,2548 | 0,2101 | 0,1269 | 0,1862 | 0,2285 | 0,2321 | 0,2105 | 0,2213 | 0,2419 | 0,2154 | 0,2914 | 0,2169 | 0,2514 | 0,1873 |

| GFP | Uridine | Bromo-succin | Inosine | L-alanine | D-alanine | D-Threonine | Fumaric Acid | Adenosine | D-L malic acid | D-malic-acid | a-glycerol-pho | α-Hydroxy Bu | G-glucose-1-p | L-Alanyl-Glyci | D-methyl-D-gl | Glycyl-L-Aspar |
|---|---|---|---|---|---|---|---|---|---|---|---|---|---|---|---|---|
| 0s | 143 | 156 | 143 | 171 | 170 | 136 | 166 | 161 | 170 | 165 | 179 | 139 | 172 | 129 | 133 | 164 |
| 897s | 141 | 151 | 139 | 156 | 158 | 130 | 165 | 162 | 170 | 156 | 168 | 136 | 172 | 125 | 128 | 165 |
| 1154s | 137 | 145 | 145 | 161 | 154 | 133 | 164 | 162 | 165 | 156 | 172 | 140 | 165 | 125 | 128 | 161 |
| 1411s | 137 | 146 | 141 | 158 | 158 | 136 | 164 | 158 | 165 | 155 | 163 | 141 | 165 | 125 | 127 | 163 |
| 1668s | 142 | 149 | 140 | 167 | 160 | 128 | 167 | 154 | 164 | 161 | 165 | 139 | 165 | 124 | 126 | 160 |
| 1926s | 143 | 150 | 142 | 162 | 164 | 137 | 170 | 160 | 163 | 160 | 168 | 141 | 164 | 127 | 130 | 165 |
| 2183s | 143 | 149 | 144 | 161 | 163 | 134 | 165 | 161 | 170 | 160 | 163 | 143 | 167 | 128 | 130 | 158 |
| 2711s | 142 | 149 | 143 | 165 | 163 | 142 | 165 | 159 | 168 | 164 | 168 | 148 | 166 | 134 | 131 | 165 |
| 3611s | 144 | 158 | 140 | 172 | 164 | 138 | 176 | 162 | 176 | 165 | 167 | 148 | 169 | 139 | 136 | 168 |
| 4511s | 141 | 158 | 134 | 170 | 164 | 141 | 174 | 154 | 174 | 167 | 179 | 155 | 166 | 142 | 135 | 169 |
| 5411s | 144 | 157 | 135 | 170 | 164 | 141 | 176 | 158 | 168 | 162 | 175 | 150 | 158 | 139 | 143 | 171 |
| 6311s | 154 | 164 | 140 | 178 | 170 | 142 | 175 | 161 | 177 | 172 | 181 | 159 | 160 | 142 | 142 | 173 |
| 7211s | 160 | 177 | 147 | 181 | 173 | 155 | 187 | 170 | 188 | 184 | 191 | 166 | 183 | 151 | 146 | 181 |
| 8111s | 180 | 192 | 168 | 199 | 183 | 161 | 201 | 183 | 198 | 190 | 198 | 180 | 199 | 168 | 156 | 191 |
| 9011s | 204 | 212 | 198 | 217 | 201 | 186 | 217 | 196 | 206 | 211 | 216 | 202 | 215 | 185 | 165 | 207 |
| 9912s | 252 | 239 | 215 | 233 | 211 | 200 | 239 | 233 | 226 | 224 | 227 | 221 | 230 | 206 | 183 | 216 |
| 10812s | 298 | 256 | 251 | 255 | 223 | 205 | 275 | 269 | 232 | 239 | 277 | 253 | 274 | 212 | 199 | 244 |
| 11712s | 368 | 316 | 312 | 285 | 241 | 232 | 327 | 326 | 285 | 291 | 311 | 301 | 323 | 271 | 213 | 293 |
| 12612s | 436 | 364 | 384 | 316 | 267 | 292 | 366 | 382 | 322 | 352 | 354 | 338 | 382 | 298 | 254 | 316 |
| 13512s | 511 | 414 | 440 | 383 | 302 | 323 | 411 | 438 | 372 | 388 | 390 | 384 | 440 | 330 | 284 | 384 |
| 14412s | 590 | 482 | 525 | 442 | 335 | 361 | 461 | 502 | 428 | 430 | 439 | 427 | 484 | 375 | 337 | 441 |
| 15312s | 658 | 523 | 593 | 480 | 348 | 397 | 517 | 561 | 475 | 484 | 489 | 463 | 532 | 389 | 376 | 495 |
| 16212s | 721 | 582 | 646 | 541 | 437 | 422 | 567 | 622 | 582 | 524 | 558 | 500 | 577 | 419 | 415 | 531 |
| 17112s | 793 | 619 | 720 | 590 | 493 | 465 | 644 | 691 | 577 | 571 | 587 | 539 | 619 | 450 | 455 | 564 |
| 18012s | 827 | 687 | 776 | 650 | 552 | 495 | 809 | 741 | 682 | 613 | 655 | 580 | 680 | 488 | 494 | 608 |
| 18912s | 903 | 707 | 833 | 677 | 599 | 534 | 844 | 805 | 712 | 655 | 716 | 628 | 741 | 527 | 528 | 621 |
| 19813s | 972 | 742 | 884 | 737 | 636 | 583 | 916 | 865 | 762 | 686 | 764 | 663 | 832 | 549 | 572 | 633 |
| 20713s | 1043 | 786 | 948 | 769 | 659 | 615 | 1006 | 910 | 792 | 733 | 822 | 698 | 958 | 579 | 632 | 660 |
| 21613s | 1101 | 918 | 1003 | 812 | 671 | 673 | 1133 | 974 | 806 | 778 | 868 | 754 | 1074 | 632 | 708 | 687 |
| 22513s | 1166 | 956 | 1049 | 864 | 689 | 714 | 1087 | 1032 | 839 | 839 | 936 | 798 | 1171 | 668 | 739 | 700 |
| 23413s | 1214 | 1060 | 1115 | 913 | 702 | 756 | 1233 | 1125 | 879 | 860 | 998 | 842 | 1285 | 713 | 827 | 712 |
| 24313s | 1311 | 1070 | 1198 | 956 | 720 | 820 | 1134 | 1160 | 925 | 909 | 1050 | 886 | 1316 | 756 | 822 | 729 |
| 25213s | 1316 | 1080 | 1245 | 995 | 732 | 841 | 1175 | 1235 | 969 | 958 | 1103 | 928 | 1415 | 788 | 927 | 746 |
| 26113s | 1395 | 1190 | 1319 | 1036 | 747 | 880 | 1220 | 1282 | 998 | 994 | 1150 | 1008 | 1477 | 817 | 1031 | 776 |
| 27013s | 1439 | 1260 | 1392 | 1083 | 779 | 920 | 1245 | 1349 | 1023 | 1032 | 1202 | 1038 | 1510 | 867 | 1078 | 814 |
| 27913s | 1499 | 1373 | 1460 | 1130 | 780 | 964 | 1313 | 1382 | 1057 | 1070 | 1251 | 1097 | 1636 | 923 | 1263 | 860 |
| 28814s | 1553 | 1435 | 1530 | 1169 | 802 | 980 | 1322 | 1398 | 1083 | 1141 | 1292 | 1132 | 1618 | 960 | 1278 | 888 |
| 29714s | 1633 | 1443 | 1603 | 1230 | 790 | 1012 | 1288 | 1439 | 1127 | 1172 | 1356 | 1135 | 1659 | 1018 | 1260 | 926 |
| 30614s | 1641 | 1527 | 1676 | 1274 | 790 | 1089 | 1339 | 1444 | 1191 | 1212 | 1407 | 1154 | 1708 | 1072 | 1302 | 978 |
| 31514s | 1632 | 1552 | 1740 | 1335 | 766 | 1116 | 1360 | 1439 | 1246 | 1276 | 1441 | 1200 | 1763 | 1133 | 1361 | 1014 |
| 32414s | 1631 | 1619 | 1757 | 1388 | 742 | 1166 | 1412 | 1446 | 1282 | 1316 | 1480 | 1215 | 1788 | 1193 | 1399 | 1046 |
| ratio | 9959,83936 | 8321,95677 | 8110,55276 | 7801,28205 | 7740,18945 | 7573,52941 | 7239,97676 | 7210,99888 | 7037,97468 | 6921,2267 | 6850,9742 | 6758,79397 | 6738,94912 | 6658,3229 | 6576,62338 | 6518,84701 |

| α-Keto-butyric acid | L-asparagine | α-Keto-Glutaric | L-aspartic acic | L-threonine | N-Acetyl-ß-Dn | α-Hydroxy Glu | Propionic Aci | D-trehalose | L-Malic Acid | D-glucosamic | Succinic Acid | Phenylethylar | 2-deoxy-aden | Mono Methyl | Glyoxylic Acid | L-Lactic acid | D-Ribose |
|---|---|---|---|---|---|---|---|---|---|---|---|---|---|---|---|---|---|
| 0,0577 | 0,0514 | 0,0532 | 0,0511 | 0,0527 | 0,0546 | 0,0552 | 0,0602 | 0,0523 | 0,0543 | 0,052 | 0,047 | 0,0604 | 0,055 | 0,0562 | 0,0548 | 0,0528 | 0,0539 |
| 0,0618 | 0,052 | 0,0529 | 0,0527 | 0,0517 | 0,0551 | 0,0572 | 0,0629 | 0,056 | 0,0537 | 0,0529 | 0,0484 | 0,0559 | 0,0632 | 0,0623 | 0,0555 | 0,0551 | 0,0546 |
| 0,0649 | 0,0523 | 0,0553 | 0,0537 | 0,053 | 0,0561 | 0,0593 | 0,0623 | 0,0551 | 0,0553 | 0,0532 | 0,0492 | 0,0569 | 0,0619 | 0,065 | 0,0564 | 0,0558 | 0,0547 |
| 0,0609 | 0,0527 | 0,0584 | 0,0545 | 0,0548 | 0,057 | 0,0622 | 0,0604 | 0,0558 | 0,0545 | 0,0535 | 0,0497 | 0,0561 | 0,0563 | 0,0597 | 0,0574 | 0,0559 | 0,055 |
| 0,0579 | 0,0541 | 0,0593 | 0,0548 | 0,0533 | 0,0566 | 0,0615 | 0,0608 | 0,0579 | 0,0555 | 0,0541 | 0,0501 | 0,0552 | 0,057 | 0,0549 | 0,0573 | 0,0576 | 0,0555 |
| 0,0634 | 0,0544 | 0,0572 | 0,0553 | 0,0536 | 0,0578 | 0,0619 | 0,0637 | 0,0572 | 0,0607 | 0,0546 | 0,0516 | 0,0589 | 0,0532 | 0,0562 | 0,0581 | 0,0573 | 0,0557 |
| 0,0563 | 0,0559 | 0,0577 | 0,0547 | 0,0563 | 0,0583 | 0,0585 | 0,0668 | 0,0591 | 0,0592 | 0,0557 | 0,0518 | 0,06 | 0,0559 | 0,0558 | 0,0581 | 0,0551 | 0,0566 |
| 0,0587 | 0,0562 | 0,0671 | 0,056 | 0,0556 | 0,0592 | 0,0617 | 0,0614 | 0,0611 | 0,0592 | 0,0568 | 0,0537 | 0,0588 | 0,0572 | 0,0592 | 0,0596 | 0,0571 | 0,0579 |
| 0,0599 | 0,0582 | 0,0575 | 0,0582 | 0,0567 | 0,0624 | 0,0641 | 0,0572 | 0,0639 | 0,063 | 0,0584 | 0,0563 | 0,0626 | 0,0625 | 0,0563 | 0,0612 | 0,06 | 0,0613 |
| 0,0642 | 0,0595 | 0,0663 | 0,0594 | 0,0583 | 0,0644 | 0,0627 | 0,0626 | 0,0682 | 0,0632 | 0,0592 | 0,0574 | 0,0607 | 0,0678 | 0,0576 | 0,0619 | 0,0617 | 0,0648 |
| 0,0647 | 0,0608 | 0,0625 | 0,0608 | 0,0602 | 0,0643 | 0,0641 | 0,0573 | 0,071 | 0,0663 | 0,0603 | 0,0591 | 0,0604 | 0,0714 | 0,0609 | 0,063 | 0,0656 | 0,0677 |
| 0,0609 | 0,0641 | 0,0644 | 0,0676 | 0,0632 | 0,0704 | 0,0689 | 0,0602 | 0,069 | 0,0652 | 0,0631 | 0,0639 | 0,0626 | 0,0635 | 0,0633 | 0,0639 | 0,0703 | 0,068 |
| 0,0657 | 0,0703 | 0,0704 | 0,0731 | 0,0701 | 0,075 | 0,0731 | 0,0713 | 0,0769 | 0,0707 | 0,0688 | 0,0691 | 0,0647 | 0,0673 | 0,0698 | 0,0681 | 0,0742 | 0,0721 |
| 0,0833 | 0,0799 | 0,0793 | 0,0825 | 0,0799 | 0,0849 | 0,0819 | 0,0784 | 0,088 | 0,0842 | 0,0769 | 0,0791 | 0,0697 | 0,0827 | 0,0801 | 0,0751 | 0,086 | 0,08 |
| 0,0821 | 0,0871 | 0,0921 | 0,0928 | 0,0907 | 0,0989 | 0,0919 | 0,0925 | 0,0997 | 0,0926 | 0,0868 | 0,089 | 0,0767 | 0,0854 | 0,089 | 0,0831 | 0,097 | 0,0908 |
| 0,0967 | 0,0984 | 0,1062 | 0,105 | 0,1059 | 0,116 | 0,1059 | 0,1014 | 0,1095 | 0,104 | 0,0964 | 0,1031 | 0,0843 | 0,1012 | 0,1005 | 0,0926 | 0,1063 | 0,098 |
| 0,1066 | 0,1069 | 0,1218 | 0,1159 | 0,1185 | 0,13 | 0,1205 | 0,1085 | 0,1246 | 0,1237 | 0,1049 | 0,1092 | 0,0953 | 0,1151 | 0,1107 | 0,101 | 0,1193 | 0,1124 |
| 0,1111 | 0,126 | 0,1376 | 0,1338 | 0,1313 | 0,1459 | 0,1323 | 0,1233 | 0,1371 | 0,134 | 0,1181 | 0,1207 | 0,1015 | 0,1217 | 0,132 | 0,1181 | 0,1332 | 0,1258 |
| 0,1263 | 0,1374 | 0,1574 | 0,1411 | 0,1411 | 0,1593 | 0,1442 | 0,1381 | 0,1465 | 0,1506 | 0,134 | 0,1323 | 0,1101 | 0,1451 | 0,1487 | 0,1376 | 0,1454 | 0,1376 |
| 0,1419 | 0,1464 | 0,1541 | 0,135 | 0,1468 | 0,1636 | 0,1525 | 0,1504 | 0,1578 | 0,1548 | 0,1416 | 0,1315 | 0,1187 | 0,1534 | 0,1512 | 0,1453 | 0,1534 | 0,1481 |
| 0,1532 | 0,1478 | 0,1606 | 0,1358 | 0,1557 | 0,1619 | 0,1539 | 0,1639 | 0,1564 | 0,1508 | 0,1466 | 0,1325 | 0,1179 | 0,1669 | 0,1561 | 0,151 | 0,1604 | 0,1573 |
| 0,1572 | 0,158 | 0,1675 | 0,142 | 0,1579 | 0,167 | 0,155 | 0,1637 | 0,1656 | 0,1523 | 0,1529 | 0,1374 | 0,1173 | 0,1754 | 0,1636 | 0,1553 | 0,1664 | 0,1643 |
| 0,1666 | 0,1736 | 0,1694 | 0,1425 | 0,1625 | 0,1814 | 0,1616 | 0,1622 | 0,1724 | 0,1595 | 0,1577 | 0,1403 | 0,1223 | 0,183 | 0,1681 | 0,1691 | 0,1675 | 0,1755 |
| 0,1703 | 0,1712 | 0,1753 | 0,1509 | 0,1635 | 0,1895 | 0,1686 | 0,1731 | 0,1752 | 0,1665 | 0,1593 | 0,1441 | 0,1199 | 0,1941 | 0,174 | 0,1703 | 0,1712 | 0,1805 |
| 0,1697 | 0,1802 | 0,1842 | 0,1524 | 0,1665 | 0,1913 | 0,1804 | 0,1862 | 0,1814 | 0,1731 | 0,1788 | 0,1473 | 0,1265 | 0,2113 | 0,175 | 0,1826 | 0,1796 | 0,1877 |
| 0,1775 | 0,1739 | 0,1865 | 0,1584 | 0,1705 | 0,1947 | 0,1883 | 0,1868 | 0,1848 | 0,177 | 0,1788 | 0,1535 | 0,1234 | 0,2209 | 0,1818 | 0,1847 | 0,1809 | 0,1991 |
| 0,1815 | 0,1785 | 0,2001 | 0,1544 | 0,1796 | 0,1937 | 0,1811 | 0,1918 | 0,192 | 0,1778 | 0,1866 | 0,1556 | 0,1217 | 0,2226 | 0,1812 | 0,1841 | 0,1864 | 0,2034 |
| 0,1862 | 0,1818 | 0,1966 | 0,1574 | 0,1831 | 0,1988 | 0,1884 | 0,1825 | 0,1992 | 0,1831 | 0,1911 | 0,1565 | 0,1284 | 0,2376 | 0,1854 | 0,1891 | 0,1877 | 0,2138 |
| 0,1888 | 0,1803 | 0,2133 | 0,1592 | 0,1856 | 0,2003 | 0,218 | 0,2006 | 0,2053 | 0,181 | 0,1986 | 0,1587 | 0,1271 | 0,2466 | 0,1904 | 0,1986 | 0,1997 | 0,2154 |
| 0,1932 | 0,1847 | 0,2066 | 0,1695 | 0,1893 | 0,2058 | 0,2229 | 0,2067 | 0,2153 | 0,1916 | 0,1962 | 0,1624 | 0,1294 | 0,2531 | 0,2038 | 0,2038 | 0,198 | 0,2256 |
| 0,2032 | 0,182 | 0,2097 | 0,1676 | 0,194 | 0,2103 | 0,2379 | 0,219 | 0,2198 | 0,1877 | 0,2013 | 0,1642 | 0,1287 | 0,2456 | 0,1997 | 0,2041 | 0,197 | 0,231 |
| 0,2068 | 0,1806 | 0,2125 | 0,1776 | 0,1954 | 0,2139 | 0,2286 | 0,2178 | 0,2241 | 0,188 | 0,1985 | 0,1673 | 0,1277 | 0,2451 | 0,2105 | 0,201 | 0,2082 | 0,2323 |
| 0,2124 | 0,18 | 0,2251 | 0,1822 | 0,1989 | 0,2206 | 0,222 | 0,2079 | 0,2233 | 0,1821 | 0,1945 | 0,1717 | 0,1337 | 0,2573 | 0,2116 | 0,2075 | 0,2163 | 0,2389 |
| 0,2124 | 0,1771 | 0,2302 | 0,1817 | 0,2034 | 0,2178 | 0,2225 | 0,2091 | 0,2268 | 0,1837 | 0,1958 | 0,1762 | 0,1318 | 0,2542 | 0,2015 | 0,2061 | 0,2172 | 0,2457 |
| 0,2057 | 0,1774 | 0,2317 | 0,1805 | 0,2075 | 0,2221 | 0,2471 | 0,2146 | 0,2278 | 0,1829 | 0,2017 | 0,178 | 0,1232 | 0,2531 | 0,2109 | 0,2096 | 0,2112 | 0,2534 |
| 0,2312 | 0,1769 | 0,2427 | 0,1828 | 0,2102 | 0,2301 | 0,2557 | 0,2283 | 0,234 | 0,1877 | 0,201 | 0,1813 | 0,1256 | 0,2543 | 0,2168 | 0,2121 | 0,2221 | 0,2546 |
| 0,2254 | 0,1777 | 0,2417 | 0,1867 | 0,2107 | 0,2358 | 0,2434 | 0,2278 | 0,2345 | 0,1899 | 0,2071 | 0,1832 | 0,1235 | 0,2574 | 0,2109 | 0,2143 | 0,2175 | 0,2617 |
| 0,2168 | 0,1803 | 0,2379 | 0,1901 | 0,2117 | 0,2359 | 0,2406 | 0,2137 | 0,2369 | 0,1955 | 0,2036 | 0,1831 | 0,126 | 0,2524 | 0,2212 | 0,2122 | 0,2319 | 0,26 |
| 0,2115 | 0,1814 | 0,241 | 0,1812 | 0,2122 | 0,2369 | 0,2425 | 0,2299 | 0,228 | 0,1998 | 0,2049 | 0,1821 | 0,1223 | 0,2581 | 0,2139 | 0,2135 | 0,232 | 0,2673 |
| 0,2336 | 0,1828 | 0,2541 | 0,1777 | 0,2112 | 0,2394 | 0,2445 | 0,2205 | 0,2272 | 0,192 | 0,208 | 0,18 | 0,1247 | 0,2499 | 0,2198 | 0,2205 | 0,2364 | 0,2704 |
| 0,2195 | 0,1825 | 0,2526 | 0,1761 | 0,2108 | 0,2366 | 0,233 | 0,2272 | 0,2259 | 0,2091 | 0,2091 | 0,1788 | 0,117 | 0,2564 | 0,214 | 0,2169 | 0,2311 | 0,2723 |

| α-Keto-butyric acid | L-asparagine | α-Keto-Glutaric | L-aspartic acic | L-threonine | N-Acetyl-ß-Dn | α-Hydroxy Glu | Propionic Aci | D-trehalose | L-Malic Acid | D-glucosamic | Succinic Acid | Phenylethylar | 2-deoxy-aden | G9 | Glyoxylic Acid | L-Lactic acid | D-Ribose |
|---|---|---|---|---|---|---|---|---|---|---|---|---|---|---|---|---|---|
| 157 | 168 | 159 | 186 | 143 | 139 | 141 | 153 | 150 | 171 | 171 | 168 | 154 | 144 | 125 | 181 | 163 | 564 |
| 154 | 158 | 154 | 172 | 139 | 139 | 135 | 150 | 145 | 162 | 165 | 157 | 152 | 143 | 120 | 170 | 157 | 542 |
| 151 | 158 | 147 | 176 | 135 | 136 | 137 | 142 | 143 | 164 | 160 | 159 | 149 | 145 | 122 | 171 | 155 | 534 |
| 155 | 165 | 150 | 172 | 138 | 135 | 138 | 145 | 140 | 166 | 162 | 161 | 146 | 138 | 122 | 175 | 155 | 542 |
| 153 | 153 | 155 | 177 | 139 | 135 | 139 | 145 | 143 | 166 | 163 | 160 | 151 | 144 | 124 | 174 | 152 | 532 |
| 149 | 155 | 152 | 173 | 139 | 139 | 140 | 145 | 139 | 166 | 163 | 158 | 155 | 143 | 127 | 178 | 154 | 531 |
| 160 | 156 | 154 | 171 | 142 | 137 | 139 | 145 | 139 | 165 | 167 | 160 | 151 | 146 | 123 | 180 | 163 | 532 |
| 157 | 164 | 153 | 168 | 149 | 136 | 143 | 146 | 136 | 169 | 172 | 164 | 156 | 145 | 126 | 182 | 151 | 533 |
| 160 | 169 | 158 | 182 | 151 | 147 | 147 | 146 | 144 | 169 | 170 | 169 | 161 | 144 | 134 | 185 | 163 | 544 |
| 155 | 170 | 156 | 180 | 150 | 146 | 153 | 144 | 143 | 172 | 173 | 174 | 165 | 145 | 134 | 190 | 160 | 562 |
| 159 | 169 | 159 | 183 | 150 | 146 | 154 | 149 | 142 | 171 | 168 | 165 | 164 | 138 | 130 | 188 | 159 | 553 |
| 155 | 175 | 158 | 190 | 156 | 149 | 159 | 150 | 156 | 174 | 170 | 179 | 178 | 140 | 138 | 191 | 169 | 544 |
| 157 | 179 | 173 | 200 | 165 | 156 | 178 | 163 | 159 | 191 | 189 | 177 | 178 | 149 | 147 | 193 | 176 | 547 |
| 164 | 204 | 186 | 214 | 180 | 179 | 184 | 173 | 178 | 198 | 198 | 187 | 187 | 161 | 161 | 207 | 188 | 555 |
| 173 | 208 | 201 | 223 | 196 | 194 | 205 | 185 | 193 | 216 | 209 | 204 | 206 | 173 | 181 | 221 | 201 | 561 |
| 185 | 221 | 213 | 245 | 211 | 213 | 226 | 202 | 202 | 235 | 224 | 222 | 213 | 188 | 200 | 239 | 225 | 571 |
| 204 | 240 | 243 | 271 | 235 | 247 | 264 | 222 | 248 | 273 | 240 | 247 | 228 | 225 | 212 | 252 | 247 | 583 |
| 229 | 295 | 287 | 329 | 280 | 306 | 308 | 275 | 307 | 311 | 271 | 293 | 254 | 237 | 261 | 275 | 301 | 617 |
| 266 | 335 | 327 | 353 | 314 | 348 | 348 | 306 | 353 | 371 | 328 | 341 | 296 | 309 | 325 | 352 | 354 | 662 |
| 320 | 376 | 371 | 405 | 363 | 390 | 389 | 347 | 399 | 422 | 365 | 373 | 325 | 356 | 359 | 400 | 382 | 706 |
| 350 | 440 | 417 | 441 | 397 | 438 | 443 | 412 | 461 | 458 | 419 | 419 | 365 | 406 | 410 | 445 | 422 | 761 |
| 398 | 498 | 451 | 472 | 437 | 478 | 474 | 437 | 507 | 506 | 450 | 458 | 389 | 464 | 459 | 498 | 457 | 800 |
| 460 | 603 | 483 | 518 | 461 | 512 | 500 | 477 | 548 | 573 | 501 | 483 | 406 | 529 | 496 | 568 | 496 | 843 |
| 503 | 642 | 524 | 567 | 480 | 535 | 555 | 509 | 596 | 607 | 516 | 532 | 422 | 584 | 531 | 574 | 528 | 886 |
| 553 | 701 | 565 | 605 | 517 | 588 | 637 | 551 | 645 | 659 | 625 | 571 | 446 | 648 | 574 | 669 | 579 | 941 |
| 589 | 702 | 603 | 645 | 531 | 622 | 620 | 576 | 673 | 715 | 652 | 601 | 455 | 691 | 608 | 646 | 609 | 973 |
| 651 | 721 | 644 | 670 | 563 | 651 | 683 | 621 | 712 | 722 | 699 | 620 | 471 | 751 | 642 | 674 | 634 | 1015 |
| 693 | 744 | 682 | 694 | 603 | 698 | 734 | 653 | 770 | 737 | 746 | 635 | 474 | 821 | 658 | 695 | 685 | 1066 |
| 735 | 743 | 766 | 717 | 636 | 747 | 873 | 709 | 822 | 741 | 771 | 655 | 485 | 881 | 676 | 788 | 718 | 1119 |
| 788 | 733 | 783 | 731 | 673 | 796 | 888 | 738 | 877 | 759 | 777 | 685 | 488 | 951 | 747 | 781 | 769 | 1154 |
| 863 | 736 | 849 | 747 | 705 | 806 | 998 | 777 | 941 | 767 | 834 | 710 | 488 | 1005 | 761 | 816 | 800 | 1203 |
| 927 | 758 | 881 | 766 | 744 | 873 | 986 | 824 | 1007 | 789 | 832 | 762 | 498 | 1042 | 808 | 831 | 864 | 1263 |
| 961 | 775 | 934 | 807 | 790 | 914 | 952 | 853 | 1070 | 849 | 873 | 770 | 507 | 1111 | 869 | 878 | 919 | 1320 |
| 1056 | 778 | 987 | 837 | 838 | 967 | 1034 | 895 | 1108 | 879 | 900 | 806 | 496 | 1143 | 899 | 895 | 1001 | 1399 |
| 1048 | 809 | 1064 | 863 | 889 | 1012 | 1140 | 898 | 1172 | 921 | 949 | 852 | 506 | 1182 | 919 | 935 | 1042 | 1450 |
| 1118 | 846 | 1115 | 891 | 935 | 1028 | 1221 | 982 | 1214 | 940 | 988 | 887 | 521 | 1194 | 895 | 971 | 1074 | 1527 |
| 1148 | 875 | 1169 | 919 | 989 | 1067 | 1200 | 1001 | 1251 | 980 | 1027 | 905 | 522 | 1236 | 951 | 1025 | 1105 | 1598 |
| 1137 | 902 | 1220 | 937 | 1020 | 1122 | 1201 | 1021 | 1272 | 1020 | 1060 | 936 | 517 | 1249 | 962 | 1044 | 1139 | 1659 |
| 1128 | 937 | 1294 | 955 | 1065 | 1184 | 1234 | 1087 | 1283 | 1038 | 1078 | 934 | 513 | 1292 | 981 | 1076 | 1173 | 1701 |
| 1232 | 972 | 1355 | 968 | 1100 | 1196 | 1267 | 1105 | 1247 | 1073 | 1102 | 961 | 497 | 1314 | 1008 | 1113 | 1169 | 1766 |
| 1197 | 1003 | 1428 | 962 | 1123 | 1255 | 1228 | 1172 | 1202 | 1108 | 1120 | 958 | 489 | 1334 | 1052 | 1131 | 1201 | 1834 |
| 6427,6885 | 6369,18383 | 6364,09228 | 6208 | 6198,60848 | 6131,86813 | 6113,6108 | 6101,79641 | 6059,90783 | 6052,97158 | 6040,73838 | 5993,9302 | 5918,72792 | 5908,63952 | 5874,52471 | 5860,57989 | 5821,64891 | 5815,01832 |

| Mucic-acid | Acetic acid | Tween 80 | 1,2-propaned | Adonitol | D-Galactonic Acid-γ-Lactone | Glycolic Acid | L-serine | Myo-inositol | Citric Acid | Methyl Pyruva | Tween 40 | Thymidine | Acetoacetic A | L-proline | Glycyl-LGluta | D-mannose | L-rhamnose | Glucuronami |
|---|---|---|---|---|---|---|---|---|---|---|---|---|---|---|---|---|---|---|
| 0,0555 | 0,0531 | 0,0546 | 0,0511 | 0,0551 | 0,0521 | 0,0581 | 0,0504 | 0,0505 | 0,0531 | 0,053 | 0,058 | 0,0498 | 0,0533 | 0,0528 | 0,0517 | 0,0528 | 0,0531 | 0,0645 |
| 0,0601 | 0,0521 | 0,0552 | 0,0519 | 0,0549 | 0,0532 | 0,0537 | 0,0514 | 0,0513 | 0,0535 | 0,0537 | 0,0587 | 0,0533 | 0,0535 | 0,0541 | 0,0515 | 0,0538 | 0,0582 | 0,055 |
| 0,0656 | 0,0526 | 0,0561 | 0,052 | 0,0614 | 0,0532 | 0,059 | 0,0516 | 0,0517 | 0,0541 | 0,0544 | 0,0584 | 0,0523 | 0,0534 | 0,0538 | 0,0523 | 0,0577 | 0,0625 | 0,0678 |
| 0,0575 | 0,0551 | 0,0556 | 0,0528 | 0,0612 | 0,0537 | 0,0559 | 0,0524 | 0,0524 | 0,0543 | 0,0547 | 0,0584 | 0,0515 | 0,0541 | 0,0573 | 0,0528 | 0,0595 | 0,0576 | 0,0543 |
| 0,0622 | 0,0547 | 0,0561 | 0,0529 | 0,0641 | 0,0543 | 0,0633 | 0,0528 | 0,0528 | 0,0549 | 0,0562 | 0,0581 | 0,0521 | 0,0544 | 0,0565 | 0,0531 | 0,0557 | 0,0559 | 0,0637 |
| 0,0624 | 0,0545 | 0,056 | 0,0536 | 0,0623 | 0,0547 | 0,0609 | 0,053 | 0,0536 | 0,0565 | 0,0562 | 0,0589 | 0,0528 | 0,0554 | 0,0578 | 0,0541 | 0,0576 | 0,0558 | 0,0554 |
| 0,0629 | 0,0558 | 0,0563 | 0,0548 | 0,0555 | 0,0556 | 0,0586 | 0,054 | 0,0542 | 0,0564 | 0,0572 | 0,0589 | 0,0532 | 0,0559 | 0,0617 | 0,0544 | 0,0578 | 0,0578 | 0,071 |
| 0,0665 | 0,0585 | 0,0581 | 0,0563 | 0,0608 | 0,0568 | 0,0633 | 0,0548 | 0,0553 | 0,057 | 0,0584 | 0,0605 | 0,055 | 0,0645 | 0,0607 | 0,0558 | 0,0578 | 0,0609 | 0,055 |
| 0,0606 | 0,0618 | 0,0613 | 0,0588 | 0,0675 | 0,0595 | 0,0655 | 0,0565 | 0,057 | 0,0588 | 0,0601 | 0,0639 | 0,0558 | 0,0575 | 0,065 | 0,0559 | 0,0638 | 0,0627 | 0,0582 |
| 0,0645 | 0,0607 | 0,0638 | 0,0604 | 0,066 | 0,0609 | 0,0602 | 0,0586 | 0,0583 | 0,0597 | 0,0616 | 0,0668 | 0,0566 | 0,059 | 0,0669 | 0,0573 | 0,0649 | 0,0712 | 0,0601 |
| 0,0628 | 0,0664 | 0,0684 | 0,0618 | 0,0683 | 0,0619 | 0,0622 | 0,0597 | 0,0594 | 0,0616 | 0,0628 | 0,0716 | 0,0581 | 0,0616 | 0,0695 | 0,0586 | 0,0685 | 0,0658 | 0,0689 |
| 0,0728 | 0,0664 | 0,0744 | 0,0638 | 0,0672 | 0,0648 | 0,0666 | 0,0615 | 0,0616 | 0,064 | 0,0663 | 0,0785 | 0,0619 | 0,0664 | 0,0743 | 0,0617 | 0,0632 | 0,071 | 0,0738 |
| 0,077 | 0,0747 | 0,0823 | 0,0702 | 0,0719 | 0,0705 | 0,0678 | 0,0667 | 0,0673 | 0,0697 | 0,0722 | 0,086 | 0,0683 | 0,0734 | 0,0779 | 0,0674 | 0,0684 | 0,0751 | 0,0683 |
| 0,0818 | 0,078 | 0,0921 | 0,0786 | 0,0888 | 0,0802 | 0,0769 | 0,0759 | 0,0757 | 0,0777 | 0,0817 | 0,0955 | 0,0788 | 0,0833 | 0,0878 | 0,076 | 0,0801 | 0,0851 | 0,0892 |
| 0,0927 | 0,0977 | 0,1034 | 0,088 | 0,0992 | 0,0909 | 0,0869 | 0,0857 | 0,0847 | 0,0862 | 0,0924 | 0,1068 | 0,0881 | 0,0934 | 0,0992 | 0,0884 | 0,0971 | 0,0915 | 0,0885 |
| 0,1007 | 0,0973 | 0,1134 | 0,0997 | 0,1129 | 0,0977 | 0,1038 | 0,0969 | 0,0979 | 0,0967 | 0,1049 | 0,1154 | 0,1018 | 0,1059 | 0,1109 | 0,1006 | 0,1054 | 0,1029 | 0,1039 |
| 0,1158 | 0,1064 | 0,1248 | 0,112 | 0,1242 | 0,1084 | 0,1193 | 0,1082 | 0,1076 | 0,1064 | 0,1201 | 0,1278 | 0,114 | 0,1327 | 0,121 | 0,1048 | 0,1176 | 0,1133 | 0,1216 |
| 0,1359 | 0,121 | 0,1368 | 0,1253 | 0,1406 | 0,1186 | 0,1427 | 0,1149 | 0,1244 | 0,1203 | 0,1415 | 0,14 | 0,1157 | 0,139 | 0,1287 | 0,1193 | 0,1379 | 0,1333 | 0,148 |
| 0,1456 | 0,1319 | 0,1433 | 0,1375 | 0,148 | 0,1327 | 0,1533 | 0,1298 | 0,1383 | 0,1327 | 0,1573 | 0,1478 | 0,1391 | 0,1465 | 0,1425 | 0,132 | 0,1486 | 0,1484 | 0,1703 |
| 0,1501 | 0,1391 | 0,148 | 0,1476 | 0,1478 | 0,1458 | 0,1457 | 0,1428 | 0,1434 | 0,1378 | 0,1662 | 0,1525 | 0,1465 | 0,1473 | 0,143 | 0,1413 | 0,1613 | 0,1535 | 0,1799 |
| 0,1598 | 0,1526 | 0,1536 | 0,1536 | 0,1539 | 0,1514 | 0,1492 | 0,1566 | 0,1495 | 0,145 | 0,177 | 0,1571 | 0,1488 | 0,1558 | 0,1444 | 0,1541 | 0,1602 | 0,1667 | 0,1865 |
| 0,161 | 0,1532 | 0,1593 | 0,1558 | 0,1584 | 0,1563 | 0,1538 | 0,1644 | 0,1532 | 0,1493 | 0,1885 | 0,1621 | 0,1546 | 0,1638 | 0,1476 | 0,1601 | 0,1698 | 0,171 | 0,1863 |
| 0,1643 | 0,1535 | 0,1639 | 0,163 | 0,1635 | 0,1699 | 0,1659 | 0,1761 | 0,1635 | 0,158 | 0,2024 | 0,1678 | 0,1656 | 0,1719 | 0,1521 | 0,1664 | 0,17 | 0,1792 | 0,2091 |
| 0,1768 | 0,1743 | 0,1716 | 0,1669 | 0,1718 | 0,1762 | 0,1696 | 0,1834 | 0,1642 | 0,1642 | 0,2049 | 0,1715 | 0,1745 | 0,1759 | 0,1579 | 0,1712 | 0,1675 | 0,1894 | 0,2162 |
| 0,1879 | 0,1771 | 0,1768 | 0,1838 | 0,1815 | 0,1851 | 0,179 | 0,2063 | 0,1874 | 0,1687 | 0,2114 | 0,179 | 0,1787 | 0,186 | 0,1608 | 0,1788 | 0,1735 | 0,2014 | 0,2238 |
| 0,1998 | 0,1878 | 0,1834 | 0,1778 | 0,1802 | 0,1879 | 0,1824 | 0,212 | 0,181 | 0,1709 | 0,2124 | 0,1831 | 0,1837 | 0,1817 | 0,1561 | 0,1827 | 0,1908 | 0,2097 | 0,2385 |
| 0,199 | 0,187 | 0,1899 | 0,1756 | 0,1809 | 0,1989 | 0,1727 | 0,2141 | 0,1984 | 0,1739 | 0,2157 | 0,1874 | 0,1896 | 0,1797 | 0,1617 | 0,1911 | 0,1875 | 0,2194 | 0,225 |
| 0,2138 | 0,1834 | 0,1961 | 0,1843 | 0,1844 | 0,2019 | 0,1892 | 0,2167 | 0,1871 | 0,175 | 0,2159 | 0,1917 | 0,1913 | 0,1962 | 0,1629 | 0,196 | 0,2012 | 0,2221 | 0,2185 |
| 0,2316 | 0,1782 | 0,2017 | 0,199 | 0,1885 | 0,2093 | 0,2015 | 0,2232 | 0,2028 | 0,1799 | 0,2187 | 0,1987 | 0,1919 | 0,1942 | 0,1685 | 0,2005 | 0,2012 | 0,243 | 0,2203 |
| 0,2522 | 0,1872 | 0,2064 | 0,2115 | 0,2016 | 0,2197 | 0,2024 | 0,2251 | 0,1935 | 0,1907 | 0,2229 | 0,2033 | 0,198 | 0,2003 | 0,1745 | 0,203 | 0,2094 | 0,2476 | 0,2286 |
| 0,2851 | 0,1873 | 0,2099 | 0,2143 | 0,202 | 0,2215 | 0,2057 | 0,2273 | 0,2006 | 0,1887 | 0,2213 | 0,2065 | 0,1968 | 0,2052 | 0,1744 | 0,2041 | 0,2149 | 0,2568 | 0,2279 |
| 0,266 | 0,1807 | 0,2154 | 0,2051 | 0,1961 | 0,2302 | 0,1959 | 0,2227 | 0,199 | 0,1902 | 0,2244 | 0,2119 | 0,1973 | 0,2005 | 0,1738 | 0,2057 | 0,2205 | 0,257 | 0,2234 |
| 0,2731 | 0,1827 | 0,2189 | 0,2035 | 0,2041 | 0,2363 | 0,1995 | 0,221 | 0,2038 | 0,1901 | 0,2295 | 0,2162 | 0,1974 | 0,2031 | 0,1856 | 0,207 | 0,2167 | 0,2674 | 0,2243 |
| 0,2725 | 0,1843 | 0,2226 | 0,2056 | 0,2109 | 0,242 | 0,2078 | 0,2263 | 0,2032 | 0,1949 | 0,2306 | 0,2208 | 0,1961 | 0,212 | 0,1865 | 0,2093 | 0,2292 | 0,2726 | 0,2311 |
| 0,2794 | 0,19 | 0,2256 | 0,2061 | 0,2127 | 0,2483 | 0,2128 | 0,2294 | 0,2071 | 0,1962 | 0,231 | 0,224 | 0,1973 | 0,2099 | 0,1902 | 0,2111 | 0,2239 | 0,2827 | 0,2334 |
| 0,2858 | 0,1901 | 0,2292 | 0,2045 | 0,2213 | 0,2512 | 0,2314 | 0,2298 | 0,2124 | 0,1987 | 0,2321 | 0,2273 | 0,197 | 0,2191 | 0,1865 | 0,2132 | 0,2266 | 0,2948 | 0,2337 |
| 0,2746 | 0,2024 | 0,2323 | 0,2063 | 0,2148 | 0,2643 | 0,219 | 0,2319 | 0,2142 | 0,2003 | 0,2358 | 0,2279 | 0,1954 | 0,2165 | 0,1882 | 0,2139 | 0,2186 | 0,2912 | 0,2263 |
| 0,2776 | 0,1982 | 0,2324 | 0,2065 | 0,21 | 0,2634 | 0,2286 | 0,2331 | 0,2133 | 0,2016 | 0,236 | 0,2346 | 0,1952 | 0,2146 | 0,1846 | 0,2176 | 0,2161 | 0,3007 | 0,2342 |
| 0,2831 | 0,2043 | 0,2345 | 0,2055 | 0,2174 | 0,2653 | 0,2255 | 0,2352 | 0,215 | 0,2016 | 0,2353 | 0,2345 | 0,189 | 0,2116 | 0,1873 | 0,2179 | 0,2092 | 0,3066 | 0,2334 |
| 0,2795 | 0,2056 | 0,2358 | 0,2069 | 0,2189 | 0,2657 | 0,2282 | 0,2366 | 0,2184 | 0,2016 | 0,2369 | 0,235 | 0,1823 | 0,2204 | 0,1709 | 0,2193 | 0,205 | 0,3172 | 0,2422 |
| 0,2814 | 0,2143 | 0,2355 | 0,2103 | 0,2262 | 0,2659 | 0,2303 | 0,2377 | 0,2169 | 0,2014 | 0,239 | 0,2359 | 0,1786 | 0,2176 | 0,167 | 0,2209 | 0,2007 | 0,3275 | 0,2376 |

| Mucic-acid | Acetic acid | Tween 80 | 1,2-propaned | Adonitol | D-Galactonic Acid-γ-Lactone | Glycolic Acid | L-serine | Myo-inositol | Citric Acid | Methyl Pyruva | Tween 40 | Thymidine | Acetoacetic A | L-proline | Glycyl-LGluta | D-mannose | L-rhamnose | Glucuronami |
|---|---|---|---|---|---|---|---|---|---|---|---|---|---|---|---|---|---|---|
| 165 | 141 | 155 | 172 | 148 | 176 | 168 | 165 | 165 | 172 | 163 | 150 | 127 | 221 | 157 | 163 | 148 | 153 | 234 |
| 160 | 137 | 150 | 166 | 140 | 175 | 158 | 156 | 161 | 166 | 160 | 146 | 124 | 215 | 145 | 159 | 143 | 147 | 230 |
| 157 | 137 | 147 | 166 | 141 | 170 | 159 | 155 | 163 | 162 | 159 | 139 | 123 | 210 | 148 | 154 | 146 | 148 | 220 |
| 154 | 141 | 145 | 158 | 139 | 173 | 161 | 155 | 163 | 166 | 158 | 139 | 120 | 203 | 149 | 154 | 142 | 145 | 223 |
| 160 | 141 | 147 | 159 | 139 | 167 | 159 | 156 | 163 | 162 | 162 | 137 | 119 | 207 | 146 | 152 | 139 | 143 | 224 |
| 161 | 140 | 147 | 162 | 141 | 171 | 158 | 158 | 164 | 163 | 160 | 136 | 125 | 207 | 148 | 155 | 132 | 146 | 220 |
| 158 | 142 | 145 | 167 | 146 | 170 | 159 | 158 | 164 | 170 | 162 | 140 | 123 | 209 | 147 | 158 | 136 | 151 | 219 |
| 157 | 137 | 147 | 165 | 144 | 174 | 163 | 162 | 165 | 162 | 169 | 135 | 123 | 206 | 150 | 157 | 137 | 152 | 212 |
| 162 | 141 | 160 | 171 | 153 | 176 | 166 | 162 | 171 | 171 | 172 | 146 | 128 | 211 | 161 | 160 | 140 | 157 | 228 |
| 157 | 144 | 165 | 174 | 155 | 172 | 172 | 168 | 172 | 173 | 165 | 151 | 122 | 211 | 162 | 158 | 135 | 160 | 220 |
| 170 | 139 | 170 | 170 | 152 | 173 | 171 | 165 | 174 | 168 | 170 | 158 | 126 | 211 | 165 | 154 | 127 | 158 | 225 |
| 174 | 144 | 179 | 175 | 155 | 182 | 176 | 173 | 172 | 176 | 176 | 170 | 125 | 214 | 167 | 160 | 144 | 167 | 222 |
| 182 | 155 | 190 | 191 | 172 | 192 | 189 | 176 | 186 | 185 | 193 | 180 | 130 | 222 | 173 | 168 | 147 | 170 | 235 |
| 196 | 162 | 204 | 202 | 190 | 208 | 202 | 194 | 200 | 196 | 205 | 199 | 147 | 235 | 185 | 173 | 154 | 188 | 246 |
| 212 | 177 | 221 | 214 | 209 | 224 | 218 | 208 | 212 | 216 | 227 | 210 | 154 | 251 | 202 | 194 | 174 | 198 | 261 |
| 228 | 198 | 242 | 231 | 229 | 228 | 243 | 226 | 230 | 230 | 248 | 235 | 165 | 253 | 222 | 206 | 192 | 226 | 280 |
| 255 | 198 | 270 | 245 | 260 | 237 | 264 | 248 | 253 | 246 | 268 | 263 | 205 | 285 | 253 | 196 | 214 | 228 | 275 |
| 321 | 243 | 320 | 294 | 319 | 259 | 320 | 264 | 296 | 276 | 316 | 304 | 232 | 314 | 309 | 234 | 258 | 294 | 329 |
| 366 | 299 | 352 | 331 | 360 | 323 | 360 | 310 | 343 | 333 | 372 | 335 | 299 | 351 | 349 | 281 | 312 | 336 | 388 |
| 402 | 344 | 395 | 384 | 411 | 368 | 399 | 352 | 377 | 371 | 419 | 383 | 364 | 397 | 402 | 311 | 366 | 368 | 435 |
| 441 | 381 | 428 | 426 | 446 | 403 | 434 | 391 | 426 | 414 | 470 | 425 | 444 | 431 | 430 | 361 | 434 | 405 | 505 |
| 479 | 423 | 470 | 454 | 489 | 450 | 463 | 440 | 470 | 463 | 543 | 449 | 502 | 462 | 481 | 397 | 493 | 432 | 574 |
| 513 | 478 | 514 | 491 | 524 | 482 | 484 | 490 | 534 | 493 | 583 | 490 | 572 | 502 | 498 | 429 | 534 | 481 | 624 |
| 546 | 511 | 552 | 519 | 550 | 537 | 507 | 526 | 543 | 526 | 626 | 523 | 627 | 535 | 528 | 466 | 580 | 515 | 699 |
| 582 | 559 | 596 | 616 | 612 | 580 | 545 | 631 | 632 | 578 | 684 | 551 | 645 | 574 | 552 | 503 | 619 | 573 | 741 |
| 630 | 594 | 644 | 614 | 610 | 622 | 564 | 681 | 638 | 586 | 705 | 595 | 678 | 625 | 552 | 531 | 654 | 604 | 785 |
| 639 | 610 | 686 | 632 | 630 | 665 | 575 | 742 | 707 | 642 | 746 | 645 | 692 | 662 | 560 | 564 | 690 | 680 | 798 |
| 700 | 651 | 731 | 657 | 664 | 704 | 619 | 789 | 686 | 673 | 766 | 682 | 702 | 696 | 585 | 596 | 743 | 713 | 835 |
| 752 | 687 | 783 | 751 | 708 | 742 | 662 | 840 | 758 | 711 | 801 | 719 | 727 | 756 | 580 | 635 | 775 | 766 | 859 |
| 841 | 718 | 833 | 775 | 752 | 788 | 681 | 861 | 741 | 734 | 842 | 758 | 744 | 765 | 603 | 634 | 839 | 817 | 882 |
| 963 | 735 | 883 | 840 | 783 | 822 | 716 | 893 | 809 | 757 | 848 | 799 | 761 | 814 | 605 | 662 | 868 | 862 | 886 |
| 997 | 767 | 918 | 794 | 798 | 896 | 716 | 910 | 833 | 799 | 887 | 828 | 805 | 848 | 624 | 690 | 924 | 930 | 898 |
| 1082 | 815 | 983 | 839 | 850 | 944 | 756 | 954 | 853 | 808 | 939 | 872 | 804 | 879 | 644 | 733 | 977 | 983 | 905 |
| 1149 | 824 | 1021 | 869 | 881 | 987 | 779 | 964 | 886 | 840 | 966 | 900 | 824 | 913 | 657 | 763 | 1006 | 1043 | 921 |
| 1242 | 856 | 1062 | 884 | 910 | 1019 | 822 | 1001 | 906 | 875 | 982 | 943 | 834 | 944 | 676 | 793 | 1055 | 1105 | 943 |
| 1283 | 885 | 1066 | 920 | 967 | 1065 | 903 | 1019 | 969 | 891 | 988 | 965 | 846 | 945 | 690 | 832 | 1030 | 1175 | 954 |
| 1298 | 900 | 1100 | 953 | 949 | 1121 | 890 | 1035 | 962 | 906 | 1023 | 1007 | 839 | 986 | 709 | 855 | 1025 | 1237 | 981 |
| 1342 | 949 | 1116 | 980 | 972 | 1173 | 935 | 1056 | 992 | 908 | 1042 | 1019 | 832 | 977 | 720 | 895 | 997 | 1301 | 1009 |
| 1393 | 956 | 1138 | 985 | 1003 | 1219 | 1014 | 1077 | 1019 | 933 | 1057 | 1036 | 813 | 1021 | 728 | 910 | 954 | 1357 | 1016 |
| 1420 | 988 | 1130 | 1026 | 1034 | 1274 | 998 | 1104 | 1022 | 945 | 1091 | 1050 | 794 | 1029 | 723 | 952 | 915 | 1427 | 1026 |
| 1456 | 1023 | 1140 | 1033 | 1061 | 1305 | 1075 | 1150 | 1034 | 944 | 1120 | 1063 | 775 | 1043 | 727 | 986 | 857 | 1461 | 1049 |
| 5714,91811 | 5471,46402 | 5444,99724 | 5408,29146 | 5336,06078 | 5280,63611 | 5267,13124 | 5258,94287 | 5222,35577 | 5205,66419 | 5145,16129 | 5132,09668 | 5031,0559 | 5003,04321 | 4991,24343 | 4864,06619 | 4793,77958 | 4766,76385 | 4708,26112 |

| Tween 20 | L-Lyxose | Tyramine | D-fructose-6-p | D-Serine | Glycyl-L-Proli | Tricarballylic | Sucrose | D-sorbitol | Glycerol | Pyruvic Acid | D-aspartic aci | glucose-6-pho | D-gluconic aci | D-gluconique | Methyl D gala | L-Galactonic | M-tartaric aci |
|---|---|---|---|---|---|---|---|---|---|---|---|---|---|---|---|---|---|
| 0,0532 | 0,0577 | 0,0546 | 0,0521 | 0,0499 | 0,0519 | 0,0507 | 0,052 | 0,0503 | 0,0512 | 0,0545 | 0,0528 | 0,051 | 0,0539 | 0,0501 | 0,0623 | 0,058 | 0,0518 |
| 0,0537 | 0,0586 | 0,0548 | 0,0516 | 0,0509 | 0,0518 | 0,0518 | 0,0549 | 0,0515 | 0,0529 | 0,0562 | 0,0532 | 0,0522 | 0,0555 | 0,0506 | 0,0597 | 0,0564 | 0,052 |
| 0,054 | 0,0603 | 0,056 | 0,0516 | 0,0518 | 0,0521 | 0,0517 | 0,0607 | 0,0515 | 0,0534 | 0,0557 | 0,0532 | 0,0529 | 0,0537 | 0,0516 | 0,066 | 0,0627 | 0,0518 |
| 0,0542 | 0,0673 | 0,0587 | 0,0519 | 0,0517 | 0,053 | 0,0524 | 0,0564 | 0,0521 | 0,0535 | 0,0547 | 0,0539 | 0,0528 | 0,0536 | 0,0518 | 0,0593 | 0,0663 | 0,0521 |
| 0,0547 | 0,0655 | 0,0583 | 0,0519 | 0,0521 | 0,0532 | 0,053 | 0,0578 | 0,0523 | 0,054 | 0,0556 | 0,0545 | 0,0537 | 0,0544 | 0,0531 | 0,0616 | 0,0606 | 0,0526 |
| 0,0552 | 0,0712 | 0,057 | 0,0525 | 0,0524 | 0,0541 | 0,0532 | 0,0538 | 0,0531 | 0,0545 | 0,0575 | 0,0554 | 0,0541 | 0,0584 | 0,0532 | 0,0539 | 0,0626 | 0,0528 |
| 0,0559 | 0,0572 | 0,0618 | 0,0524 | 0,0537 | 0,0548 | 0,0539 | 0,055 | 0,0532 | 0,055 | 0,0575 | 0,0567 | 0,0553 | 0,0611 | 0,0531 | 0,0591 | 0,0609 | 0,0538 |
| 0,0571 | 0,0586 | 0,0592 | 0,0529 | 0,0547 | 0,0557 | 0,0554 | 0,0551 | 0,0555 | 0,0574 | 0,0571 | 0,0576 | 0,0576 | 0,059 | 0,0552 | 0,06 | 0,0611 | 0,0553 |
| 0,0599 | 0,0711 | 0,0597 | 0,054 | 0,0562 | 0,0575 | 0,0567 | 0,0557 | 0,0594 | 0,0591 | 0,0614 | 0,0597 | 0,0624 | 0,0638 | 0,057 | 0,0592 | 0,0629 | 0,0571 |
| 0,0631 | 0,0651 | 0,0632 | 0,0556 | 0,0577 | 0,0582 | 0,0581 | 0,0622 | 0,0637 | 0,0615 | 0,0604 | 0,0611 | 0,0686 | 0,0649 | 0,0586 | 0,0609 | 0,063 | 0,0577 |
| 0,0685 | 0,0601 | 0,0631 | 0,0581 | 0,0591 | 0,0593 | 0,0592 | 0,0595 | 0,0655 | 0,0627 | 0,0738 | 0,0618 | 0,0747 | 0,0705 | 0,06 | 0,0726 | 0,0681 | 0,0584 |
| 0,076 | 0,0713 | 0,0618 | 0,062 | 0,0624 | 0,0619 | 0,0615 | 0,0655 | 0,069 | 0,0673 | 0,0645 | 0,0646 | 0,0826 | 0,0768 | 0,0635 | 0,0644 | 0,0654 | 0,06 |
| 0,0864 | 0,0667 | 0,0677 | 0,0644 | 0,0676 | 0,0672 | 0,0663 | 0,069 | 0,0768 | 0,0741 | 0,0708 | 0,0702 | 0,0929 | 0,0854 | 0,0691 | 0,0712 | 0,069 | 0,0631 |
| 0,0975 | 0,0743 | 0,0701 | 0,0679 | 0,0733 | 0,0759 | 0,0739 | 0,0738 | 0,087 | 0,0842 | 0,0886 | 0,0779 | 0,1151 | 0,0972 | 0,0776 | 0,083 | 0,0808 | 0,0693 |
| 0,1091 | 0,0963 | 0,0806 | 0,076 | 0,0815 | 0,0872 | 0,084 | 0,0803 | 0,0975 | 0,0951 | 0,0973 | 0,0878 | 0,1324 | 0,1091 | 0,0895 | 0,0969 | 0,0923 | 0,0772 |
| 0,119 | 0,0994 | 0,091 | 0,0915 | 0,0873 | 0,098 | 0,0934 | 0,0944 | 0,1096 | 0,105 | 0,1124 | 0,0988 | 0,1499 | 0,1218 | 0,1003 | 0,1013 | 0,1045 | 0,087 |
| 0,1264 | 0,1118 | 0,1015 | 0,108 | 0,0927 | 0,1042 | 0,1024 | 0,0993 | 0,1106 | 0,1054 | 0,1354 | 0,1079 | 0,1842 | 0,1393 | 0,1076 | 0,1194 | 0,1237 | 0,092 |
| 0,1406 | 0,132 | 0,1106 | 0,1285 | 0,1013 | 0,1178 | 0,1152 | 0,1111 | 0,1399 | 0,1331 | 0,1577 | 0,124 | 0,198 | 0,1713 | 0,1296 | 0,1359 | 0,1463 | 0,1038 |
| 0,1517 | 0,1487 | 0,1233 | 0,1495 | 0,1128 | 0,1291 | 0,1301 | 0,1306 | 0,1528 | 0,1411 | 0,1785 | 0,1323 | 0,2378 | 0,1914 | 0,1522 | 0,1451 | 0,1694 | 0,1162 |
| 0,1551 | 0,1563 | 0,1375 | 0,1674 | 0,1219 | 0,1408 | 0,1342 | 0,1404 | 0,1619 | 0,1544 | 0,1966 | 0,1324 | 0,261 | 0,2307 | 0,1721 | 0,1616 | 0,1891 | 0,1261 |
| 0,1595 | 0,1642 | 0,1497 | 0,1993 | 0,1318 | 0,1511 | 0,1438 | 0,1489 | 0,174 | 0,1628 | 0,2204 | 0,1333 | 0,2729 | 0,2351 | 0,1969 | 0,1657 | 0,2136 | 0,1408 |
| 0,1649 | 0,1623 | 0,1597 | 0,2388 | 0,1363 | 0,1576 | 0,1448 | 0,1435 | 0,1825 | 0,1665 | 0,2433 | 0,136 | 0,2532 | 0,2347 | 0,223 | 0,1734 | 0,2394 | 0,1439 |
| 0,1692 | 0,1643 | 0,1679 | 0,2524 | 0,1398 | 0,1618 | 0,1537 | 0,1585 | 0,2033 | 0,1755 | 0,2465 | 0,1398 | 0,2566 | 0,2483 | 0,2493 | 0,1917 | 0,2402 | 0,1555 |
| 0,1742 | 0,1881 | 0,1744 | 0,2572 | 0,1459 | 0,1657 | 0,1618 | 0,1603 | 0,2047 | 0,1857 | 0,2619 | 0,1424 | 0,2542 | 0,2428 | 0,2393 | 0,1975 | 0,2547 | 0,146 |
| 0,1802 | 0,1853 | 0,1757 | 0,2788 | 0,154 | 0,1665 | 0,166 | 0,1613 | 0,2141 | 0,1919 | 0,2615 | 0,1464 | 0,26 | 0,2455 | 0,2591 | 0,2101 | 0,2515 | 0,1484 |
| 0,184 | 0,1789 | 0,1759 | 0,2732 | 0,1625 | 0,1685 | 0,1681 | 0,1609 | 0,2186 | 0,2069 | 0,2634 | 0,148 | 0,2358 | 0,25 | 0,2766 | 0,2264 | 0,2582 | 0,148 |
| 0,1888 | 0,1939 | 0,1777 | 0,2883 | 0,1761 | 0,17 | 0,1719 | 0,1614 | 0,2194 | 0,2084 | 0,2594 | 0,1578 | 0,2366 | 0,2441 | 0,2479 | 0,2275 | 0,2494 | 0,1475 |
| 0,1939 | 0,1942 | 0,1812 | 0,3048 | 0,179 | 0,1698 | 0,1744 | 0,1676 | 0,2236 | 0,2117 | 0,2577 | 0,1542 | 0,2405 | 0,2417 | 0,2509 | 0,2386 | 0,2462 | 0,1472 |
| 0,1974 | 0,1964 | 0,1819 | 0,3258 | 0,1858 | 0,1715 | 0,1786 | 0,1673 | 0,2332 | 0,2176 | 0,2606 | 0,1597 | 0,238 | 0,2427 | 0,2478 | 0,2499 | 0,2459 | 0,1495 |
| 0,2022 | 0,2001 | 0,1859 | 0,3312 | 0,1929 | 0,1723 | 0,1826 | 0,1688 | 0,242 | 0,2209 | 0,2569 | 0,1584 | 0,2412 | 0,2417 | 0,2475 | 0,2652 | 0,2465 | 0,1541 |
| 0,2064 | 0,1926 | 0,1855 | 0,3013 | 0,1961 | 0,1743 | 0,1855 | 0,1841 | 0,2453 | 0,2279 | 0,2536 | 0,1624 | 0,2464 | 0,2405 | 0,2451 | 0,2683 | 0,2441 | 0,1568 |
| 0,2105 | 0,1937 | 0,189 | 0,2911 | 0,1987 | 0,1753 | 0,1855 | 0,1788 | 0,2525 | 0,233 | 0,2503 | 0,1623 | 0,2481 | 0,2411 | 0,2417 | 0,2719 | 0,2545 | 0,1617 |
| 0,2169 | 0,1966 | 0,1898 | 0,2902 | 0,2066 | 0,1753 | 0,1873 | 0,1817 | 0,2613 | 0,2388 | 0,253 | 0,1633 | 0,2448 | 0,241 | 0,2432 | 0,2844 | 0,2524 | 0,1663 |
| 0,2186 | 0,2026 | 0,1881 | 0,281 | 0,2081 | 0,1762 | 0,1906 | 0,1806 | 0,2675 | 0,2437 | 0,2508 | 0,1645 | 0,2478 | 0,2405 | 0,2473 | 0,2908 | 0,2513 | 0,1701 |
| 0,2208 | 0,1964 | 0,1911 | 0,2876 | 0,2021 | 0,1766 | 0,1945 | 0,1886 | 0,2715 | 0,2498 | 0,2501 | 0,167 | 0,2485 | 0,238 | 0,2411 | 0,2965 | 0,2522 | 0,1777 |
| 0,2248 | 0,2042 | 0,1933 | 0,2971 | 0,1978 | 0,1771 | 0,1975 | 0,1917 | 0,2775 | 0,2514 | 0,256 | 0,1676 | 0,248 | 0,2433 | 0,2435 | 0,3046 | 0,2578 | 0,1817 |
| 0,2259 | 0,2067 | 0,1962 | 0,2914 | 0,2075 | 0,1774 | 0,1976 | 0,192 | 0,2822 | 0,2572 | 0,2566 | 0,1691 | 0,2489 | 0,2459 | 0,2464 | 0,3006 | 0,2522 | 0,1845 |
| 0,2279 | 0,1964 | 0,1934 | 0,2858 | 0,2121 | 0,1771 | 0,1975 | 0,1907 | 0,2868 | 0,2646 | 0,2573 | 0,1716 | 0,2498 | 0,2473 | 0,251 | 0,2978 | 0,2488 | 0,1868 |
| 0,2318 | 0,1948 | 0,196 | 0,2873 | 0,203 | 0,1772 | 0,1996 | 0,196 | 0,2911 | 0,2694 | 0,2602 | 0,1743 | 0,2496 | 0,2499 | 0,2516 | 0,3061 | 0,249 | 0,1893 |
| 0,2336 | 0,2003 | 0,1939 | 0,2786 | 0,2 | 0,1752 | 0,1996 | 0,1939 | 0,2966 | 0,2752 | 0,2589 | 0,1746 | 0,2491 | 0,2528 | 0,2526 | 0,3082 | 0,2533 | 0,1912 |
| 0,2317 | 0,1998 | 0,1948 | 0,2819 | 0,1844 | 0,1721 | 0,2001 | 0,1983 | 0,2984 | 0,2791 | 0,2585 | 0,1758 | 0,2492 | 0,2571 | 0,2557 | 0,3183 | 0,264 | 0,1943 |

| Tween 20 | L-Lyxose | Tyramine | D-fructose-6-p | D-Serine | Glycyl-L-Proli | Tricarballylic | Sucrose | D-sorbitol | Glycerol | Pyruvic Acid | D-aspartic aci | glucose-6-pho | D-gluconic aci | D-gluconique | Methyl D gala | L-Galactonic | M-tartaric aci |
|---|---|---|---|---|---|---|---|---|---|---|---|---|---|---|---|---|---|
| 140 | 211 | 148 | 194 | 177 | 136 | 150 | 142 | 154 | 200 | 190 | 177 | 176 | 176 | 207 | 132 | 183 | 176 |
| 136 | 206 | 142 | 190 | 167 | 132 | 145 | 139 | 147 | 189 | 184 | 171 | 171 | 174 | 199 | 127 | 175 | 169 |
| 134 | 201 | 142 | 188 | 167 | 133 | 144 | 144 | 142 | 190 | 183 | 166 | 169 | 171 | 200 | 130 | 168 | 167 |
| 137 | 210 | 142 | 186 | 161 | 136 | 146 | 137 | 151 | 194 | 179 | 168 | 164 | 169 | 199 | 127 | 176 | 162 |
| 135 | 211 | 147 | 192 | 161 | 136 | 147 | 136 | 148 | 187 | 180 | 165 | 163 | 167 | 195 | 124 | 171 | 167 |
| 141 | 216 | 144 | 190 | 166 | 138 | 146 | 138 | 145 | 193 | 187 | 165 | 167 | 167 | 197 | 127 | 179 | 170 |
| 139 | 208 | 142 | 190 | 165 | 136 | 147 | 143 | 144 | 197 | 177 | 166 | 165 | 169 | 199 | 129 | 173 | 166 |
| 137 | 210 | 146 | 188 | 170 | 140 | 150 | 141 | 145 | 188 | 187 | 170 | 162 | 162 | 199 | 129 | 174 | 169 |
| 144 | 215 | 152 | 191 | 171 | 142 | 155 | 148 | 146 | 196 | 186 | 170 | 166 | 168 | 203 | 138 | 176 | 172 |
| 149 | 213 | 154 | 194 | 166 | 145 | 154 | 149 | 145 | 186 | 183 | 175 | 162 | 156 | 203 | 135 | 170 | 172 |
| 158 | 218 | 157 | 193 | 168 | 142 | 157 | 147 | 139 | 185 | 182 | 176 | 157 | 150 | 205 | 133 | 172 | 174 |
| 166 | 211 | 161 | 191 | 170 | 148 | 158 | 149 | 137 | 185 | 176 | 180 | 155 | 158 | 209 | 141 | 173 | 170 |
| 181 | 225 | 158 | 187 | 186 | 149 | 171 | 159 | 145 | 191 | 185 | 190 | 155 | 161 | 220 | 146 | 176 | 175 |
| 196 | 245 | 171 | 182 | 197 | 159 | 183 | 167 | 161 | 201 | 192 | 205 | 158 | 171 | 228 | 160 | 186 | 184 |
| 214 | 253 | 179 | 181 | 216 | 173 | 194 | 182 | 179 | 202 | 202 | 220 | 167 | 179 | 238 | 172 | 199 | 195 |
| 239 | 264 | 192 | 188 | 217 | 178 | 211 | 197 | 193 | 222 | 208 | 234 | 174 | 187 | 255 | 185 | 202 | 213 |
| 260 | 286 | 205 | 192 | 233 | 183 | 226 | 201 | 209 | 227 | 241 | 255 | 186 | 211 | 266 | 234 | 234 | 219 |
| 294 | 331 | 217 | 206 | 251 | 208 | 249 | 246 | 258 | 268 | 255 | 296 | 212 | 244 | 286 | 272 | 249 | 223 |
| 343 | 389 | 248 | 218 | 258 | 239 | 312 | 303 | 305 | 305 | 297 | 324 | 240 | 298 | 319 | 311 | 284 | 259 |
| 386 | 422 | 296 | 230 | 327 | 268 | 337 | 340 | 339 | 346 | 342 | 355 | 293 | 352 | 343 | 364 | 313 | 306 |
| 428 | 459 | 329 | 273 | 364 | 295 | 375 | 377 | 392 | 391 | 430 | 376 | 371 | 407 | 399 | 407 | 391 | 342 |
| 459 | 507 | 370 | 349 | 415 | 334 | 414 | 429 | 425 | 418 | 496 | 393 | 436 | 465 | 465 | 432 | 470 | 379 |
| 492 | 556 | 400 | 434 | 453 | 354 | 447 | 474 | 469 | 448 | 549 | 412 | 499 | 496 | 524 | 480 | 525 | 409 |
| 519 | 583 | 425 | 510 | 478 | 374 | 482 | 502 | 503 | 476 | 589 | 433 | 544 | 531 | 559 | 500 | 581 | 404 |
| 553 | 623 | 466 | 606 | 511 | 388 | 514 | 531 | 556 | 514 | 647 | 456 | 602 | 563 | 630 | 566 | 629 | 425 |
| 598 | 643 | 482 | 650 | 549 | 416 | 525 | 563 | 600 | 542 | 688 | 470 | 628 | 612 | 691 | 611 | 666 | 434 |
| 645 | 653 | 502 | 749 | 573 | 420 | 565 | 581 | 630 | 572 | 730 | 479 | 635 | 636 | 713 | 647 | 692 | 439 |
| 671 | 685 | 525 | 818 | 597 | 436 | 583 | 626 | 662 | 596 | 777 | 508 | 665 | 663 | 754 | 704 | 724 | 441 |
| 705 | 709 | 550 | 871 | 633 | 466 | 618 | 640 | 711 | 635 | 795 | 518 | 690 | 688 | 775 | 760 | 759 | 448 |
| 739 | 718 | 574 | 914 | 676 | 492 | 636 | 633 | 761 | 660 | 802 | 516 | 722 | 729 | 808 | 821 | 772 | 455 |
| 770 | 727 | 590 | 938 | 702 | 501 | 645 | 645 | 817 | 697 | 822 | 526 | 747 | 740 | 828 | 878 | 783 | 463 |
| 810 | 739 | 633 | 943 | 734 | 531 | 664 | 673 | 859 | 736 | 843 | 543 | 774 | 764 | 840 | 927 | 786 | 489 |
| 841 | 762 | 644 | 996 | 752 | 544 | 695 | 683 | 919 | 782 | 844 | 552 | 813 | 786 | 868 | 987 | 795 | 502 |
| 876 | 793 | 675 | 1007 | 769 | 566 | 705 | 707 | 958 | 812 | 858 | 557 | 839 | 803 | 877 | 1034 | 811 | 523 |
| 892 | 803 | 706 | 1033 | 779 | 584 | 717 | 721 | 999 | 850 | 874 | 575 | 860 | 819 | 894 | 1071 | 830 | 543 |
| 920 | 813 | 736 | 1122 | 796 | 614 | 752 | 732 | 1048 | 889 | 895 | 591 | 888 | 838 | 897 | 1109 | 847 | 557 |
| 933 | 836 | 744 | 1118 | 794 | 631 | 754 | 745 | 1091 | 947 | 922 | 604 | 906 | 858 | 911 | 1101 | 868 | 583 |
| 958 | 849 | 769 | 1119 | 803 | 643 | 764 | 748 | 1101 | 975 | 925 | 609 | 927 | 873 | 948 | 1109 | 890 | 599 |
| 948 | 847 | 776 | 1153 | 797 | 653 | 768 | 752 | 1122 | 1017 | 946 | 631 | 935 | 894 | 953 | 1113 | 889 | 622 |
| 963 | 853 | 768 | 1155 | 792 | 664 | 769 | 744 | 1141 | 1063 | 964 | 642 | 943 | 913 | 969 | 1093 | 897 | 639 |
| 963 | 857 | 783 | 1194 | 762 | 657 | 779 | 742 | 1157 | 1096 | 989 | 657 | 941 | 951 | 991 | 1063 | 917 | 668 |
| 4610,64426 | 4546,0943 | 4529,24394 | 4351,6101 | 4349,44238 | 4334,4426 | 4210,17403 | 4101,162 | 4042,72471 | 3931,54892 | 3916,66667 | 3902,43902 | 3859,73764 | 3813,97638 | 3813,22957 | 3636,71875 | 3563,1068 | 3452,63158 |

| L-glutamine | m-Hydroxy Ph | 2-Aminoethar | D-saccharide a | L-glutamic aci | D-Psicose | L-fucose | Phenyl Acetic | Formic acid | D-melibiose | Lactulose | MOPS no sugar | p-Hydroxy Phenyl Acetic Acid | Dulcitol | D-mannitol | D-galactose |
|---|---|---|---|---|---|---|---|---|---|---|---|---|---|---|---|
| 0,0486 | 0,0532 | 0,0595 | 0,0504 | 0,0532 | 0,0563 | 0,0503 | 0,054 | 0,053 | 0,0506 | 0,0521 | 0,0488 | 0,0542 | 0,0513 | 0,0563 | 0,0536 |
| 0,0487 | 0,0533 | 0,0577 | 0,0503 | 0,0529 | 0,0603 | 0,0513 | 0,0554 | 0,0544 | 0,0515 | 0,0531 | 0,0501 | 0,055 | 0,0536 | 0,0555 | 0,0543 |
| 0,0488 | 0,0539 | 0,0607 | 0,0514 | 0,0541 | 0,0606 | 0,0514 | 0,0556 | 0,0556 | 0,0528 | 0,053 | 0,0505 | 0,0555 | 0,0514 | 0,0563 | 0,0552 |
| 0,0496 | 0,0542 | 0,0664 | 0,0534 | 0,054 | 0,0587 | 0,0525 | 0,056 | 0,0575 | 0,0532 | 0,0533 | 0,0516 | 0,0555 | 0,0525 | 0,0553 | 0,0578 |
| 0,0503 | 0,0543 | 0,0581 | 0,0545 | 0,0547 | 0,0586 | 0,0529 | 0,0562 | 0,0587 | 0,0529 | 0,0529 | 0,0516 | 0,0561 | 0,0528 | 0,0594 | 0,057 |
| 0,0508 | 0,0552 | 0,0559 | 0,0559 | 0,056 | 0,0634 | 0,0535 | 0,0571 | 0,0558 | 0,0532 | 0,0545 | 0,0524 | 0,0569 | 0,0532 | 0,0556 | 0,0594 |
| 0,0515 | 0,0552 | 0,0541 | 0,0559 | 0,0564 | 0,063 | 0,0537 | 0,0573 | 0,0564 | 0,0536 | 0,0553 | 0,053 | 0,0571 | 0,0535 | 0,0574 | 0,0579 |
| 0,0523 | 0,0557 | 0,0562 | 0,0574 | 0,0556 | 0,0684 | 0,056 | 0,0588 | 0,058 | 0,0562 | 0,057 | 0,0546 | 0,0585 | 0,0548 | 0,0574 | 0,0599 |
| 0,0532 | 0,057 | 0,0579 | 0,06 | 0,0575 | 0,0663 | 0,0593 | 0,0607 | 0,0596 | 0,0605 | 0,0602 | 0,0563 | 0,0606 | 0,058 | 0,0611 | 0,0618 |
| 0,0546 | 0,0582 | 0,0601 | 0,0608 | 0,0591 | 0,0683 | 0,0618 | 0,0618 | 0,0636 | 0,0596 | 0,0636 | 0,0576 | 0,0619 | 0,064 | 0,0616 | 0,069 |
| 0,0555 | 0,0589 | 0,0608 | 0,0624 | 0,0624 | 0,0622 | 0,0641 | 0,0633 | 0,0616 | 0,0636 | 0,0606 | 0,0591 | 0,0629 | 0,0617 | 0,0639 | 0,0695 |
| 0,0582 | 0,0603 | 0,0742 | 0,0664 | 0,0634 | 0,0637 | 0,0665 | 0,0645 | 0,0631 | 0,063 | 0,0638 | 0,0628 | 0,065 | 0,0632 | 0,0681 | 0,0771 |
| 0,0634 | 0,0636 | 0,0753 | 0,0712 | 0,0718 | 0,0687 | 0,0716 | 0,0695 | 0,0685 | 0,0729 | 0,0723 | 0,0677 | 0,0696 | 0,0684 | 0,077 | 0,0835 |
| 0,0704 | 0,0683 | 0,0781 | 0,0801 | 0,0811 | 0,0738 | 0,0812 | 0,0782 | 0,0741 | 0,0776 | 0,0862 | 0,0776 | 0,0786 | 0,0781 | 0,0912 | 0,0952 |
| 0,0788 | 0,0755 | 0,0904 | 0,0916 | 0,089 | 0,0832 | 0,092 | 0,0891 | 0,0864 | 0,0911 | 0,0981 | 0,0876 | 0,0894 | 0,0897 | 0,1067 | 0,1087 |
| 0,0884 | 0,083 | 0,1088 | 0,0999 | 0,0987 | 0,0896 | 0,1003 | 0,0994 | 0,0941 | 0,1023 | 0,1105 | 0,0967 | 0,0993 | 0,0967 | 0,1202 | 0,1206 |
| 0,0956 | 0,0907 | 0,1171 | 0,1076 | 0,1082 | 0,099 | 0,1099 | 0,1101 | 0,1023 | 0,1173 | 0,1329 | 0,0999 | 0,1098 | 0,109 | 0,142 | 0,1325 |
| 0,1046 | 0,0969 | 0,1326 | 0,1194 | 0,1272 | 0,1078 | 0,1287 | 0,1172 | 0,115 | 0,1338 | 0,1464 | 0,1158 | 0,1185 | 0,1193 | 0,1598 | 0,149 |
| 0,1221 | 0,11 | 0,1435 | 0,1329 | 0,1395 | 0,1246 | 0,1432 | 0,1335 | 0,1295 | 0,1542 | 0,1624 | 0,1284 | 0,1344 | 0,1332 | 0,175 | 0,1622 |
| 0,126 | 0,1198 | 0,1555 | 0,1401 | 0,1458 | 0,1358 | 0,1522 | 0,1556 | 0,1368 | 0,1609 | 0,1731 | 0,143 | 0,1531 | 0,1406 | 0,1869 | 0,19 |
| 0,1299 | 0,1265 | 0,1539 | 0,1403 | 0,1496 | 0,1436 | 0,1656 | 0,1644 | 0,1437 | 0,1677 | 0,1794 | 0,1474 | 0,1644 | 0,134 | 0,2062 | 0,2064 |
| 0,1357 | 0,1299 | 0,1543 | 0,1444 | 0,1504 | 0,1491 | 0,1749 | 0,168 | 0,1384 | 0,181 | 0,1909 | 0,1353 | 0,167 | 0,1332 | 0,2175 | 0,2279 |
| 0,1435 | 0,136 | 0,1613 | 0,1501 | 0,1502 | 0,1581 | 0,1953 | 0,1755 | 0,1461 | 0,1963 | 0,204 | 0,1366 | 0,174 | 0,1323 | 0,23 | 0,2381 |
| 0,1485 | 0,1407 | 0,1705 | 0,1441 | 0,1493 | 0,1667 | 0,192 | 0,1808 | 0,1454 | 0,2019 | 0,2156 | 0,1399 | 0,1791 | 0,1358 | 0,2451 | 0,2469 |
| 0,1549 | 0,1448 | 0,1655 | 0,1526 | 0,1469 | 0,1702 | 0,2047 | 0,1827 | 0,1529 | 0,2166 | 0,2318 | 0,1363 | 0,179 | 0,1408 | 0,2636 | 0,2558 |
| 0,1561 | 0,146 | 0,1639 | 0,1481 | 0,1514 | 0,1736 | 0,2204 | 0,1836 | 0,1544 | 0,2358 | 0,2419 | 0,1374 | 0,1801 | 0,1443 | 0,281 | 0,2645 |
| 0,1574 | 0,1478 | 0,1733 | 0,1509 | 0,165 | 0,1768 | 0,2264 | 0,1837 | 0,1506 | 0,2348 | 0,2532 | 0,138 | 0,1794 | 0,1573 | 0,2806 | 0,2724 |
| 0,1584 | 0,1487 | 0,1732 | 0,1502 | 0,164 | 0,1794 | 0,2393 | 0,1828 | 0,1492 | 0,2474 | 0,2653 | 0,1392 | 0,1784 | 0,1522 | 0,2943 | 0,2646 |
| 0,1607 | 0,1483 | 0,1731 | 0,1531 | 0,167 | 0,1808 | 0,2427 | 0,1837 | 0,15 | 0,2515 | 0,2734 | 0,141 | 0,1791 | 0,1516 | 0,2913 | 0,266 |
| 0,1571 | 0,1496 | 0,169 | 0,159 | 0,1732 | 0,184 | 0,255 | 0,1845 | 0,1621 | 0,2613 | 0,2816 | 0,1421 | 0,1824 | 0,1547 | 0,3001 | 0,2658 |
| 0,1612 | 0,1489 | 0,1789 | 0,1607 | 0,171 | 0,1836 | 0,2608 | 0,1834 | 0,1689 | 0,2687 | 0,285 | 0,144 | 0,1818 | 0,1508 | 0,3029 | 0,2648 |
| 0,1626 | 0,1491 | 0,1768 | 0,1646 | 0,1781 | 0,1866 | 0,2665 | 0,1845 | 0,1612 | 0,2718 | 0,2921 | 0,1454 | 0,1835 | 0,1518 | 0,3084 | 0,2596 |
| 0,1631 | 0,1499 | 0,1757 | 0,1673 | 0,1766 | 0,1874 | 0,2746 | 0,1855 | 0,1774 | 0,2761 | 0,3033 | 0,1476 | 0,1848 | 0,1506 | 0,3092 | 0,2588 |
| 0,1659 | 0,1499 | 0,1856 | 0,1741 | 0,1806 | 0,1885 | 0,2805 | 0,1862 | 0,1626 | 0,2796 | 0,3064 | 0,1495 | 0,1858 | 0,1462 | 0,3081 | 0,2599 |
| 0,1667 | 0,1503 | 0,1824 | 0,1792 | 0,181 | 0,1909 | 0,272 | 0,1876 | 0,1804 | 0,2805 | 0,3108 | 0,1502 | 0,1861 | 0,1422 | 0,3066 | 0,2576 |
| 0,1642 | 0,1508 | 0,175 | 0,1838 | 0,1758 | 0,1927 | 0,2723 | 0,189 | 0,1707 | 0,2798 | 0,3148 | 0,1509 | 0,1879 | 0,1378 | 0,3057 | 0,2558 |
| 0,1612 | 0,1508 | 0,1773 | 0,1907 | 0,1762 | 0,1938 | 0,2711 | 0,1911 | 0,1772 | 0,2803 | 0,3135 | 0,1494 | 0,1894 | 0,1332 | 0,3029 | 0,2546 |
| 0,1595 | 0,151 | 0,1787 | 0,1956 | 0,1749 | 0,1978 | 0,2723 | 0,1904 | 0,1755 | 0,2794 | 0,3107 | 0,1467 | 0,1896 | 0,129 | 0,2997 | 0,2529 |
| 0,1557 | 0,1516 | 0,1673 | 0,1971 | 0,1641 | 0,2004 | 0,2737 | 0,1937 | 0,1712 | 0,2797 | 0,3104 | 0,1419 | 0,191 | 0,12 | 0,3001 | 0,2446 |
| 0,1537 | 0,1504 | 0,1633 | 0,1982 | 0,1617 | 0,203 | 0,2725 | 0,1932 | 0,1856 | 0,282 | 0,3075 | 0,136 | 0,1906 | 0,1118 | 0,2988 | 0,2395 |
| 0,1491 | 0,1491 | 0,1583 | 0,1986 | 0,1558 | 0,2043 | 0,2744 | 0,1922 | 0,1848 | 0,2828 | 0,3089 | 0,1292 | 0,1891 | 0,104 | 0,2957 | 0,235 |

| L-glutamine | m-Hydroxy Ph | 2-Aminoethar | D-saccharide a | L-glutamic aci | D-Psicose | L-fucose | Phenyl Acetic | Formic acid | D-melibiose | Lactulose | MOPS no sugar | p-Hydroxy Phenyl Acetic Acid | Dulcitol | D-mannitol | D-galactose |
|---|---|---|---|---|---|---|---|---|---|---|---|---|---|---|---|
| 134 | 144 | 146 | 180 | 166 | 634 | 148 | 155 | 172 | 148 | 148 | 176 | 156 | 153 | 154 | 156 |
| 135 | 145 | 142 | 175 | 162 | 580 | 142 | 149 | 167 | 146 | 144 | 171 | 157 | 144 | 148 | 151 |
| 129 | 142 | 144 | 173 | 160 | 559 | 142 | 151 | 165 | 147 | 144 | 166 | 154 | 142 | 150 | 148 |
| 133 | 136 | 144 | 169 | 162 | 562 | 134 | 155 | 162 | 142 | 146 | 171 | 155 | 147 | 144 | 149 |
| 133 | 135 | 148 | 176 | 164 | 565 | 137 | 149 | 165 | 143 | 145 | 169 | 153 | 144 | 141 | 149 |
| 134 | 142 | 146 | 164 | 156 | 569 | 137 | 149 | 164 | 145 | 143 | 169 | 154 | 144 | 149 | 149 |
| 127 | 137 | 148 | 169 | 158 | 572 | 137 | 155 | 169 | 146 | 147 | 170 | 151 | 143 | 144 | 149 |
| 132 | 139 | 149 | 167 | 162 | 542 | 140 | 156 | 161 | 145 | 149 | 169 | 153 | 147 | 145 | 149 |
| 136 | 147 | 147 | 169 | 166 | 551 | 145 | 163 | 170 | 152 | 147 | 174 | 164 | 158 | 145 | 151 |
| 130 | 150 | 155 | 178 | 164 | 558 | 147 | 166 | 168 | 147 | 144 | 169 | 163 | 151 | 144 | 159 |
| 139 | 152 | 160 | 176 | 168 | 557 | 149 | 165 | 166 | 146 | 142 | 171 | 161 | 149 | 135 | 153 |
| 137 | 157 | 165 | 191 | 177 | 545 | 150 | 165 | 169 | 150 | 142 | 182 | 163 | 163 | 145 | 162 |
| 150 | 164 | 174 | 190 | 183 | 514 | 160 | 170 | 180 | 157 | 146 | 184 | 169 | 161 | 152 | 170 |
| 158 | 167 | 186 | 200 | 194 | 545 | 181 | 185 | 190 | 171 | 159 | 196 | 175 | 177 | 154 | 174 |
| 166 | 179 | 200 | 220 | 203 | 541 | 193 | 196 | 203 | 187 | 168 | 212 | 186 | 192 | 168 | 189 |
| 181 | 200 | 228 | 229 | 215 | 568 | 202 | 211 | 221 | 206 | 170 | 223 | 196 | 205 | 183 | 194 |
| 191 | 205 | 231 | 259 | 228 | 578 | 228 | 222 | 237 | 240 | 223 | 256 | 211 | 230 | 217 | 240 |
| 213 | 217 | 273 | 279 | 287 | 589 | 262 | 232 | 252 | 278 | 267 | 277 | 209 | 276 | 252 | 270 |
| 273 | 254 | 353 | 343 | 345 | 610 | 298 | 277 | 317 | 344 | 320 | 333 | 237 | 330 | 291 | 315 |
| 295 | 294 | 384 | 386 | 381 | 650 | 332 | 327 | 345 | 382 | 361 | 390 | 284 | 366 | 325 | 379 |
| 350 | 319 | 433 | 440 | 426 | 691 | 362 | 359 | 388 | 438 | 409 | 411 | 311 | 412 | 359 | 431 |
| 385 | 363 | 477 | 473 | 458 | 735 | 394 | 396 | 414 | 474 | 455 | 445 | 333 | 440 | 401 | 484 |
| 402 | 389 | 502 | 496 | 495 | 765 | 428 | 419 | 449 | 510 | 493 | 466 | 345 | 451 | 430 | 538 |
| 416 | 403 | 526 | 513 | 521 | 788 | 440 | 463 | 454 | 543 | 544 | 475 | 371 | 459 | 476 | 585 |
| 440 | 425 | 537 | 530 | 530 | 817 | 495 | 482 | 487 | 578 | 604 | 484 | 386 | 463 | 534 | 612 |
| 443 | 442 | 558 | 531 | 526 | 821 | 509 | 496 | 502 | 635 | 671 | 493 | 400 | 463 | 580 | 624 |
| 454 | 449 | 557 | 553 | 545 | 839 | 561 | 506 | 493 | 676 | 701 | 507 | 399 | 480 | 616 | 636 |
| 461 | 461 | 585 | 557 | 548 | 856 | 579 | 516 | 499 | 727 | 771 | 513 | 408 | 480 | 678 | 625 |
| 474 | 478 | 562 | 568 | 560 | 869 | 617 | 519 | 501 | 779 | 811 | 519 | 411 | 485 | 691 | 630 |
| 489 | 474 | 575 | 576 | 569 | 880 | 647 | 540 | 520 | 821 | 880 | 518 | 419 | 487 | 696 | 629 |
| 500 | 477 | 583 | 582 | 575 | 907 | 649 | 547 | 514 | 848 | 898 | 532 | 421 | 492 | 714 | 612 |
| 522 | 478 | 588 | 586 | 582 | 912 | 676 | 550 | 516 | 874 | 939 | 529 | 422 | 492 | 701 | 604 |
| 527 | 481 | 591 | 592 | 592 | 942 | 686 | 557 | 529 | 881 | 974 | 530 | 427 | 483 | 688 | 585 |
| 530 | 485 | 590 | 613 | 601 | 965 | 688 | 582 | 526 | 885 | 977 | 518 | 435 | 465 | 679 | 555 |
| 553 | 484 | 607 | 632 | 603 | 983 | 699 | 576 | 539 | 871 | 966 | 511 | 427 | 456 | 652 | 543 |
| 554 | 479 | 583 | 643 | 594 | 1005 | 732 | 585 | 547 | 854 | 954 | 497 | 430 | 419 | 640 | 523 |
| 542 | 488 | 574 | 656 | 583 | 1021 | 750 | 585 | 546 | 854 | 925 | 477 | 425 | 398 | 615 | 508 |
| 531 | 483 | 572 | 670 | 576 | 1050 | 782 | 589 | 558 | 825 | 908 | 448 | 424 | 357 | 592 | 482 |
| 522 | 477 | 546 | 670 | 553 | 1048 | 797 | 586 | 560 | 805 | 879 | 417 | 409 | 328 | 582 | 475 |
| 506 | 469 | 513 | 669 | 520 | 1076 | 806 | 602 | 570 | 792 | 844 | 387 | 400 | 281 | 559 | 463 |
| 480 | 468 | 471 | 658 | 495 | 1098 | 841 | 581 | 556 | 760 | 821 | 344 | 400 | 243 | 550 | 446 |
| 3442,78607 | 3378,51929 | 3289,47368 | 3225,37112 | 3206,62768 | 3135,13514 | 3092,36948 | 3082,48915 | 2913,50531 | 2635,65891 | 2620,71651 | 2089,552239 | 1808,74722 | 1707,77989 | 1654,13534 | 1598,67696 |

171

| D-lactose | D-xylose | Maltotriose | Maltose | D-fructose | D-glucose | N-acetyl-D-glu | Arabinose | D-Cellobiose |
|---|---|---|---|---|---|---|---|---|
| 0,0522 | 0,0523 | 0,0564 | 0,05 | 0,0529 | 0,0499 | 0,0468 | 0,0523 | 0,1938 |
| 0,0601 | 0,055 | 0,0577 | 0,0535 | 0,0534 | 0,0538 | 0,0477 | 0,0534 | 0,1333 |
| 0,0569 | 0,0563 | 0,0583 | 0,0519 | 0,0527 | 0,053 | 0,0484 | 0,0539 | 0,1147 |
| 0,0572 | 0,0592 | 0,0582 | 0,0529 | 0,0537 | 0,0521 | 0,0496 | 0,0549 | 0,0885 |
| 0,0606 | 0,0583 | 0,0567 | 0,0539 | 0,0545 | 0,0562 | 0,0494 | 0,0554 | 0,0856 |
| 0,0582 | 0,0629 | 0,0567 | 0,0545 | 0,054 | 0,0575 | 0,0507 | 0,056 | 0,0904 |
| 0,06 | 0,0617 | 0,0582 | 0,0568 | 0,0542 | 0,0583 | 0,0511 | 0,0564 | 0,1001 |
| 0,0589 | 0,0637 | 0,0631 | 0,0561 | 0,0563 | 0,0606 | 0,0529 | 0,0581 | 0,1206 |
| 0,0574 | 0,063 | 0,0664 | 0,0617 | 0,0594 | 0,0649 | 0,0569 | 0,0621 | 0,1218 |
| 0,0689 | 0,0669 | 0,068 | 0,0607 | 0,0651 | 0,0667 | 0,0608 | 0,065 | 0,1075 |
| 0,0631 | 0,0653 | 0,0665 | 0,0629 | 0,0627 | 0,0752 | 0,0631 | 0,0679 | 0,0945 |
| 0,065 | 0,0655 | 0,0742 | 0,0692 | 0,0644 | 0,0814 | 0,0673 | 0,0694 | 0,0861 |
| 0,0714 | 0,0694 | 0,0783 | 0,0675 | 0,0705 | 0,0923 | 0,0764 | 0,0763 | 0,0805 |
| 0,0934 | 0,082 | 0,0907 | 0,0851 | 0,0814 | 0,1076 | 0,089 | 0,0832 | 0,08 |
| 0,0971 | 0,0934 | 0,1051 | 0,1013 | 0,0982 | 0,1277 | 0,1018 | 0,096 | 0,0758 |
| 0,1123 | 0,098 | 0,1208 | 0,1027 | 0,1148 | 0,1454 | 0,115 | 0,1049 | 0,0798 |
| 0,1392 | 0,1149 | 0,1512 | 0,1189 | 0,1262 | 0,1671 | 0,1322 | 0,1139 | 0,0895 |
| 0,1709 | 0,1245 | 0,1627 | 0,1324 | 0,147 | 0,1954 | 0,1564 | 0,1295 | 0,0942 |
| 0,1961 | 0,1459 | 0,1847 | 0,1513 | 0,1734 | 0,241 | 0,1819 | 0,1514 | 0,1058 |
| 0,2098 | 0,1493 | 0,2009 | 0,1643 | 0,1943 | 0,2664 | 0,2024 | 0,1725 | 0,1076 |
| 0,2417 | 0,1714 | 0,2279 | 0,1721 | 0,2102 | 0,3048 | 0,2306 | 0,2008 | 0,1095 |
| 0,2666 | 0,1794 | 0,2511 | 0,1808 | 0,2208 | 0,3396 | 0,2462 | 0,2182 | 0,1139 |
| 0,2794 | 0,1865 | 0,2696 | 0,191 | 0,2486 | 0,3566 | 0,2698 | 0,2413 | 0,1154 |
| 0,3111 | 0,1984 | 0,2882 | 0,2075 | 0,2564 | 0,3562 | 0,2639 | 0,248 | 0,1199 |
| 0,3376 | 0,2159 | 0,3091 | 0,2284 | 0,2877 | 0,3503 | 0,2687 | 0,2518 | 0,1237 |
| 0,3194 | 0,2239 | 0,3306 | 0,2376 | 0,3042 | 0,3437 | 0,2562 | 0,2479 | 0,1203 |
| 0,3035 | 0,2322 | 0,3507 | 0,247 | 0,2992 | 0,3316 | 0,2572 | 0,2475 | 0,1178 |
| 0,3019 | 0,2454 | 0,3348 | 0,2583 | 0,3137 | 0,326 | 0,2537 | 0,2478 | 0,1139 |
| 0,3003 | 0,2534 | 0,3328 | 0,2672 | 0,3112 | 0,3211 | 0,2495 | 0,2434 | 0,112 |
| 0,3028 | 0,2578 | 0,332 | 0,2752 | 0,3076 | 0,3176 | 0,2495 | 0,244 | 0,1076 |
| 0,2953 | 0,2635 | 0,3223 | 0,2737 | 0,3072 | 0,3136 | 0,2495 | 0,2385 | 0,1024 |
| 0,293 | 0,2695 | 0,3122 | 0,2793 | 0,3052 | 0,3098 | 0,2413 | 0,2319 | 0,0956 |
| 0,296 | 0,2704 | 0,3183 | 0,2854 | 0,3034 | 0,3083 | 0,2416 | 0,2306 | 0,0919 |
| 0,2919 | 0,2741 | 0,3168 | 0,284 | 0,3021 | 0,3027 | 0,24 | 0,2288 | 0,0818 |
| 0,2889 | 0,2763 | 0,314 | 0,2856 | 0,301 | 0,3032 | 0,2363 | 0,2249 | 0,0764 |
| 0,2899 | 0,2783 | 0,3138 | 0,2808 | 0,299 | 0,3 | 0,2335 | 0,22 | 0,0697 |
| 0,2885 | 0,2763 | 0,3152 | 0,2783 | 0,2966 | 0,2975 | 0,2303 | 0,2176 | 0,0655 |
| 0,2843 | 0,276 | 0,3089 | 0,2769 | 0,2905 | 0,2942 | 0,2258 | 0,214 | 0,0594 |
| 0,2817 | 0,2758 | 0,3074 | 0,2738 | 0,2943 | 0,2914 | 0,2194 | 0,2081 | 0,0566 |
| 0,2789 | 0,2796 | 0,3036 | 0,2718 | 0,2923 | 0,2882 | 0,212 | 0,203 | 0,0533 |
| 0,2759 | 0,275 | 0,2999 | 0,2689 | 0,2883 | 0,2861 | 0,2066 | 0,1993 | 0,0516 |

| D-lactose | D-xylose | Maltotriose | Maltose | D-fructose | D-glucose | N-acetyl-D-glu | Arabinose | D-Cellobiose |
|---|---|---|---|---|---|---|---|---|
| 155 | 477 | 147 | 150 | 149 | 153 | 147 | 390 | 96 |
| 148 | 439 | 143 | 144 | 145 | 143 | 135 | 380 | 90 |
| 147 | 439 | 147 | 146 | 147 | 143 | 137 | 375 | 87 |
| 143 | 433 | 141 | 142 | 143 | 144 | 132 | 368 | 87 |
| 141 | 427 | 146 | 143 | 140 | 142 | 135 | 364 | 93 |
| 147 | 424 | 143 | 145 | 149 | 139 | 139 | 369 | 92 |
| 149 | 425 | 145 | 140 | 142 | 143 | 134 | 361 | 91 |
| 147 | 426 | 145 | 148 | 144 | 139 | 133 | 376 | 95 |
| 150 | 428 | 147 | 148 | 149 | 142 | 136 | 383 | 101 |
| 141 | 429 | 149 | 149 | 151 | 143 | 131 | 397 | 97 |
| 137 | 419 | 152 | 150 | 144 | 140 | 127 | 380 | 97 |
| 141 | 414 | 149 | 149 | 142 | 140 | 132 | 380 | 104 |
| 147 | 433 | 156 | 159 | 145 | 145 | 135 | 390 | 106 |
| 165 | 438 | 167 | 170 | 156 | 146 | 142 | 379 | 112 |
| 178 | 443 | 177 | 175 | 163 | 161 | 150 | 391 | 119 |
| 182 | 458 | 182 | 188 | 174 | 171 | 156 | 393 | 118 |
| 221 | 468 | 214 | 230 | 191 | 179 | 177 | 404 | 128 |
| 264 | 493 | 241 | 257 | 222 | 208 | 206 | 407 | 143 |
| 326 | 514 | 271 | 297 | 273 | 250 | 248 | 410 | 167 |
| 384 | 559 | 313 | 328 | 311 | 302 | 295 | 457 | 186 |
| 460 | 598 | 362 | 362 | 380 | 341 | 390 | 493 | 202 |
| 535 | 622 | 424 | 396 | 432 | 391 | 419 | 540 | 218 |
| 584 | 649 | 497 | 419 | 487 | 405 | 420 | 561 | 219 |
| 614 | 674 | 527 | 449 | 515 | 394 | 413 | 585 | 223 |
| 647 | 705 | 565 | 481 | 590 | 384 | 374 | 564 | 227 |
| 607 | 725 | 576 | 515 | 571 | 372 | 376 | 562 | 222 |
| 594 | 768 | 598 | 539 | 639 | 352 | 368 | 562 | 222 |
| 583 | 791 | 585 | 579 | 636 | 346 | 369 | 562 | 216 |
| 561 | 826 | 562 | 608 | 604 | 343 | 352 | 568 | 205 |
| 543 | 852 | 553 | 638 | 550 | 331 | 345 | 553 | 196 |
| 512 | 850 | 538 | 627 | 529 | 329 | 322 | 543 | 177 |
| 515 | 886 | 538 | 637 | 518 | 336 | 314 | 532 | 156 |
| 521 | 903 | 535 | 633 | 493 | 323 | 308 | 523 | 142 |
| 515 | 895 | 537 | 621 | 474 | 331 | 304 | 517 | 126 |
| 523 | 888 | 538 | 589 | 448 | 332 | 294 | 503 | 114 |
| 524 | 876 | 542 | 565 | 432 | 330 | 289 | 501 | 107 |
| 528 | 872 | 537 | 546 | 420 | 336 | 281 | 486 | 89 |
| 526 | 851 | 531 | 518 | 401 | 325 | 275 | 469 | 85 |
| 509 | 846 | 526 | 502 | 407 | 327 | 268 | 454 | 80 |
| 504 | 829 | 530 | 493 | 403 | 321 | 251 | 439 | 73 |
| 507 | 824 | 522 | 484 | 407 | 318 | 238 | 420 | 73 |
| 1573,53599 | 1558,14998 | 1540,04107 | 1525,81087 | 1096,0068 | 698,560542 | 569,461827 | 204,081633 | 161,744023 |

**Plate PM2A**:

| | | | | | | | | | | | | |
|---|---|---|---|---|---|---|---|---|---|---|---|---|
| OD | | | | | | | | | | | | |
| | N-Acetyl-DGlu | 3-0-ß-DGalact | Dextrin | L-Leucine | Quinic Acid | 3-Methyl Gluc | D-Glucosamin | Succinamic A | Turanose | L-Histidine | L-Isoleucine | D-Ribono-1,4- |
| 0s | 0,054 | 0,0537 | 0,0575 | 0,0557 | 0,0553 | 0,0549 | 0,0594 | 0,0553 | 0,0566 | 0,057 | 0,0545 | 0,0554 |
| 900s | 0,056 | 0,0537 | 0,0599 | 0,0564 | 0,057 | 0,0565 | 0,0604 | 0,0569 | 0,0584 | 0,0585 | 0,0548 | 0,0571 |
| 1800s | 0,058 | 0,0557 | 0,065 | 0,06 | 0,0605 | 0,0597 | 0,0641 | 0,0605 | 0,0628 | 0,0625 | 0,0586 | 0,0606 |
| 2700s | 0,0616 | 0,057 | 0,0722 | 0,0632 | 0,0654 | 0,0655 | 0,0693 | 0,0643 | 0,0684 | 0,0662 | 0,0626 | 0,066 |
| 3601s | 0,065 | 0,0597 | 0,0821 | 0,0664 | 0,0693 | 0,0737 | 0,0744 | 0,0688 | 0,0719 | 0,0705 | 0,0655 | 0,0691 |
| 4501s | 0,0683 | 0,0639 | 0,0919 | 0,0684 | 0,0737 | 0,0791 | 0,0803 | 0,0715 | 0,0759 | 0,0736 | 0,0671 | 0,0741 |
| 5401s | 0,0713 | 0,0669 | 0,1006 | 0,0702 | 0,0769 | 0,0839 | 0,0841 | 0,0757 | 0,0801 | 0,0771 | 0,0707 | 0,0778 |
| 6301s | 0,0747 | 0,0714 | 0,1073 | 0,0717 | 0,0833 | 0,0884 | 0,0873 | 0,0812 | 0,0842 | 0,0817 | 0,0748 | 0,0818 |
| 7201s | 0,0796 | 0,0759 | 0,116 | 0,0736 | 0,0942 | 0,0974 | 0,0962 | 0,0912 | 0,0964 | 0,0911 | 0,081 | 0,0933 |
| 8101s | 0,09 | 0,0805 | 0,1293 | 0,0778 | 0,1042 | 0,1042 | 0,1052 | 0,0991 | 0,1048 | 0,0997 | 0,0869 | 0,1028 |
| 9001s | 0,0966 | 0,0878 | 0,1398 | 0,0811 | 0,1142 | 0,1152 | 0,1172 | 0,1109 | 0,1153 | 0,1098 | 0,0948 | 0,1157 |
| 9901s | 0,1069 | 0,0929 | 0,1554 | 0,0832 | 0,1261 | 0,1299 | 0,1268 | 0,1204 | 0,1297 | 0,1148 | 0,1008 | 0,1251 |
| 10801s | 0,1176 | 0,1015 | 0,1697 | 0,0878 | 0,1349 | 0,14 | 0,14 | 0,1303 | 0,1376 | 0,1255 | 0,1087 | 0,138 |
| 11701s | 0,1255 | 0,1108 | 0,1853 | 0,0899 | 0,1448 | 0,1461 | 0,1507 | 0,1389 | 0,1441 | 0,1391 | 0,1141 | 0,1466 |
| 12601s | 0,1313 | 0,1197 | 0,1747 | 0,0928 | 0,1488 | 0,1495 | 0,1558 | 0,1401 | 0,1479 | 0,1431 | 0,1244 | 0,1483 |
| 13502s | 0,1362 | 0,1312 | 0,1944 | 0,1013 | 0,1526 | 0,1588 | 0,1596 | 0,148 | 0,1511 | 0,1474 | 0,1296 | 0,1516 |
| 14402s | 0,1422 | 0,1331 | 0,1837 | 0,1093 | 0,1577 | 0,1582 | 0,1694 | 0,1515 | 0,1564 | 0,1497 | 0,1327 | 0,1576 |
| 15302s | 0,1447 | 0,1392 | 0,1877 | 0,1175 | 0,1618 | 0,1646 | 0,1758 | 0,1569 | 0,1598 | 0,1557 | 0,1372 | 0,1601 |
| 16202s | 0,1458 | 0,1456 | 0,1866 | 0,1222 | 0,1622 | 0,1694 | 0,1806 | 0,1635 | 0,1662 | 0,156 | 0,1408 | 0,1637 |
| 17102s | 0,1479 | 0,1473 | 0,1876 | 0,1255 | 0,1701 | 0,1759 | 0,1899 | 0,1707 | 0,1703 | 0,1637 | 0,1489 | 0,1707 |
| 18002s | 0,1518 | 0,1424 | 0,1853 | 0,1275 | 0,1757 | 0,179 | 0,1936 | 0,1695 | 0,1746 | 0,1714 | 0,1526 | 0,1718 |
| 18902s | 0,1549 | 0,1416 | 0,1833 | 0,1332 | 0,1805 | 0,1786 | 0,2039 | 0,1754 | 0,1819 | 0,1775 | 0,1549 | 0,1787 |
| 19802s | 0,1565 | 0,145 | 0,1853 | 0,137 | 0,1873 | 0,1927 | 0,2112 | 0,1834 | 0,1854 | 0,1812 | 0,1601 | 0,1863 |
| 20702s | 0,1765 | 0,1469 | 0,1834 | 0,1409 | 0,2121 | 0,199 | 0,2199 | 0,1961 | 0,1979 | 0,1871 | 0,1655 | 0,204 |
| 21602s | 0,166 | 0,1499 | 0,1808 | 0,1462 | 0,2212 | 0,205 | 0,2309 | 0,1962 | 0,2029 | 0,1908 | 0,1618 | 0,2057 |
| 22503s | 0,1767 | 0,1534 | 0,1882 | 0,1509 | 0,2097 | 0,2081 | 0,2361 | 0,2018 | 0,206 | 0,1899 | 0,1698 | 0,2102 |
| 23403s | 0,1694 | 0,1549 | 0,1869 | 0,1505 | 0,2071 | 0,2036 | 0,2444 | 0,2019 | 0,2008 | 0,1923 | 0,1698 | 0,2046 |
| 24303s | 0,1721 | 0,1613 | 0,1882 | 0,1551 | 0,2221 | 0,2098 | 0,257 | 0,2062 | 0,2112 | 0,1973 | 0,1707 | 0,2185 |
| 25203s | 0,178 | 0,1655 | 0,1903 | 0,1571 | 0,2132 | 0,2167 | 0,2557 | 0,2078 | 0,2134 | 0,2013 | 0,1735 | 0,2149 |
| 26103s | 0,1797 | 0,1661 | 0,1945 | 0,1583 | 0,2227 | 0,2169 | 0,2709 | 0,2085 | 0,2211 | 0,2049 | 0,1764 | 0,2232 |
| 27003s | 0,1808 | 0,1669 | 0,1995 | 0,1621 | 0,2218 | 0,2164 | 0,2715 | 0,2126 | 0,2219 | 0,2103 | 0,1778 | 0,2249 |
| 27903s | 0,1831 | 0,1666 | 0,2022 | 0,1646 | 0,2294 | 0,218 | 0,2763 | 0,215 | 0,228 | 0,2122 | 0,1801 | 0,2277 |
| 28803s | 0,1814 | 0,167 | 0,2039 | 0,1651 | 0,2273 | 0,221 | 0,2788 | 0,2173 | 0,2295 | 0,2185 | 0,1813 | 0,2313 |
| 29703s | 0,1835 | 0,1682 | 0,2031 | 0,1667 | 0,2375 | 0,2217 | 0,2925 | 0,2198 | 0,2342 | 0,2223 | 0,187 | 0,2392 |
| 30603s | 0,1837 | 0,1687 | 0,2035 | 0,1702 | 0,2397 | 0,2243 | 0,2957 | 0,2217 | 0,2328 | 0,2287 | 0,1894 | 0,2405 |
| 31503s | 0,1837 | 0,1681 | 0,2003 | 0,1706 | 0,2385 | 0,228 | 0,3002 | 0,2247 | 0,2362 | 0,2314 | 0,1895 | 0,242 |
| gfp | N-Acetyl-DGlu | 3-0-ß-DGalact | Dextrin | L-Leucine | Quinic Acid | 3-Methyl Gluc | D-Glucosamin | Succinamic A | Turanose | L-Histidine | L-Isoleucine | D-Ribono-1,4- |
| 0s | 151 | 202 | 187 | 183 | 184 | 158 | 707 | 137 | 159 | 175 | 175 | 180 |
| 900s | 152 | 189 | 177 | 179 | 181 | 163 | 668 | 138 | 159 | 179 | 172 | 176 |
| 1800s | 154 | 188 | 178 | 182 | 187 | 163 | 664 | 149 | 170 | 180 | 179 | 188 |
| 2700s | 161 | 188 | 183 | 200 | 201 | 161 | 665 | 161 | 184 | 198 | 194 | 198 |
| 3600s | 167 | 183 | 186 | 209 | 219 | 166 | 673 | 176 | 204 | 218 | 210 | 220 |
| 4500s | 188 | 188 | 195 | 228 | 231 | 194 | 677 | 191 | 224 | 242 | 220 | 238 |
| 5401s | 205 | 185 | 206 | 230 | 257 | 222 | 684 | 211 | 239 | 255 | 241 | 244 |
| 6301s | 210 | 195 | 230 | 238 | 281 | 252 | 678 | 231 | 257 | 281 | 258 | 279 |
| 7201s | 238 | 196 | 271 | 255 | 322 | 297 | 720 | 271 | 298 | 314 | 294 | 323 |
| 8101s | 281 | 203 | 325 | 268 | 350 | 339 | 738 | 298 | 335 | 352 | 317 | 357 |
| 9001s | 315 | 210 | 380 | 293 | 386 | 372 | 759 | 337 | 370 | 381 | 341 | 395 |
| 9901s | 366 | 226 | 476 | 307 | 442 | 460 | 822 | 383 | 425 | 419 | 376 | 457 |
| 10801s | 443 | 260 | 603 | 336 | 504 | 537 | 894 | 455 | 498 | 469 | 410 | 509 |
| 11701s | 562 | 310 | 760 | 351 | 587 | 622 | 958 | 545 | 592 | 541 | 448 | 592 |
| 12601s | 682 | 379 | 830 | 368 | 659 | 707 | 1058 | 583 | 661 | 608 | 525 | 677 |
| 13501s | 823 | 477 | 954 | 420 | 733 | 782 | 1133 | 656 | 719 | 706 | 593 | 739 |
| 14402s | 985 | 579 | 998 | 476 | 807 | 853 | 1242 | 719 | 787 | 750 | 662 | 804 |
| 15302s | 1124 | 736 | 1103 | 544 | 874 | 927 | 1312 | 794 | 846 | 823 | 729 | 876 |
| 16202s | 1273 | 913 | 1136 | 632 | 946 | 974 | 1429 | 862 | 926 | 890 | 794 | 953 |
| 17102s | 1387 | 1071 | 1229 | 717 | 1029 | 1033 | 1506 | 949 | 1006 | 960 | 874 | 1028 |
| 18002s | 1547 | 1226 | 1282 | 808 | 1111 | 1130 | 1600 | 1013 | 1087 | 1042 | 927 | 1095 |
| 18902s | 1656 | 1342 | 1353 | 862 | 1185 | 1171 | 1695 | 1076 | 1164 | 1077 | 995 | 1152 |
| 19802s | 1751 | 1413 | 1412 | 925 | 1249 | 1256 | 1785 | 1169 | 1243 | 1146 | 1052 | 1243 |
| 20702s | 2058 | 1461 | 1463 | 1001 | 1459 | 1378 | 1891 | 1249 | 1379 | 1229 | 1117 | 1403 |
| 21602s | 2113 | 1564 | 1535 | 1098 | 1621 | 1445 | 2028 | 1330 | 1449 | 1379 | 1191 | 1503 |
| 22502s | 2226 | 1495 | 1592 | 1164 | 1548 | 1528 | 2102 | 1424 | 1480 | 1366 | 1254 | 1491 |
| 23402s | 2245 | 1538 | 1629 | 1204 | 1600 | 1540 | 2205 | 1460 | 1531 | 1412 | 1303 | 1539 |
| 24303s | 2402 | 1536 | 1698 | 1291 | 1784 | 1616 | 2340 | 1549 | 1661 | 1519 | 1339 | 1657 |
| 25203s | 2521 | 1547 | 1745 | 1326 | 1730 | 1696 | 2446 | 1636 | 1694 | 1605 | 1396 | 1674 |
| 26103s | 2597 | 1607 | 1795 | 1378 | 1893 | 1750 | 2648 | 1706 | 1825 | 1686 | 1447 | 1821 |
| 27003s | 2695 | 1614 | 1842 | 1425 | 1886 | 1847 | 2733 | 1771 | 1827 | 1735 | 1512 | 1835 |
| 27903s | 2791 | 1623 | 1873 | 1449 | 1983 | 1910 | 2881 | 1845 | 1929 | 1824 | 1556 | 1936 |
| 28803s | 2836 | 1626 | 1901 | 1486 | 2076 | 1999 | 3042 | 1935 | 1999 | 1907 | 1589 | 2006 |
| 29703s | 2901 | 1605 | 1916 | 1520 | 2278 | 2046 | 3196 | 1968 | 2085 | 1990 | 1655 | 2184 |
| 30603s | 2943 | 1584 | 1919 | 1518 | 2277 | 2103 | 3325 | 1970 | 2105 | 2066 | 1642 | 2168 |
| 31503s | 2943 | 1596 | 1915 | 1571 | 2334 | 2182 | 3498 | 2098 | 2215 | 2170 | 1711 | 2260 |
| ratio | 21526,5998 | 12185,3147 | 12100,8403 | 12080,0696 | 11735,8079 | 11692,6632 | 11590,5316 | 11576,1511 | 11447,6615 | 11439,2202 | 11377,7778 | 11146,8382 |

| Maltitol | Stachyose | Sedoheptulos | L-Alaninamide | D-Melezitose | Oxalic Acid | Hydroxy-LPro | D-Arabitol | Glycolic Acid | α-Methyl-DMa | L-Arabitol | Citraconic Aci | α-Keto-Valeric |
|---|---|---|---|---|---|---|---|---|---|---|---|---|
| 0,0516 | 0,0559 | 0,0561 | 0,0541 | 0,0548 | 0,0613 | 0,0551 | 0,0542 | 0,0533 | 0,0541 | 0,0543 | 0,0548 | 0,0536 |
| 0,0532 | 0,0577 | 0,0583 | 0,0557 | 0,0566 | 0,0633 | 0,0558 | 0,0557 | 0,0543 | 0,0552 | 0,0554 | 0,0564 | 0,055 |
| 0,0574 | 0,0614 | 0,0615 | 0,0588 | 0,0608 | 0,0662 | 0,0598 | 0,0603 | 0,0583 | 0,0584 | 0,0586 | 0,0596 | 0,0589 |
| 0,0615 | 0,0667 | 0,0658 | 0,0607 | 0,0643 | 0,0705 | 0,0645 | 0,0643 | 0,0616 | 0,0621 | 0,0623 | 0,0636 | 0,0622 |
| 0,0658 | 0,0703 | 0,0699 | 0,0651 | 0,0686 | 0,0747 | 0,068 | 0,0675 | 0,0645 | 0,0662 | 0,0674 | 0,0671 | 0,0657 |
| 0,0692 | 0,0739 | 0,0725 | 0,0666 | 0,0714 | 0,0782 | 0,0705 | 0,0737 | 0,0679 | 0,0693 | 0,0725 | 0,0706 | 0,0692 |
| 0,0728 | 0,0778 | 0,0755 | 0,0697 | 0,0747 | 0,0809 | 0,0755 | 0,0771 | 0,0718 | 0,0734 | 0,0753 | 0,0741 | 0,0746 |
| 0,0784 | 0,0819 | 0,0806 | 0,0737 | 0,081 | 0,086 | 0,0807 | 0,0819 | 0,0756 | 0,0787 | 0,0819 | 0,0797 | 0,0796 |
| 0,088 | 0,0934 | 0,0905 | 0,082 | 0,0904 | 0,0961 | 0,0911 | 0,0895 | 0,0851 | 0,0881 | 0,0907 | 0,089 | 0,0877 |
| 0,0962 | 0,1015 | 0,0985 | 0,0883 | 0,099 | 0,1037 | 0,1013 | 0,1001 | 0,0926 | 0,094 | 0,1001 | 0,0966 | 0,0978 |
| 0,1058 | 0,1107 | 0,1073 | 0,0966 | 0,1084 | 0,1121 | 0,1141 | 0,1157 | 0,1026 | 0,1057 | 0,1114 | 0,1044 | 0,1104 |
| 0,1172 | 0,1203 | 0,115 | 0,1025 | 0,1135 | 0,1131 | 0,1223 | 0,1223 | 0,1145 | 0,1165 | 0,1194 | 0,1116 | 0,1166 |
| 0,1259 | 0,1312 | 0,1272 | 0,1073 | 0,1275 | 0,1308 | 0,1349 | 0,1337 | 0,1234 | 0,1263 | 0,1351 | 0,1249 | 0,1309 |
| 0,136 | 0,1421 | 0,1398 | 0,1203 | 0,1368 | 0,1448 | 0,1438 | 0,1434 | 0,1323 | 0,1373 | 0,1431 | 0,1344 | 0,1431 |
| 0,1412 | 0,1444 | 0,1428 | 0,1277 | 0,1431 | 0,1478 | 0,1474 | 0,1504 | 0,1349 | 0,1374 | 0,1464 | 0,1375 | 0,1489 |
| 0,1448 | 0,1475 | 0,1485 | 0,1363 | 0,1446 | 0,1509 | 0,1527 | 0,1572 | 0,1422 | 0,1476 | 0,1526 | 0,1414 | 0,157 |
| 0,1509 | 0,1542 | 0,151 | 0,1368 | 0,1495 | 0,1575 | 0,1571 | 0,165 | 0,1454 | 0,1465 | 0,1536 | 0,1462 | 0,1532 |
| 0,1529 | 0,1568 | 0,1585 | 0,1432 | 0,1542 | 0,1604 | 0,159 | 0,1603 | 0,1492 | 0,1506 | 0,1582 | 0,1526 | 0,1591 |
| 0,1595 | 0,1577 | 0,157 | 0,1444 | 0,1584 | 0,1639 | 0,1608 | 0,1839 | 0,1511 | 0,1578 | 0,1602 | 0,1503 | 0,1608 |
| 0,1669 | 0,1656 | 0,1628 | 0,1557 | 0,17 | 0,1694 | 0,1708 | 0,1971 | 0,1591 | 0,166 | 0,1694 | 0,1609 | 0,1612 |
| 0,1701 | 0,171 | 0,1677 | 0,1531 | 0,1694 | 0,1721 | 0,1744 | 0,2049 | 0,16 | 0,166 | 0,1702 | 0,1629 | 0,1668 |
| 0,1719 | 0,1774 | 0,1707 | 0,1562 | 0,1725 | 0,1763 | 0,1772 | 0,2196 | 0,1662 | 0,1644 | 0,1748 | 0,1641 | 0,1686 |
| 0,1783 | 0,1873 | 0,1839 | 0,1593 | 0,1773 | 0,1843 | 0,1876 | 0,2053 | 0,1693 | 0,1744 | 0,1815 | 0,1721 | 0,1768 |
| 0,1942 | 0,1976 | 0,2063 | 0,1608 | 0,1926 | 0,1924 | 0,1964 | 0,2221 | 0,1814 | 0,1801 | 0,1781 | 0,1789 | 0,1912 |
| 0,1964 | 0,2029 | 0,2081 | 0,1675 | 0,1928 | 0,203 | 0,1939 | 0,2223 | 0,1804 | 0,1804 | 0,1823 | 0,192 | 0,1881 |
| 0,1908 | 0,201 | 0,194 | 0,1717 | 0,1921 | 0,2005 | 0,1994 | 0,216 | 0,1938 | 0,1807 | 0,1881 | 0,1883 | 0,191 |
| 0,1855 | 0,1987 | 0,197 | 0,1735 | 0,1872 | 0,2025 | 0,1988 | 0,2081 | 0,1879 | 0,183 | 0,1794 | 0,188 | 0,1894 |
| 0,1935 | 0,2081 | 0,2024 | 0,1742 | 0,1924 | 0,2064 | 0,2037 | 0,2054 | 0,1939 | 0,1822 | 0,1844 | 0,1902 | 0,1997 |
| 0,1953 | 0,2069 | 0,2037 | 0,1783 | 0,1955 | 0,2058 | 0,2096 | 0,1954 | 0,2074 | 0,1874 | 0,1884 | 0,194 | 0,2001 |
| 0,1948 | 0,2146 | 0,207 | 0,1788 | 0,1942 | 0,2097 | 0,2104 | 0,1965 | 0,2089 | 0,1883 | 0,1867 | 0,1997 | 0,2018 |
| 0,2021 | 0,2164 | 0,2099 | 0,1823 | 0,2018 | 0,2128 | 0,2142 | 0,199 | 0,21 | 0,1894 | 0,1909 | 0,2006 | 0,202 |
| 0,2083 | 0,2214 | 0,2125 | 0,1846 | 0,2065 | 0,218 | 0,2182 | 0,2014 | 0,2079 | 0,1905 | 0,1944 | 0,2042 | 0,2018 |
| 0,2115 | 0,2229 | 0,2156 | 0,1881 | 0,2109 | 0,2188 | 0,2209 | 0,203 | 0,2082 | 0,1927 | 0,1984 | 0,2083 | 0,2029 |
| 0,2104 | 0,2278 | 0,2198 | 0,1919 | 0,2084 | 0,2208 | 0,2274 | 0,2066 | 0,2131 | 0,1953 | 0,1989 | 0,2108 | 0,2075 |
| 0,2133 | 0,2278 | 0,2221 | 0,1944 | 0,2113 | 0,2287 | 0,2323 | 0,205 | 0,2161 | 0,1981 | 0,2007 | 0,2139 | 0,2115 |
| 0,2161 | 0,2286 | 0,2259 | 0,1971 | 0,2131 | 0,2275 | 0,2344 | 0,2065 | 0,2141 | 0,2052 | 0,2017 | 0,2183 | 0,2099 |

| Maltitol | Stachyose | Sedoheptulos | L-Alaninamide | D-Melezitose | Oxalic Acid | Hydroxy-LPro | D-Arabitol | Glycolic Acid | α-Methyl-DMa | L-Arabitol | Citraconic Aci | α-Keto-Valeric |
|---|---|---|---|---|---|---|---|---|---|---|---|---|
| 162 | 160 | 152 | 146 | 157 | 169 | 164 | 158 | 166 | 137 | 159 | 169 | 166 |
| 158 | 159 | 149 | 146 | 152 | 167 | 161 | 154 | 161 | 143 | 152 | 169 | 163 |
| 167 | 164 | 154 | 150 | 158 | 169 | 168 | 163 | 168 | 141 | 158 | 173 | 172 |
| 178 | 182 | 162 | 160 | 171 | 178 | 177 | 169 | 183 | 159 | 172 | 192 | 181 |
| 201 | 200 | 181 | 185 | 187 | 203 | 198 | 170 | 196 | 177 | 183 | 212 | 200 |
| 219 | 206 | 193 | 190 | 198 | 216 | 217 | 195 | 211 | 188 | 203 | 221 | 213 |
| 226 | 223 | 207 | 202 | 210 | 236 | 227 | 205 | 216 | 200 | 217 | 238 | 234 |
| 240 | 244 | 229 | 222 | 233 | 255 | 250 | 232 | 231 | 217 | 249 | 263 | 251 |
| 287 | 276 | 263 | 260 | 263 | 292 | 289 | 271 | 274 | 258 | 275 | 295 | 286 |
| 312 | 313 | 295 | 291 | 294 | 317 | 322 | 310 | 298 | 276 | 311 | 315 | 311 |
| 342 | 337 | 309 | 318 | 317 | 348 | 358 | 340 | 324 | 305 | 347 | 358 | 354 |
| 392 | 384 | 344 | 349 | 360 | 375 | 407 | 372 | 371 | 353 | 385 | 383 | 376 |
| 460 | 455 | 420 | 403 | 421 | 456 | 466 | 424 | 427 | 413 | 463 | 471 | 455 |
| 525 | 511 | 489 | 478 | 479 | 515 | 551 | 511 | 495 | 493 | 538 | 527 | 540 |
| 595 | 591 | 564 | 544 | 556 | 581 | 616 | 603 | 563 | 570 | 610 | 597 | 601 |
| 671 | 653 | 631 | 602 | 623 | 652 | 698 | 681 | 624 | 630 | 668 | 662 | 687 |
| 743 | 730 | 699 | 674 | 696 | 727 | 770 | 779 | 697 | 688 | 733 | 728 | 748 |
| 787 | 788 | 761 | 744 | 745 | 805 | 825 | 836 | 753 | 757 | 794 | 782 | 803 |
| 861 | 855 | 836 | 799 | 798 | 875 | 902 | 971 | 795 | 802 | 848 | 849 | 865 |
| 912 | 921 | 900 | 880 | 867 | 938 | 986 | 1070 | 867 | 895 | 927 | 903 | 906 |
| 998 | 999 | 973 | 943 | 908 | 1018 | 1034 | 1187 | 939 | 920 | 983 | 963 | 978 |
| 1071 | 1066 | 1017 | 978 | 979 | 1054 | 1088 | 1271 | 988 | 975 | 1035 | 1017 | 1014 |
| 1116 | 1131 | 1105 | 1048 | 1034 | 1116 | 1170 | 1316 | 1050 | 1036 | 1071 | 1075 | 1064 |
| 1258 | 1285 | 1247 | 1106 | 1173 | 1189 | 1244 | 1401 | 1145 | 1100 | 1139 | 1136 | 1164 |
| 1361 | 1368 | 1333 | 1189 | 1256 | 1324 | 1341 | 1427 | 1200 | 1174 | 1199 | 1237 | 1217 |
| 1322 | 1371 | 1323 | 1233 | 1242 | 1351 | 1387 | 1450 | 1312 | 1233 | 1235 | 1259 | 1278 |
| 1326 | 1406 | 1356 | 1274 | 1243 | 1406 | 1436 | 1436 | 1325 | 1247 | 1256 | 1314 | 1311 |
| 1406 | 1515 | 1391 | 1331 | 1315 | 1475 | 1520 | 1490 | 1402 | 1306 | 1300 | 1352 | 1388 |
| 1483 | 1544 | 1467 | 1345 | 1372 | 1551 | 1611 | 1434 | 1480 | 1401 | 1398 | 1393 | 1427 |
| 1522 | 1638 | 1523 | 1412 | 1425 | 1612 | 1687 | 1485 | 1547 | 1431 | 1440 | 1473 | 1482 |
| 1588 | 1666 | 1592 | 1455 | 1487 | 1636 | 1737 | 1512 | 1595 | 1486 | 1492 | 1522 | 1542 |
| 1674 | 1776 | 1676 | 1503 | 1571 | 1726 | 1834 | 1578 | 1650 | 1542 | 1562 | 1585 | 1602 |
| 1769 | 1876 | 1751 | 1544 | 1665 | 1784 | 1921 | 1616 | 1723 | 1578 | 1617 | 1680 | 1664 |
| 1815 | 1907 | 1838 | 1595 | 1710 | 1873 | 1990 | 1702 | 1838 | 1640 | 1653 | 1738 | 1713 |
| 1905 | 1982 | 1943 | 1626 | 1814 | 1933 | 2036 | 1738 | 1863 | 1737 | 1717 | 1780 | 1716 |
| 1990 | 2077 | 2022 | 1703 | 1876 | 1970 | 2105 | 1803 | 1902 | 1767 | 1744 | 1887 | 1805 |
| 11112,462 | 11100,1737 | 11012,9564 | 10888,1119 | 10859,1282 | 10836,3418 | 10825,4322 | 10801,0506 | 10796,0199 | 10787,5579 | 10753,0529 | 10507,6453 | 10486,2444 |

| ß-Hydroxy But | γ-Amino Buty | 5-Keto-DGluc | Xylitol | Caproic Acid | ß-Methyl-DGl | L-Arginine | D-Tagatose | Citramalic Ac | Sebacic Acid | L-Valine | N-Acetyl-LGlu | D,L-Carnitine |
|---|---|---|---|---|---|---|---|---|---|---|---|---|
| 0,0559 | 0,0539 | 0,0546 | 0,0553 | 0,0561 | 0,0534 | 0,0547 | 0,0562 | 0,0549 | 0,0553 | 0,0568 | 0,0555 | 0,0557 |
| 0,0569 | 0,0551 | 0,0556 | 0,0567 | 0,0568 | 0,0542 | 0,0559 | 0,058 | 0,0563 | 0,0564 | 0,0577 | 0,0568 | 0,0571 |
| 0,0608 | 0,059 | 0,0587 | 0,0597 | 0,0599 | 0,0598 | 0,0597 | 0,0603 | 0,0599 | 0,0608 | 0,0607 | 0,0605 | 0,0604 |
| 0,0649 | 0,0624 | 0,0616 | 0,0639 | 0,0636 | 0,0636 | 0,0629 | 0,064 | 0,064 | 0,0656 | 0,0646 | 0,0638 | 0,064 |
| 0,0682 | 0,0659 | 0,0654 | 0,0664 | 0,0673 | 0,0651 | 0,0668 | 0,0662 | 0,0678 | 0,0696 | 0,0688 | 0,0677 | 0,0677 |
| 0,0719 | 0,069 | 0,069 | 0,0705 | 0,0704 | 0,0696 | 0,07 | 0,0717 | 0,0711 | 0,0736 | 0,0721 | 0,0703 | 0,0702 |
| 0,0763 | 0,0738 | 0,0729 | 0,0748 | 0,0734 | 0,0743 | 0,0737 | 0,0776 | 0,0744 | 0,0795 | 0,076 | 0,0742 | 0,0749 |
| 0,0813 | 0,0789 | 0,0779 | 0,0796 | 0,0778 | 0,0792 | 0,0792 | 0,0806 | 0,0802 | 0,083 | 0,084 | 0,0785 | 0,0805 |
| 0,0912 | 0,0897 | 0,0866 | 0,0872 | 0,0827 | 0,0862 | 0,0891 | 0,0849 | 0,088 | 0,0942 | 0,0933 | 0,0891 | 0,0921 |
| 0,1012 | 0,097 | 0,0957 | 0,0993 | 0,0891 | 0,0918 | 0,0984 | 0,0956 | 0,0962 | 0,1036 | 0,1023 | 0,0975 | 0,101 |
| 0,1123 | 0,1079 | 0,1048 | 0,1093 | 0,096 | 0,1034 | 0,1104 | 0,1045 | 0,1063 | 0,1159 | 0,1117 | 0,1076 | 0,1103 |
| 0,1168 | 0,1185 | 0,111 | 0,1214 | 0,1095 | 0,1135 | 0,1191 | 0,1175 | 0,1133 | 0,1256 | 0,1178 | 0,1118 | 0,1175 |
| 0,1294 | 0,1285 | 0,1254 | 0,1304 | 0,1194 | 0,1234 | 0,1317 | 0,1252 | 0,1253 | 0,138 | 0,1301 | 0,1279 | 0,1313 |
| 0,1414 | 0,1385 | 0,1366 | 0,1423 | 0,13 | 0,1326 | 0,1435 | 0,1372 | 0,133 | 0,148 | 0,1407 | 0,1423 | 0,1421 |
| 0,1436 | 0,1387 | 0,1378 | 0,1459 | 0,147 | 0,1333 | 0,1453 | 0,144 | 0,1376 | 0,1494 | 0,1418 | 0,1412 | 0,1436 |
| 0,1477 | 0,1478 | 0,1432 | 0,1494 | 0,1502 | 0,1416 | 0,1482 | 0,1501 | 0,1394 | 0,1528 | 0,1491 | 0,1477 | 0,1504 |
| 0,1537 | 0,1492 | 0,1493 | 0,1536 | 0,1562 | 0,1419 | 0,1556 | 0,1559 | 0,1469 | 0,157 | 0,1494 | 0,1521 | 0,1528 |
| 0,1595 | 0,1518 | 0,1578 | 0,1581 | 0,1622 | 0,1463 | 0,1597 | 0,1572 | 0,151 | 0,1622 | 0,1596 | 0,1553 | 0,162 |
| 0,1617 | 0,1546 | 0,1573 | 0,1607 | 0,1638 | 0,1481 | 0,1597 | 0,1612 | 0,1541 | 0,1662 | 0,162 | 0,1563 | 0,1657 |
| 0,1685 | 0,1628 | 0,1599 | 0,1669 | 0,1676 | 0,1531 | 0,1686 | 0,1641 | 0,1599 | 0,1758 | 0,1667 | 0,168 | 0,1686 |
| 0,1725 | 0,1689 | 0,1645 | 0,1738 | 0,1733 | 0,1581 | 0,1744 | 0,1725 | 0,1659 | 0,1773 | 0,1631 | 0,1706 | 0,1651 |
| 0,1788 | 0,1662 | 0,1711 | 0,1762 | 0,1762 | 0,1609 | 0,1804 | 0,1782 | 0,1701 | 0,1813 | 0,1673 | 0,1743 | 0,1704 |
| 0,1838 | 0,179 | 0,1696 | 0,1816 | 0,1771 | 0,167 | 0,1828 | 0,1831 | 0,1772 | 0,1893 | 0,1751 | 0,1781 | 0,1772 |
| 0,2012 | 0,1886 | 0,1756 | 0,1935 | 0,1806 | 0,1717 | 0,1881 | 0,1994 | 0,1995 | 0,2083 | 0,1775 | 0,1803 | 0,1796 |
| 0,2 | 0,19 | 0,1805 | 0,2017 | 0,1885 | 0,1768 | 0,1875 | 0,2053 | 0,202 | 0,212 | 0,1843 | 0,1825 | 0,1864 |
| 0,2075 | 0,1933 | 0,1781 | 0,2045 | 0,1891 | 0,1858 | 0,1977 | 0,2009 | 0,1845 | 0,2086 | 0,1857 | 0,1955 | 0,1889 |
| 0,2029 | 0,1921 | 0,1821 | 0,2045 | 0,1933 | 0,1855 | 0,1966 | 0,1997 | 0,1894 | 0,206 | 0,1885 | 0,1916 | 0,1923 |
| 0,2124 | 0,1958 | 0,1813 | 0,2121 | 0,1915 | 0,1847 | 0,1992 | 0,2109 | 0,2006 | 0,2194 | 0,19 | 0,1937 | 0,1931 |
| 0,2194 | 0,2007 | 0,1848 | 0,2154 | 0,1912 | 0,1928 | 0,2057 | 0,2082 | 0,1928 | 0,2162 | 0,1912 | 0,2 | 0,1952 |
| 0,2323 | 0,2006 | 0,1833 | 0,2247 | 0,2022 | 0,1939 | 0,2062 | 0,2204 | 0,2063 | 0,2246 | 0,1921 | 0,2012 | 0,197 |
| 0,2245 | 0,2012 | 0,1847 | 0,2195 | 0,1961 | 0,1987 | 0,2124 | 0,2188 | 0,2018 | 0,223 | 0,1937 | 0,2061 | 0,1979 |
| 0,2255 | 0,201 | 0,1838 | 0,2287 | 0,2049 | 0,2052 | 0,2131 | 0,2281 | 0,208 | 0,2246 | 0,1956 | 0,2084 | 0,1997 |
| 0,2268 | 0,204 | 0,1836 | 0,2273 | 0,2006 | 0,2082 | 0,2222 | 0,226 | 0,2087 | 0,2279 | 0,1977 | 0,2145 | 0,2008 |
| 0,241 | 0,2068 | 0,1849 | 0,2349 | 0,2071 | 0,2059 | 0,2295 | 0,2303 | 0,2106 | 0,2374 | 0,1989 | 0,2212 | 0,2062 |
| 0,244 | 0,2092 | 0,1848 | 0,2334 | 0,2025 | 0,2109 | 0,2324 | 0,2302 | 0,2103 | 0,2423 | 0,199 | 0,2218 | 0,2051 |
| 0,2375 | 0,2139 | 0,1861 | 0,2353 | 0,2023 | 0,2169 | 0,2327 | 0,231 | 0,2135 | 0,2357 | 0,1998 | 0,2221 | 0,207 |

| ß-Hydroxy But | γ-Amino Buty | 5-Keto-DGluc | Xylitol | Caproic Acid | ß-Methyl-DGl | L-Arginine | D-Tagatose | Citramalic Ac | Sebacic Acid | L-Valine | N-Acetyl-LGlu | D,L-Carnitine |
|---|---|---|---|---|---|---|---|---|---|---|---|---|
| 144 | 144 | 251 | 158 | 177 | 147 | 152 | 170 | 162 | 182 | 175 | 148 | 156 |
| 149 | 148 | 255 | 158 | 172 | 145 | 153 | 169 | 160 | 176 | 172 | 148 | 153 |
| 156 | 157 | 264 | 166 | 172 | 152 | 159 | 176 | 166 | 182 | 177 | 150 | 163 |
| 166 | 165 | 279 | 179 | 183 | 164 | 169 | 182 | 185 | 203 | 190 | 161 | 174 |
| 187 | 178 | 298 | 193 | 195 | 181 | 186 | 196 | 203 | 215 | 200 | 184 | 186 |
| 196 | 193 | 323 | 210 | 221 | 197 | 203 | 210 | 217 | 243 | 215 | 192 | 205 |
| 215 | 213 | 339 | 220 | 228 | 210 | 210 | 231 | 236 | 260 | 233 | 212 | 224 |
| 235 | 237 | 347 | 234 | 252 | 226 | 233 | 234 | 254 | 278 | 269 | 233 | 246 |
| 268 | 262 | 382 | 263 | 273 | 271 | 278 | 249 | 292 | 323 | 293 | 275 | 280 |
| 291 | 302 | 407 | 304 | 302 | 291 | 299 | 291 | 314 | 343 | 326 | 297 | 311 |
| 329 | 335 | 439 | 336 | 327 | 323 | 335 | 316 | 340 | 385 | 357 | 334 | 343 |
| 369 | 375 | 461 | 359 | 367 | 365 | 381 | 338 | 367 | 418 | 406 | 348 | 379 |
| 424 | 455 | 529 | 426 | 424 | 430 | 452 | 390 | 436 | 480 | 464 | 439 | 458 |
| 504 | 530 | 608 | 508 | 471 | 474 | 507 | 447 | 482 | 550 | 537 | 500 | 529 |
| 566 | 604 | 660 | 568 | 560 | 540 | 584 | 516 | 552 | 621 | 594 | 563 | 594 |
| 631 | 654 | 727 | 635 | 620 | 585 | 657 | 577 | 611 | 692 | 662 | 633 | 657 |
| 709 | 721 | 806 | 703 | 720 | 642 | 737 | 661 | 688 | 748 | 734 | 702 | 726 |
| 765 | 781 | 875 | 766 | 807 | 683 | 782 | 731 | 754 | 800 | 808 | 760 | 805 |
| 807 | 850 | 933 | 837 | 909 | 722 | 846 | 797 | 818 | 871 | 863 | 823 | 861 |
| 897 | 916 | 979 | 901 | 960 | 787 | 925 | 845 | 866 | 947 | 930 | 903 | 917 |
| 965 | 962 | 1022 | 978 | 1038 | 832 | 963 | 935 | 933 | 1012 | 994 | 946 | 961 |
| 1025 | 999 | 1075 | 1044 | 1111 | 893 | 1039 | 985 | 974 | 1056 | 1054 | 1005 | 1024 |
| 1110 | 1080 | 1113 | 1106 | 1117 | 946 | 1098 | 1062 | 1041 | 1131 | 1104 | 1069 | 1090 |
| 1242 | 1211 | 1163 | 1238 | 1196 | 1004 | 1162 | 1236 | 1188 | 1271 | 1177 | 1139 | 1150 |
| 1339 | 1247 | 1221 | 1313 | 1277 | 1079 | 1271 | 1295 | 1302 | 1408 | 1227 | 1220 | 1213 |
| 1342 | 1318 | 1253 | 1357 | 1328 | 1154 | 1300 | 1288 | 1235 | 1380 | 1267 | 1249 | 1260 |
| 1360 | 1334 | 1306 | 1408 | 1396 | 1199 | 1373 | 1355 | 1254 | 1384 | 1300 | 1320 | 1291 |
| 1496 | 1399 | 1362 | 1508 | 1449 | 1256 | 1436 | 1465 | 1366 | 1481 | 1340 | 1371 | 1335 |
| 1524 | 1456 | 1417 | 1545 | 1496 | 1316 | 1503 | 1484 | 1346 | 1527 | 1371 | 1423 | 1345 |
| 1681 | 1508 | 1468 | 1660 | 1567 | 1368 | 1585 | 1600 | 1458 | 1621 | 1402 | 1493 | 1396 |
| 1664 | 1572 | 1512 | 1668 | 1552 | 1480 | 1628 | 1611 | 1444 | 1629 | 1462 | 1543 | 1446 |
| 1753 | 1653 | 1539 | 1767 | 1602 | 1532 | 1710 | 1695 | 1519 | 1705 | 1484 | 1623 | 1486 |
| 1808 | 1682 | 1582 | 1831 | 1619 | 1604 | 1791 | 1748 | 1578 | 1757 | 1525 | 1663 | 1526 |
| 2008 | 1733 | 1603 | 1930 | 1659 | 1680 | 1856 | 1792 | 1641 | 1897 | 1564 | 1734 | 1564 |
| 2010 | 1780 | 1624 | 1950 | 1647 | 1700 | 1881 | 1844 | 1653 | 1910 | 1578 | 1728 | 1601 |
| 2040 | 1814 | 1618 | 2022 | 1687 | 1793 | 1929 | 1911 | 1712 | 1941 | 1567 | 1766 | 1616 |
| 10440,5286 | 10437,5 | 10395,4373 | 10355,5556 | 10328,3174 | 10067,2783 | 9983,14607 | 9959,95423 | 9773,01387 | 9750,55432 | 9734,26573 | 9711,88475 | 9649,70258 |

| Sec-Butylamir | L-Lysine | i-Erythrito | D,L-Octopami | Oxalomalic A | Glycogen | Putrescine | γ-Cyclodextrir | Glycine | Amygdalin | L-Glucose | δ-Amino Valer |
|---|---|---|---|---|---|---|---|---|---|---|---|
| 0,0562 | 0,0539 | 0,053 | 0,0569 | 0,0562 | 0,0604 | 0,0545 | 0,0523 | 0,0569 | 0,0529 | 0,055 | 0,0543 |
| 0,0576 | 0,0538 | 0,0547 | 0,0558 | 0,058 | 0,0639 | 0,0555 | 0,0542 | 0,0585 | 0,0545 | 0,0572 | 0,0555 |
| 0,0608 | 0,0581 | 0,0576 | 0,0593 | 0,0613 | 0,068 | 0,0592 | 0,0592 | 0,0626 | 0,0591 | 0,0607 | 0,059 |
| 0,0642 | 0,0615 | 0,0608 | 0,0628 | 0,0662 | 0,072 | 0,0619 | 0,0646 | 0,0661 | 0,0624 | 0,0636 | 0,0633 |
| 0,0691 | 0,0656 | 0,0653 | 0,0675 | 0,0708 | 0,0776 | 0,0653 | 0,0683 | 0,0704 | 0,066 | 0,0679 | 0,067 |
| 0,0717 | 0,0687 | 0,0691 | 0,0702 | 0,0748 | 0,0843 | 0,068 | 0,0716 | 0,0738 | 0,0708 | 0,07 | 0,0694 |
| 0,0773 | 0,0727 | 0,0735 | 0,0745 | 0,0778 | 0,0916 | 0,0726 | 0,0762 | 0,0771 | 0,0742 | 0,0729 | 0,0724 |
| 0,0855 | 0,0785 | 0,0802 | 0,0814 | 0,0814 | 0,0994 | 0,0821 | 0,0811 | 0,0852 | 0,079 | 0,0776 | 0,0777 |
| 0,0952 | 0,0878 | 0,0889 | 0,0914 | 0,091 | 0,1124 | 0,0902 | 0,0873 | 0,0928 | 0,0864 | 0,0865 | 0,0875 |
| 0,1049 | 0,0976 | 0,0985 | 0,0991 | 0,0996 | 0,1204 | 0,0988 | 0,0988 | 0,1025 | 0,0964 | 0,0957 | 0,0941 |
| 0,1142 | 0,1111 | 0,1096 | 0,1084 | 0,1101 | 0,1284 | 0,1107 | 0,1055 | 0,1124 | 0,106 | 0,1015 | 0,1049 |
| 0,1235 | 0,1207 | 0,1189 | 0,1161 | 0,1234 | 0,1394 | 0,1182 | 0,1153 | 0,1242 | 0,108 | 0,1045 | 0,1109 |
| 0,1331 | 0,1316 | 0,1288 | 0,1296 | 0,1308 | 0,1438 | 0,1323 | 0,1255 | 0,134 | 0,1253 | 0,1185 | 0,1254 |
| 0,1424 | 0,1404 | 0,1417 | 0,14 | 0,1415 | 0,1595 | 0,1427 | 0,1357 | 0,1436 | 0,1361 | 0,1296 | 0,1381 |
| 0,1422 | 0,141 | 0,1428 | 0,1406 | 0,1476 | 0,1498 | 0,1429 | 0,1441 | 0,1449 | 0,1401 | 0,1334 | 0,1371 |
| 0,1498 | 0,1476 | 0,1546 | 0,1479 | 0,1491 | 0,16 | 0,1532 | 0,1442 | 0,1469 | 0,1442 | 0,1381 | 0,1465 |
| 0,1536 | 0,1542 | 0,1478 | 0,1517 | 0,1556 | 0,1605 | 0,1548 | 0,1503 | 0,1537 | 0,1472 | 0,1355 | 0,1498 |
| 0,161 | 0,1574 | 0,1549 | 0,1563 | 0,1615 | 0,1606 | 0,1625 | 0,1483 | 0,1598 | 0,1468 | 0,1408 | 0,1517 |
| 0,1663 | 0,1652 | 0,158 | 0,16 | 0,1616 | 0,1612 | 0,1661 | 0,1451 | 0,163 | 0,1471 | 0,1505 | 0,1596 |
| 0,1672 | 0,166 | 0,1701 | 0,1645 | 0,1684 | 0,1625 | 0,1665 | 0,1498 | 0,1739 | 0,1568 | 0,1577 | 0,1663 |
| 0,167 | 0,1657 | 0,1686 | 0,1625 | 0,1726 | 0,1621 | 0,1658 | 0,1511 | 0,177 | 0,1596 | 0,1564 | 0,1707 |
| 0,1705 | 0,1711 | 0,1645 | 0,1644 | 0,1806 | 0,1641 | 0,1698 | 0,1557 | 0,1764 | 0,1558 | 0,1616 | 0,1689 |
| 0,1786 | 0,1721 | 0,1703 | 0,1722 | 0,1843 | 0,1631 | 0,1759 | 0,153 | 0,1776 | 0,1597 | 0,1647 | 0,1789 |
| 0,1809 | 0,1779 | 0,1719 | 0,1727 | 0,2028 | 0,1621 | 0,1793 | 0,1546 | 0,1821 | 0,1609 | 0,1671 | 0,1755 |
| 0,1865 | 0,1793 | 0,1696 | 0,1785 | 0,2076 | 0,1652 | 0,1833 | 0,153 | 0,1922 | 0,1641 | 0,1681 | 0,1847 |
| 0,1897 | 0,1832 | 0,1756 | 0,1825 | 0,2006 | 0,1647 | 0,186 | 0,1556 | 0,1913 | 0,1703 | 0,1647 | 0,1793 |
| 0,1936 | 0,1866 | 0,1735 | 0,1844 | 0,2034 | 0,1672 | 0,1895 | 0,1563 | 0,1964 | 0,1706 | 0,1686 | 0,1789 |
| 0,1963 | 0,188 | 0,1747 | 0,1851 | 0,2156 | 0,1704 | 0,1881 | 0,1588 | 0,1996 | 0,1713 | 0,172 | 0,1862 |
| 0,1991 | 0,1927 | 0,1787 | 0,1883 | 0,2086 | 0,1736 | 0,1889 | 0,161 | 0,2047 | 0,177 | 0,1761 | 0,1857 |
| 0,2014 | 0,1936 | 0,1828 | 0,1895 | 0,2168 | 0,1778 | 0,1921 | 0,1632 | 0,2084 | 0,1798 | 0,181 | 0,1891 |
| 0,2026 | 0,1963 | 0,1848 | 0,1901 | 0,2155 | 0,1826 | 0,1923 | 0,1663 | 0,2138 | 0,1834 | 0,1862 | 0,1891 |
| 0,205 | 0,1977 | 0,1855 | 0,1942 | 0,2204 | 0,1837 | 0,1944 | 0,1672 | 0,2159 | 0,1855 | 0,1875 | 0,191 |
| 0,2063 | 0,2005 | 0,1889 | 0,1959 | 0,2208 | 0,1825 | 0,1971 | 0,1687 | 0,2222 | 0,1891 | 0,1912 | 0,196 |
| 0,2092 | 0,2 | 0,189 | 0,1992 | 0,2299 | 0,1829 | 0,1982 | 0,1716 | 0,2273 | 0,1895 | 0,1907 | 0,2009 |
| 0,2083 | 0,2009 | 0,1903 | 0,1999 | 0,2322 | 0,1849 | 0,1988 | 0,1764 | 0,2334 | 0,1894 | 0,1915 | 0,2073 |
| 0,2088 | 0,2022 | 0,194 | 0,1989 | 0,2325 | 0,1844 | 0,1986 | 0,1768 | 0,2393 | 0,1919 | 0,1941 | 0,2075 |

| Sec-Butylamir | L-Lysine | i-Erythrito | D,L-Octopami | Oxalomalic A | Glycogen | Putrescine | γ-Cyclodextrir | Glycine | Amygdalin | L-Glucose | δ-Amino Valer |
|---|---|---|---|---|---|---|---|---|---|---|---|
| 148 | 171 | 153 | 173 | 262 | 174 | 160 | 151 | 188 | 151 | 153 | 147 |
| 149 | 175 | 147 | 165 | 257 | 166 | 161 | 152 | 180 | 150 | 147 | 147 |
| 161 | 171 | 154 | 175 | 266 | 167 | 171 | 158 | 185 | 153 | 154 | 154 |
| 168 | 187 | 165 | 181 | 274 | 183 | 181 | 176 | 199 | 169 | 164 | 166 |
| 185 | 205 | 170 | 203 | 290 | 204 | 192 | 185 | 224 | 181 | 178 | 179 |
| 200 | 216 | 193 | 216 | 304 | 226 | 208 | 202 | 233 | 197 | 188 | 198 |
| 213 | 233 | 205 | 238 | 320 | 248 | 222 | 224 | 253 | 216 | 205 | 214 |
| 244 | 258 | 233 | 257 | 337 | 280 | 251 | 241 | 282 | 233 | 218 | 225 |
| 265 | 294 | 265 | 298 | 373 | 311 | 275 | 263 | 317 | 259 | 261 | 268 |
| 296 | 321 | 289 | 318 | 398 | 356 | 299 | 288 | 342 | 276 | 281 | 301 |
| 331 | 358 | 330 | 353 | 441 | 393 | 339 | 329 | 385 | 307 | 313 | 324 |
| 373 | 405 | 361 | 400 | 464 | 456 | 388 | 378 | 446 | 329 | 337 | 360 |
| 445 | 481 | 436 | 465 | 530 | 531 | 449 | 450 | 525 | 411 | 408 | 435 |
| 509 | 542 | 515 | 548 | 596 | 606 | 536 | 526 | 584 | 478 | 479 | 507 |
| 579 | 624 | 593 | 640 | 662 | 666 | 604 | 605 | 647 | 542 | 538 | 585 |
| 653 | 675 | 673 | 710 | 736 | 734 | 659 | 666 | 699 | 604 | 608 | 645 |
| 728 | 746 | 727 | 790 | 805 | 774 | 745 | 738 | 758 | 664 | 649 | 711 |
| 829 | 828 | 781 | 873 | 854 | 849 | 847 | 781 | 806 | 705 | 712 | 775 |
| 825 | 876 | 828 | 895 | 933 | 902 | 861 | 843 | 863 | 741 | 769 | 824 |
| 878 | 921 | 924 | 969 | 990 | 972 | 920 | 887 | 933 | 797 | 852 | 889 |
| 936 | 977 | 932 | 1009 | 1060 | 992 | 954 | 906 | 977 | 841 | 862 | 931 |
| 992 | 1035 | 964 | 1045 | 1116 | 1047 | 1019 | 948 | 1024 | 882 | 905 | 971 |
| 1067 | 1082 | 1030 | 1120 | 1169 | 1109 | 1080 | 1006 | 1076 | 932 | 960 | 1021 |
| 1143 | 1148 | 1063 | 1179 | 1316 | 1144 | 1147 | 1029 | 1133 | 951 | 1003 | 1093 |
| 1200 | 1216 | 1099 | 1227 | 1435 | 1189 | 1203 | 1068 | 1203 | 1024 | 1046 | 1112 |
| 1272 | 1285 | 1149 | 1270 | 1397 | 1217 | 1242 | 1125 | 1258 | 1037 | 1075 | 1178 |
| 1318 | 1302 | 1166 | 1301 | 1418 | 1255 | 1259 | 1137 | 1302 | 1098 | 1137 | 1214 |
| 1368 | 1354 | 1200 | 1337 | 1557 | 1266 | 1300 | 1179 | 1361 | 1109 | 1153 | 1251 |
| 1404 | 1394 | 1251 | 1367 | 1537 | 1283 | 1316 | 1202 | 1414 | 1164 | 1193 | 1317 |
| 1434 | 1438 | 1275 | 1403 | 1685 | 1329 | 1363 | 1240 | 1474 | 1208 | 1228 | 1339 |
| 1480 | 1487 | 1336 | 1437 | 1647 | 1356 | 1397 | 1268 | 1533 | 1240 | 1277 | 1361 |
| 1536 | 1506 | 1378 | 1459 | 1700 | 1385 | 1446 | 1292 | 1586 | 1311 | 1318 | 1406 |
| 1571 | 1557 | 1423 | 1502 | 1744 | 1391 | 1471 | 1323 | 1654 | 1339 | 1373 | 1441 |
| 1589 | 1565 | 1473 | 1527 | 1858 | 1377 | 1487 | 1321 | 1723 | 1374 | 1392 | 1487 |
| 1609 | 1577 | 1488 | 1526 | 1887 | 1347 | 1499 | 1313 | 1787 | 1390 | 1403 | 1489 |
| 1612 | 1592 | 1498 | 1526 | 1940 | 1327 | 1495 | 1298 | 1834 | 1403 | 1395 | 1513 |
| 9593,70904 | 9581,92852 | 9539,00709 | 9528,16901 | 9517,86727 | 9298,3871 | 9264,39972 | 9212,85141 | 9024,12281 | 9007,19424 | 8928,82818 | 8916,44909 |

| 2,3-Butanedic | ß-Methyl-DXy | Malonic Acid | L-Pyroglutami | D-Lactic Acid | D-Arabinos | L-Tartaric Aci | ß-Cyclodextri | L-Methionine | Palatinose | L-Homoserine | α-Methyl-DGl | Inulin |
|---|---|---|---|---|---|---|---|---|---|---|---|---|
| 0,0541 | 0,0556 | 0,0538 | 0,0555 | 0,0544 | 0,058 | 0,0532 | 0,0526 | 0,053 | 0,0559 | 0,0553 | 0,0484 | 0,059 |
| 0,0548 | 0,0563 | 0,0552 | 0,0562 | 0,0556 | 0,0601 | 0,0537 | 0,0545 | 0,0535 | 0,0562 | 0,0566 | 0,0492 | 0,0595 |
| 0,0583 | 0,059 | 0,0579 | 0,0589 | 0,058 | 0,0646 | 0,0571 | 0,0598 | 0,0578 | 0,0588 | 0,0607 | 0,0518 | 0,0625 |
| 0,0628 | 0,0615 | 0,0619 | 0,0626 | 0,0628 | 0,0684 | 0,0609 | 0,0648 | 0,0603 | 0,0613 | 0,0645 | 0,0548 | 0,0661 |
| 0,0658 | 0,0647 | 0,0655 | 0,0671 | 0,0657 | 0,0726 | 0,0653 | 0,0676 | 0,064 | 0,0643 | 0,0685 | 0,0583 | 0,0688 |
| 0,0684 | 0,0658 | 0,0687 | 0,0694 | 0,0684 | 0,078 | 0,0683 | 0,0722 | 0,068 | 0,0655 | 0,0722 | 0,0629 | 0,0717 |
| 0,0736 | 0,0679 | 0,0718 | 0,0748 | 0,0721 | 0,0806 | 0,0722 | 0,0767 | 0,0717 | 0,0678 | 0,0771 | 0,0687 | 0,0758 |
| 0,0783 | 0,0789 | 0,0766 | 0,0798 | 0,08 | 0,0849 | 0,0785 | 0,0821 | 0,0766 | 0,0725 | 0,083 | 0,0736 | 0,0842 |
| 0,0882 | 0,0801 | 0,0883 | 0,0911 | 0,0909 | 0,0949 | 0,0872 | 0,0892 | 0,0866 | 0,0776 | 0,0934 | 0,0795 | 0,0917 |
| 0,0973 | 0,0867 | 0,0947 | 0,1003 | 0,0995 | 0,1035 | 0,0965 | 0,1005 | 0,0967 | 0,0826 | 0,1039 | 0,0863 | 0,1017 |
| 0,1106 | 0,0974 | 0,1041 | 0,1107 | 0,1091 | 0,1118 | 0,1062 | 0,1064 | 0,1077 | 0,0911 | 0,1167 | 0,1018 | 0,1068 |
| 0,1208 | 0,1071 | 0,1076 | 0,1151 | 0,1169 | 0,1215 | 0,1157 | 0,1151 | 0,1184 | 0,1014 | 0,119 | 0,1122 | 0,1184 |
| 0,1313 | 0,1226 | 0,1235 | 0,1289 | 0,129 | 0,1306 | 0,1286 | 0,1232 | 0,1296 | 0,1091 | 0,1231 | 0,1202 | 0,1278 |
| 0,1371 | 0,1328 | 0,1355 | 0,1424 | 0,1367 | 0,1392 | 0,1349 | 0,1329 | 0,1387 | 0,1173 | 0,1266 | 0,1306 | 0,1359 |
| 0,1404 | 0,136 | 0,1362 | 0,1392 | 0,1348 | 0,1452 | 0,1395 | 0,1381 | 0,1427 | 0,1233 | 0,1281 | 0,1384 | 0,143 |
| 0,1485 | 0,1399 | 0,1404 | 0,1498 | 0,1409 | 0,1473 | 0,1438 | 0,1404 | 0,1449 | 0,1371 | 0,1321 | 0,1419 | 0,1457 |
| 0,1536 | 0,145 | 0,1409 | 0,1489 | 0,1462 | 0,1516 | 0,1512 | 0,144 | 0,1479 | 0,1391 | 0,1342 | 0,1449 | 0,1512 |
| 0,1589 | 0,1461 | 0,1453 | 0,1583 | 0,1471 | 0,1547 | 0,1582 | 0,1422 | 0,1537 | 0,1476 | 0,1393 | 0,1489 | 0,1531 |
| 0,161 | 0,1486 | 0,1509 | 0,1614 | 0,1531 | 0,1559 | 0,1586 | 0,1409 | 0,1552 | 0,1523 | 0,142 | 0,1511 | 0,1466 |
| 0,1616 | 0,149 | 0,1659 | 0,1637 | 0,1588 | 0,1629 | 0,1629 | 0,147 | 0,1554 | 0,1568 | 0,1463 | 0,1581 | 0,148 |
| 0,1649 | 0,156 | 0,1638 | 0,1621 | 0,1556 | 0,1635 | 0,164 | 0,1504 | 0,1584 | 0,1552 | 0,1509 | 0,1589 | 0,1466 |
| 0,1679 | 0,1568 | 0,1665 | 0,1661 | 0,1614 | 0,1648 | 0,1684 | 0,1533 | 0,1631 | 0,1565 | 0,1543 | 0,1631 | 0,1453 |
| 0,1713 | 0,1607 | 0,1692 | 0,173 | 0,1654 | 0,1701 | 0,1709 | 0,154 | 0,1655 | 0,1587 | 0,1569 | 0,168 | 0,1455 |
| 0,1721 | 0,1613 | 0,173 | 0,1757 | 0,1708 | 0,1712 | 0,1754 | 0,1507 | 0,1675 | 0,1641 | 0,1655 | 0,1765 | 0,1456 |
| 0,1769 | 0,1675 | 0,1769 | 0,1812 | 0,1761 | 0,1768 | 0,1796 | 0,1504 | 0,1698 | 0,1762 | 0,1655 | 0,1811 | 0,1475 |
| 0,1769 | 0,166 | 0,1844 | 0,1798 | 0,1769 | 0,1797 | 0,1772 | 0,1507 | 0,1685 | 0,1755 | 0,1656 | 0,1835 | 0,1524 |
| 0,1797 | 0,1706 | 0,1829 | 0,1832 | 0,1787 | 0,1762 | 0,1797 | 0,1531 | 0,1718 | 0,1827 | 0,1659 | 0,1814 | 0,1584 |
| 0,1801 | 0,1739 | 0,1853 | 0,1827 | 0,1788 | 0,1829 | 0,1803 | 0,1545 | 0,1719 | 0,1721 | 0,1678 | 0,1855 | 0,1627 |
| 0,1829 | 0,1785 | 0,1923 | 0,1838 | 0,182 | 0,1857 | 0,1841 | 0,1564 | 0,1742 | 0,1793 | 0,1692 | 0,1869 | 0,166 |
| 0,1838 | 0,182 | 0,1923 | 0,1852 | 0,1847 | 0,1858 | 0,1855 | 0,159 | 0,1735 | 0,1842 | 0,1721 | 0,1944 | 0,1681 |
| 0,1849 | 0,1884 | 0,1969 | 0,188 | 0,1863 | 0,191 | 0,1874 | 0,1619 | 0,1763 | 0,182 | 0,1727 | 0,1941 | 0,166 |
| 0,1871 | 0,1877 | 0,1975 | 0,1912 | 0,1864 | 0,1915 | 0,1874 | 0,1614 | 0,1778 | 0,194 | 0,1739 | 0,1975 | 0,1689 |
| 0,1884 | 0,1872 | 0,2021 | 0,1899 | 0,1873 | 0,1965 | 0,1896 | 0,1635 | 0,1788 | 0,1886 | 0,1766 | 0,2007 | 0,1712 |
| 0,1893 | 0,1888 | 0,2071 | 0,1924 | 0,1862 | 0,1977 | 0,1908 | 0,167 | 0,1789 | 0,1909 | 0,1769 | 0,2016 | 0,1704 |
| 0,1885 | 0,188 | 0,2072 | 0,1916 | 0,1865 | 0,2004 | 0,191 | 0,1697 | 0,1786 | 0,1788 | 0,1796 | 0,2058 | 0,1685 |
| 0,1907 | 0,1898 | 0,2094 | 0,1928 | 0,1869 | 0,2043 | 0,1919 | 0,1694 | 0,1777 | 0,1793 | 0,1798 | 0,2066 | 0,1662 |

| 2,3-Butanedic | ß-Methyl-DXy | Malonic Acid | L-Pyroglutami | D-Lactic Acid | D-Arabinos | L-Tartaric Aci | ß-Cyclodextri | L-Methionine | Palatinose | L-Homoserine | α-Methyl-DGl | Inulin |
|---|---|---|---|---|---|---|---|---|---|---|---|---|
| 177 | 151 | 172 | 157 | 177 | 519 | 181 | 167 | 163 | 156 | 166 | 151 | 155 |
| 174 | 108 | 173 | 156 | 172 | 514 | 177 | 169 | 164 | 148 | 168 | 152 | 146 |
| 186 | 82 | 171 | 165 | 175 | 527 | 177 | 168 | 163 | 151 | 176 | 154 | 151 |
| 191 | 90 | 184 | 178 | 190 | 536 | 190 | 183 | 179 | 148 | 182 | 161 | 164 |
| 205 | 100 | 206 | 190 | 201 | 552 | 199 | 194 | 185 | 151 | 200 | 170 | 173 |
| 211 | 90 | 220 | 209 | 209 | 555 | 225 | 215 | 209 | 161 | 219 | 189 | 182 |
| 232 | 98 | 229 | 225 | 226 | 565 | 237 | 234 | 221 | 187 | 237 | 199 | 202 |
| 255 | 103 | 246 | 253 | 253 | 585 | 259 | 257 | 237 | 209 | 259 | 218 | 230 |
| 278 | 119 | 289 | 282 | 273 | 614 | 296 | 276 | 277 | 220 | 297 | 239 | 255 |
| 311 | 149 | 301 | 312 | 302 | 636 | 319 | 301 | 301 | 241 | 325 | 262 | 282 |
| 340 | 309 | 339 | 345 | 331 | 652 | 352 | 339 | 328 | 265 | 365 | 308 | 298 |
| 386 | 333 | 355 | 379 | 383 | 704 | 392 | 382 | 385 | 278 | 409 | 342 | 341 |
| 451 | 424 | 439 | 451 | 442 | 774 | 474 | 462 | 451 | 309 | 444 | 366 | 394 |
| 540 | 476 | 494 | 522 | 492 | 840 | 539 | 537 | 529 | 335 | 486 | 426 | 466 |
| 603 | 559 | 549 | 575 | 562 | 904 | 619 | 608 | 598 | 405 | 516 | 491 | 539 |
| 671 | 612 | 618 | 639 | 638 | 977 | 701 | 662 | 654 | 479 | 570 | 558 | 603 |
| 744 | 681 | 673 | 703 | 702 | 1039 | 758 | 724 | 735 | 555 | 601 | 615 | 668 |
| 825 | 717 | 729 | 765 | 751 | 1078 | 810 | 781 | 801 | 639 | 630 | 667 | 704 |
| 855 | 782 | 788 | 822 | 815 | 1136 | 877 | 810 | 845 | 692 | 668 | 705 | 756 |
| 911 | 805 | 854 | 887 | 861 | 1190 | 918 | 850 | 900 | 763 | 707 | 773 | 813 |
| 958 | 849 | 899 | 925 | 898 | 1235 | 945 | 891 | 930 | 795 | 748 | 807 | 842 |
| 997 | 897 | 956 | 972 | 937 | 1267 | 1000 | 911 | 964 | 816 | 769 | 850 | 880 |
| 1052 | 949 | 1012 | 1031 | 975 | 1296 | 1025 | 960 | 987 | 853 | 802 | 887 | 918 |
| 1098 | 974 | 1044 | 1082 | 1028 | 1322 | 1066 | 985 | 1022 | 876 | 848 | 946 | 950 |
| 1122 | 1030 | 1141 | 1108 | 1083 | 1371 | 1107 | 1033 | 1063 | 912 | 906 | 997 | 984 |
| 1155 | 1056 | 1168 | 1147 | 1097 | 1417 | 1134 | 1053 | 1088 | 930 | 919 | 1058 | 1035 |
| 1173 | 1109 | 1197 | 1171 | 1126 | 1455 | 1161 | 1061 | 1124 | 961 | 936 | 1070 | 1040 |
| 1219 | 1155 | 1248 | 1180 | 1170 | 1474 | 1190 | 1108 | 1142 | 995 | 974 | 1121 | 1066 |
| 1260 | 1186 | 1282 | 1241 | 1215 | 1550 | 1226 | 1124 | 1174 | 1049 | 1001 | 1139 | 1091 |
| 1281 | 1206 | 1325 | 1260 | 1251 | 1577 | 1253 | 1147 | 1213 | 1062 | 1028 | 1182 | 1117 |
| 1317 | 1269 | 1364 | 1301 | 1292 | 1619 | 1296 | 1185 | 1232 | 1109 | 1068 | 1214 | 1118 |
| 1363 | 1284 | 1417 | 1330 | 1308 | 1655 | 1314 | 1199 | 1233 | 1128 | 1090 | 1277 | 1099 |
| 1390 | 1312 | 1445 | 1329 | 1333 | 1672 | 1332 | 1196 | 1231 | 1162 | 1127 | 1305 | 1099 |
| 1380 | 1294 | 1486 | 1319 | 1320 | 1687 | 1340 | 1182 | 1233 | 1171 | 1133 | 1333 | 1077 |
| 1377 | 1295 | 1494 | 1317 | 1322 | 1712 | 1346 | 1174 | 1202 | 1166 | 1155 | 1386 | 1042 |
| 1371 | 1300 | 1493 | 1315 | 1292 | 1749 | 1335 | 1130 | 1183 | 1156 | 1166 | 1413 | 1010 |
| 8740,84919 | 8561,84799 | 8489,71722 | 8434,08594 | 8415,09434 | 8407,38209 | 8320,11536 | 8244,86301 | 8179,63111 | 8103,72771 | 8032,12851 | 7977,24399 | 7975,74627 |

| Salicin | α-Cyclodextri | Acetamide | 4-Hydroxy Ber | Butyric Acid | L-Sorbose | Sorbic Acid | Arbutin | L-Phenylalani | D-Fucose | Mannan | 2-Hydroxy Be | Melibionic Ac |
|---|---|---|---|---|---|---|---|---|---|---|---|---|
| 0,0551 | 0,0523 | 0,0539 | 0,0511 | 0,053 | 0,0553 | 0,057 | 0,0559 | 0,0539 | 0,0541 | 0,0546 | 0,0529 | 0,0914 |
| 0,057 | 0,0537 | 0,0551 | 0,0521 | 0,0546 | 0,057 | 0,057 | 0,0577 | 0,0549 | 0,0559 | 0,0561 | 0,053 | 0,1307 |
| 0,0598 | 0,0573 | 0,0585 | 0,0563 | 0,0582 | 0,0595 | 0,0599 | 0,063 | 0,0585 | 0,0594 | 0,06 | 0,0548 | 0,0883 |
| 0,0641 | 0,0617 | 0,062 | 0,0613 | 0,0622 | 0,0632 | 0,0632 | 0,0681 | 0,0619 | 0,0635 | 0,0644 | 0,0565 | 0,1372 |
| 0,0676 | 0,0653 | 0,065 | 0,0652 | 0,0661 | 0,0662 | 0,0674 | 0,0719 | 0,0665 | 0,0677 | 0,0692 | 0,057 | 0,1749 |
| 0,0708 | 0,07 | 0,0674 | 0,0689 | 0,07 | 0,0687 | 0,0716 | 0,0747 | 0,0693 | 0,0717 | 0,0758 | 0,0586 | 0,0997 |
| 0,0739 | 0,0749 | 0,0715 | 0,0749 | 0,0741 | 0,0721 | 0,0774 | 0,079 | 0,0747 | 0,0746 | 0,0837 | 0,06 | 0,109 |
| 0,077 | 0,0813 | 0,0794 | 0,0776 | 0,0799 | 0,0767 | 0,0845 | 0,0842 | 0,0805 | 0,0803 | 0,0907 | 0,0614 | 0,1181 |
| 0,0879 | 0,0886 | 0,0897 | 0,0834 | 0,0889 | 0,0831 | 0,0931 | 0,0901 | 0,0886 | 0,0891 | 0,1024 | 0,0639 | 0,1292 |
| 0,095 | 0,1 | 0,0987 | 0,0956 | 0,0974 | 0,0947 | 0,1007 | 0,1013 | 0,098 | 0,0987 | 0,1085 | 0,0643 | 0,1503 |
| 0,1011 | 0,1055 | 0,1085 | 0,1063 | 0,1096 | 0,1037 | 0,1092 | 0,1133 | 0,1095 | 0,1095 | 0,1157 | 0,0666 | 0,1665 |
| 0,1071 | 0,1145 | 0,1136 | 0,1196 | 0,1155 | 0,1157 | 0,1205 | 0,1213 | 0,1184 | 0,118 | 0,1235 | 0,0668 | 0,3277 |
| 0,1186 | 0,1236 | 0,1295 | 0,1303 | 0,1259 | 0,1259 | 0,1277 | 0,1346 | 0,1323 | 0,1323 | 0,132 | 0,0675 | 0,3918 |
| 0,1279 | 0,132 | 0,1369 | 0,14 | 0,1329 | 0,1351 | 0,1429 | 0,1485 | 0,1426 | 0,1414 | 0,1438 | 0,0688 | 0,4108 |
| 0,1288 | 0,1346 | 0,1344 | 0,1502 | 0,1416 | 0,1445 | 0,151 | 0,1499 | 0,139 | 0,1397 | 0,1356 | 0,0712 | 0,3806 |
| 0,1341 | 0,1418 | 0,1389 | 0,162 | 0,1483 | 0,1475 | 0,1597 | 0,1604 | 0,1456 | 0,1511 | 0,1401 | 0,0716 | 0,3267 |
| 0,1348 | 0,1384 | 0,1441 | 0,1683 | 0,1522 | 0,1523 | 0,1596 | 0,1562 | 0,1492 | 0,1504 | 0,1448 | 0,0734 | 0,3335 |
| 0,1381 | 0,1453 | 0,1462 | 0,1728 | 0,1545 | 0,1582 | 0,1638 | 0,1635 | 0,156 | 0,154 | 0,1502 | 0,0747 | 0,343 |
| 0,1416 | 0,1447 | 0,1509 | 0,1778 | 0,1536 | 0,157 | 0,1672 | 0,1666 | 0,1598 | 0,156 | 0,154 | 0,0755 | 0,3962 |
| 0,1444 | 0,1434 | 0,1572 | 0,1827 | 0,1556 | 0,1646 | 0,173 | 0,1716 | 0,1617 | 0,1595 | 0,1502 | 0,0762 | 0,3849 |
| 0,1459 | 0,1486 | 0,1548 | 0,19 | 0,1504 | 0,1714 | 0,1712 | 0,1709 | 0,1601 | 0,1636 | 0,1523 | 0,0775 | 0,3752 |
| 0,1515 | 0,1437 | 0,163 | 0,1992 | 0,1547 | 0,1753 | 0,174 | 0,1753 | 0,1625 | 0,1586 | 0,1491 | 0,0779 | 0,3686 |
| 0,1564 | 0,1415 | 0,1681 | 0,1985 | 0,1536 | 0,187 | 0,1767 | 0,1806 | 0,1681 | 0,159 | 0,1477 | 0,0787 | 0,354 |
| 0,1588 | 0,1418 | 0,1709 | 0,2232 | 0,1604 | 0,194 | 0,1869 | 0,1852 | 0,1698 | 0,1595 | 0,149 | 0,0803 | 0,3588 |
| 0,1686 | 0,1423 | 0,1736 | 0,2352 | 0,1698 | 0,1964 | 0,1858 | 0,1855 | 0,1736 | 0,1627 | 0,1501 | 0,0817 | 0,359 |
| 0,1628 | 0,1435 | 0,1752 | 0,227 | 0,1709 | 0,2024 | 0,1908 | 0,1907 | 0,1742 | 0,1609 | 0,1551 | 0,0825 | 0,3396 |
| 0,1653 | 0,1478 | 0,1768 | 0,2215 | 0,1718 | 0,2012 | 0,1875 | 0,1843 | 0,176 | 0,1615 | 0,1544 | 0,0824 | 0,3421 |
| 0,167 | 0,1492 | 0,1781 | 0,2343 | 0,174 | 0,2059 | 0,1922 | 0,1889 | 0,1774 | 0,1646 | 0,1571 | 0,0832 | 0,3542 |
| 0,1713 | 0,1509 | 0,1814 | 0,2349 | 0,1727 | 0,2095 | 0,1976 | 0,193 | 0,1793 | 0,1686 | 0,1589 | 0,0839 | 0,338 |
| 0,1732 | 0,1539 | 0,1826 | 0,257 | 0,1715 | 0,2183 | 0,2004 | 0,1909 | 0,1794 | 0,1694 | 0,1564 | 0,0859 | 0,3594 |
| 0,176 | 0,1551 | 0,186 | 0,2529 | 0,1727 | 0,2204 | 0,2034 | 0,1953 | 0,1831 | 0,1721 | 0,1522 | 0,0867 | 0,3479 |
| 0,1796 | 0,1573 | 0,1849 | 0,2441 | 0,1712 | 0,2258 | 0,2155 | 0,1973 | 0,1835 | 0,1727 | 0,153 | 0,0872 | 0,3534 |
| 0,1828 | 0,1621 | 0,1862 | 0,2395 | 0,1724 | 0,2248 | 0,2211 | 0,1993 | 0,1853 | 0,1758 | 0,155 | 0,0891 | 0,3615 |
| 0,1854 | 0,1619 | 0,1857 | 0,2667 | 0,1712 | 0,2327 | 0,2171 | 0,1979 | 0,1865 | 0,1805 | 0,1534 | 0,0936 | 0,3752 |
| 0,1865 | 0,1619 | 0,1873 | 0,2735 | 0,1745 | 0,232 | 0,2166 | 0,2006 | 0,1853 | 0,1809 | 0,151 | 0,0938 | 0,3809 |
| 0,1885 | 0,1614 | 0,1871 | 0,2689 | 0,1752 | 0,2306 | 0,2156 | 0,2039 | 0,1863 | 0,1824 | 0,1468 | 0,095 | 0,3824 |

| Salicin | α-Cyclodextri | Acetamide | 4-Hydroxy Ber | Butyric Acid | L-Sorbose | Sorbic Acid | Arbutin | L-Phenylalani | D-Fucose | Mannan | 2-Hydroxy Be | Melibionic Ac |
|---|---|---|---|---|---|---|---|---|---|---|---|---|
| 150 | 161 | 175 | 168 | 168 | 165 | 326 | 172 | 172 | 165 | 175 | 159 | 180 |
| 159 | 157 | 174 | 167 | 166 | 162 | 333 | 167 | 167 | 154 | 179 | 160 | 215 |
| 159 | 168 | 172 | 177 | 173 | 166 | 339 | 173 | 176 | 159 | 177 | 156 | 183 |
| 172 | 180 | 180 | 192 | 179 | 179 | 355 | 193 | 188 | 170 | 193 | 158 | 234 |
| 192 | 186 | 201 | 215 | 194 | 188 | 367 | 203 | 211 | 184 | 206 | 162 | 285 |
| 203 | 207 | 212 | 236 | 209 | 194 | 377 | 219 | 218 | 192 | 221 | 166 | 225 |
| 218 | 221 | 227 | 251 | 217 | 202 | 399 | 239 | 238 | 200 | 240 | 175 | 248 |
| 242 | 253 | 254 | 267 | 230 | 222 | 418 | 261 | 262 | 218 | 271 | 181 | 283 |
| 276 | 281 | 278 | 297 | 252 | 241 | 440 | 287 | 288 | 256 | 300 | 186 | 362 |
| 293 | 299 | 298 | 343 | 287 | 267 | 472 | 310 | 317 | 271 | 333 | 193 | 451 |
| 323 | 340 | 333 | 373 | 331 | 291 | 501 | 360 | 353 | 299 | 377 | 196 | 512 |
| 353 | 378 | 359 | 394 | 365 | 311 | 534 | 385 | 386 | 333 | 432 | 213 | 837 |
| 404 | 447 | 448 | 431 | 410 | 351 | 553 | 460 | 463 | 392 | 491 | 215 | 971 |
| 443 | 529 | 484 | 482 | 486 | 405 | 614 | 528 | 521 | 455 | 565 | 221 | 930 |
| 507 | 605 | 568 | 514 | 572 | 449 | 655 | 585 | 572 | 506 | 636 | 229 | 891 |
| 550 | 661 | 630 | 583 | 639 | 503 | 717 | 640 | 625 | 567 | 700 | 240 | 797 |
| 581 | 732 | 698 | 664 | 730 | 559 | 772 | 693 | 682 | 627 | 756 | 246 | 827 |
| 638 | 787 | 763 | 724 | 786 | 614 | 828 | 735 | 731 | 688 | 817 | 262 | 922 |
| 664 | 815 | 804 | 802 | 848 | 658 | 873 | 755 | 765 | 733 | 859 | 267 | 1103 |
| 713 | 852 | 841 | 862 | 880 | 709 | 916 | 832 | 827 | 789 | 872 | 278 | 1139 |
| 730 | 869 | 887 | 940 | 899 | 760 | 951 | 863 | 857 | 805 | 883 | 284 | 1164 |
| 790 | 900 | 931 | 985 | 919 | 805 | 984 | 894 | 887 | 839 | 926 | 289 | 1211 |
| 814 | 943 | 974 | 1045 | 923 | 837 | 1036 | 930 | 912 | 860 | 943 | 289 | 1234 |
| 866 | 973 | 1005 | 1207 | 957 | 928 | 1072 | 975 | 951 | 890 | 970 | 320 | 1275 |
| 913 | 989 | 1043 | 1330 | 960 | 971 | 1108 | 993 | 980 | 927 | 1022 | 328 | 1361 |
| 955 | 1019 | 1068 | 1299 | 992 | 1011 | 1172 | 1021 | 996 | 971 | 1039 | 325 | 1344 |
| 982 | 1037 | 1075 | 1269 | 1004 | 1018 | 1158 | 1015 | 1018 | 998 | 1043 | 342 | 1437 |
| 1021 | 1075 | 1134 | 1401 | 1032 | 1080 | 1202 | 1036 | 1057 | 1025 | 1056 | 345 | 1491 |
| 1051 | 1090 | 1158 | 1405 | 1031 | 1091 | 1249 | 1059 | 1081 | 1048 | 1060 | 354 | 1515 |
| 1092 | 1120 | 1212 | 1558 | 1055 | 1182 | 1281 | 1078 | 1122 | 1076 | 1060 | 363 | 1623 |
| 1110 | 1107 | 1224 | 1565 | 1079 | 1198 | 1316 | 1110 | 1155 | 1077 | 1027 | 383 | 1648 |
| 1162 | 1115 | 1237 | 1525 | 1097 | 1269 | 1381 | 1139 | 1135 | 1080 | 993 | 394 | 1763 |
| 1180 | 1091 | 1231 | 1515 | 1108 | 1314 | 1415 | 1154 | 1152 | 1094 | 974 | 406 | 1839 |
| 1191 | 1084 | 1226 | 1723 | 1101 | 1337 | 1442 | 1158 | 1132 | 1093 | 922 | 424 | 1964 |
| 1210 | 1048 | 1218 | 1818 | 1108 | 1361 | 1438 | 1182 | 1120 | 1072 | 869 | 437 | 2039 |
| 1207 | 1015 | 1197 | 1797 | 1070 | 1431 | 1468 | 1206 | 1096 | 1060 | 816 | 447 | 2116 |
| 7923,53823 | 7827,68103 | 7672,67267 | 7479,33884 | 7381,34206 | 7221,90531 | 7200,50441 | 6986,48649 | 6978,85196 | 6975,83788 | 6952,27766 | 6840,85511 | 6652,92096 |

| L-Ornithine | Chondroitin S | D-ribose | N-Acetyl-DGal | Laminarin | Pectin | D-Raffinose | Itaconic Acid | MOPS no sug | Gentiobiose | D-glucose |
|---|---|---|---|---|---|---|---|---|---|---|
| 0,0536 | 0,0557 | 0,0531 | 0,053 | 0,0534 | 0,067 | 0,0537 | 0,0547 | 0,0542 | 0,0536 | 0,0597 |
| 0,055 | 0,0615 | 0,0539 | 0,0548 | 0,0554 | 0,0691 | 0,0555 | 0,0556 | 0,0568 | 0,0558 | 0,0594 |
| 0,0564 | 0,061 | 0,0573 | 0,0577 | 0,0609 | 0,072 | 0,0608 | 0,0593 | 0,058 | 0,0597 | 0,0623 |
| 0,0603 | 0,065 | 0,06 | 0,0618 | 0,0634 | 0,0763 | 0,0622 | 0,0629 | 0,0626 | 0,0629 | 0,0658 |
| 0,0639 | 0,0695 | 0,0646 | 0,0656 | 0,0676 | 0,081 | 0,0671 | 0,0651 | 0,0664 | 0,0676 | 0,0693 |
| 0,0667 | 0,0746 | 0,0676 | 0,0684 | 0,0711 | 0,0871 | 0,0709 | 0,0682 | 0,0695 | 0,0703 | 0,0745 |
| 0,0706 | 0,0814 | 0,0722 | 0,0729 | 0,0759 | 0,0965 | 0,0755 | 0,0722 | 0,0744 | 0,0748 | 0,08 |
| 0,0784 | 0,0904 | 0,0786 | 0,0787 | 0,0826 | 0,1029 | 0,0837 | 0,0772 | 0,0839 | 0,0822 | 0,0857 |
| 0,0888 | 0,098 | 0,0882 | 0,0896 | 0,092 | 0,1141 | 0,0962 | 0,0861 | 0,0888 | 0,0939 | 0,0951 |
| 0,0982 | 0,1054 | 0,0974 | 0,0952 | 0,1 | 0,1212 | 0,1061 | 0,0916 | 0,0986 | 0,1007 | 0,105 |
| 0,1079 | 0,1113 | 0,107 | 0,1021 | 0,1078 | 0,1281 | 0,1126 | 0,0956 | 0,0991 | 0,1083 | 0,1164 |
| 0,117 | 0,1243 | 0,1194 | 0,1062 | 0,1149 | 0,1352 | 0,1226 | 0,1077 | 0,1097 | 0,1147 | 0,1281 |
| 0,1267 | 0,1286 | 0,1271 | 0,1183 | 0,1244 | 0,1423 | 0,1329 | 0,1269 | 0,1151 | 0,1272 | 0,1384 |
| 0,1327 | 0,1277 | 0,1344 | 0,1304 | 0,1303 | 0,1468 | 0,1511 | 0,1321 | 0,1338 | 0,1378 | 0,1477 |
| 0,1358 | 0,1396 | 0,1371 | 0,1307 | 0,1284 | 0,1428 | 0,142 | 0,1547 | 0,1366 | 0,1487 | 0,1559 |
| 0,14 | 0,1379 | 0,1377 | 0,1296 | 0,1331 | 0,1449 | 0,1559 | 0,1323 | 0,1304 | 0,1451 | 0,1611 |
| 0,1419 | 0,1441 | 0,1448 | 0,1427 | 0,1478 | 0,1535 | 0,1645 | 0,1565 | 0,136 | 0,1478 | 0,1661 |
| 0,1457 | 0,1398 | 0,1468 | 0,135 | 0,1498 | 0,1463 | 0,1661 | 0,1508 | 0,1326 | 0,1488 | 0,1655 |
| 0,1471 | 0,1438 | 0,1492 | 0,1381 | 0,1417 | 0,1439 | 0,1797 | 0,1588 | 0,136 | 0,149 | 0,1673 |
| 0,1509 | 0,1409 | 0,1538 | 0,1368 | 0,1381 | 0,1451 | 0,1813 | 0,1672 | 0,133 | 0,1494 | 0,1663 |
| 0,1559 | 0,138 | 0,1562 | 0,1441 | 0,1389 | 0,1461 | 0,182 | 0,1642 | 0,1345 | 0,1522 | 0,1676 |
| 0,1559 | 0,1333 | 0,1578 | 0,1521 | 0,1378 | 0,1473 | 0,189 | 0,1683 | 0,1328 | 0,1548 | 0,1678 |
| 0,1599 | 0,1335 | 0,1583 | 0,1554 | 0,1368 | 0,1479 | 0,1917 | 0,1719 | 0,1319 | 0,1578 | 0,1688 |
| 0,159 | 0,1337 | 0,1595 | 0,1588 | 0,1413 | 0,1489 | 0,1974 | 0,1755 | 0,1344 | 0,1626 | 0,1693 |
| 0,159 | 0,1381 | 0,161 | 0,1664 | 0,1451 | 0,1498 | 0,2116 | 0,1759 | 0,1365 | 0,1705 | 0,1716 |
| 0,1589 | 0,138 | 0,1609 | 0,1686 | 0,1466 | 0,1489 | 0,2154 | 0,1764 | 0,1391 | 0,1773 | 0,17 |
| 0,1596 | 0,1382 | 0,1618 | 0,1699 | 0,1529 | 0,1472 | 0,2224 | 0,1812 | 0,1418 | 0,1831 | 0,1701 |
| 0,1616 | 0,1398 | 0,1636 | 0,172 | 0,1531 | 0,1443 | 0,2244 | 0,1821 | 0,1434 | 0,1865 | 0,1688 |
| 0,163 | 0,1414 | 0,166 | 0,1729 | 0,1535 | 0,1445 | 0,226 | 0,1826 | 0,1447 | 0,1916 | 0,1686 |
| 0,1632 | 0,1432 | 0,1673 | 0,1723 | 0,1521 | 0,1429 | 0,2269 | 0,1837 | 0,1482 | 0,1924 | 0,168 |
| 0,1641 | 0,1454 | 0,1692 | 0,1747 | 0,152 | 0,1353 | 0,2278 | 0,1834 | 0,1487 | 0,1948 | 0,168 |
| 0,1627 | 0,145 | 0,1695 | 0,172 | 0,1515 | 0,1329 | 0,2335 | 0,1841 | 0,1471 | 0,1955 | 0,1701 |
| 0,1611 | 0,1453 | 0,1691 | 0,1674 | 0,1508 | 0,1282 | 0,2375 | 0,1614 | 0,1448 | 0,1966 | 0,1713 |
| 0,1607 | 0,1428 | 0,1683 | 0,166 | 0,1489 | 0,1206 | 0,2372 | 0,1616 | 0,14 | 0,1963 | 0,1696 |
| 0,158 | 0,1416 | 0,1626 | 0,1638 | 0,1456 | 0,1122 | 0,2374 | 0,1619 | 0,1346 | 0,1956 | 0,1696 |
| 0,1548 | 0,1388 | 0,1606 | 0,1615 | 0,1418 | 0,1013 | 0,2396 | 0,1614 | 0,127 | 0,1936 | 0,169 |

| L-Ornithine | Chondroitin S | D-ribose | N-Acetyl-DGal | Laminarin | Pectin | D-Raffinose | Itaconic Acid | MOPS no sub | Gentiobiose | D-glucose |
|---|---|---|---|---|---|---|---|---|---|---|
| 165 | 167 | 178 | 149 | 148 | 174 | 154 | 145 | 176 | 151 | 486 |
| 167 | 162 | 173 | 153 | 153 | 175 | 154 | 141 | 175 | 148 | 487 |
| 165 | 169 | 178 | 150 | 153 | 178 | 153 | 147 | 176 | 155 | 502 |
| 179 | 176 | 188 | 161 | 166 | 179 | 166 | 159 | 183 | 164 | 513 |
| 198 | 191 | 207 | 175 | 187 | 193 | 185 | 178 | 198 | 176 | 515 |
| 213 | 214 | 215 | 188 | 199 | 223 | 186 | 199 | 207 | 193 | 526 |
| 227 | 230 | 232 | 197 | 215 | 243 | 199 | 209 | 223 | 204 | 538 |
| 242 | 253 | 248 | 226 | 241 | 267 | 214 | 225 | 256 | 237 | 545 |
| 271 | 282 | 286 | 249 | 272 | 299 | 243 | 261 | 272 | 268 | 566 |
| 293 | 305 | 307 | 269 | 286 | 335 | 255 | 278 | 300 | 276 | 585 |
| 325 | 324 | 348 | 276 | 330 | 376 | 278 | 294 | 331 | 308 | 602 |
| 377 | 380 | 395 | 316 | 367 | 436 | 304 | 369 | 379 | 349 | 618 |
| 445 | 432 | 474 | 373 | 428 | 523 | 353 | 420 | 443 | 408 | 665 |
| 503 | 500 | 522 | 428 | 489 | 586 | 400 | 403 | 513 | 457 | 706 |
| 573 | 521 | 600 | 480 | 561 | 676 | 439 | 528 | 568 | 478 | 748 |
| 638 | 608 | 641 | 526 | 608 | 731 | 479 | 541 | 611 | 523 | 778 |
| 697 | 672 | 716 | 585 | 674 | 792 | 512 | 597 | 675 | 556 | 847 |
| 747 | 718 | 774 | 617 | 698 | 833 | 562 | 596 | 740 | 598 | 888 |
| 768 | 737 | 800 | 644 | 734 | 871 | 594 | 673 | 753 | 636 | 909 |
| 798 | 768 | 841 | 670 | 759 | 883 | 621 | 735 | 765 | 651 | 939 |
| 819 | 756 | 866 | 714 | 755 | 878 | 654 | 760 | 793 | 681 | 975 |
| 829 | 811 | 888 | 701 | 776 | 889 | 675 | 774 | 808 | 709 | 983 |
| 866 | 829 | 908 | 744 | 804 | 917 | 698 | 822 | 844 | 737 | 1031 |
| 893 | 859 | 916 | 788 | 820 | 912 | 725 | 850 | 855 | 761 | 1046 |
| 915 | 891 | 949 | 798 | 835 | 908 | 774 | 862 | 888 | 785 | 1070 |
| 943 | 892 | 981 | 833 | 859 | 908 | 808 | 860 | 900 | 817 | 1039 |
| 957 | 909 | 996 | 856 | 872 | 878 | 839 | 885 | 890 | 824 | 1077 |
| 970 | 912 | 1021 | 881 | 876 | 863 | 855 | 888 | 909 | 839 | 1057 |
| 988 | 928 | 1041 | 896 | 890 | 843 | 906 | 865 | 902 | 844 | 1071 |
| 978 | 912 | 1036 | 897 | 875 | 796 | 939 | 855 | 872 | 847 | 1058 |
| 982 | 896 | 1043 | 929 | 860 | 763 | 970 | 845 | 837 | 864 | 1072 |
| 957 | 868 | 1024 | 927 | 830 | 703 | 1001 | 828 | 792 | 843 | 1064 |
| 928 | 845 | 988 | 914 | 803 | 627 | 1033 | 741 | 744 | 864 | 1057 |
| 890 | 812 | 948 | 887 | 759 | 550 | 1097 | 721 | 677 | 875 | 1051 |
| 848 | 754 | 891 | 851 | 720 | 469 | 1125 | 711 | 630 | 876 | 1045 |
| 814 | 695 | 846 | 823 | 682 | 368 | 1166 | 720 | 556 | 880 | 1016 |
| 6413,04348 | 6353,79061 | 6213,95349 | 6211,98157 | 6040,72398 | 5655,97668 | 5443,78698 | 5388,94096 | 5219,7802 | 5207,14286 | 4849,03934 |

| Lactitol | N-AcetylNeura | ß-Methyl-DGa | D-Tartaric Aci | ß-D-Allose | Dihydroxy Ac | Capric Acid | 2-Deoxy-DRib | Gelatin |
|---|---|---|---|---|---|---|---|---|
| 0,0551 | 0,0539 | 0,0551 | 0,0553 | 0,0547 | 0,0646 | 0,0563 | 0,0593 | 0,6105 |
| 0,0578 | 0,0559 | 0,0567 | 0,0561 | 0,0569 | 0,0642 | 0,0572 | 0,0605 | 0,6092 |
| 0,0617 | 0,0586 | 0,061 | 0,0591 | 0,0595 | 0,0654 | 0,0599 | 0,063 | 0,5519 |
| 0,0654 | 0,0624 | 0,0659 | 0,0625 | 0,0609 | 0,067 | 0,0625 | 0,0646 | 0,3053 |
| 0,0706 | 0,0666 | 0,0704 | 0,0669 | 0,0631 | 0,0701 | 0,0641 | 0,0666 | 0,0724 |
| 0,0745 | 0,0696 | 0,0754 | 0,069 | 0,0655 | 0,0731 | 0,0672 | 0,0682 | 0,0777 |
| 0,0777 | 0,0745 | 0,0805 | 0,0731 | 0,0674 | 0,0783 | 0,0709 | 0,0696 | 0,0848 |
| 0,0837 | 0,0792 | 0,0855 | 0,0778 | 0,0698 | 0,0817 | 0,0771 | 0,0712 | 0,0938 |
| 0,0938 | 0,0908 | 0,0966 | 0,0846 | 0,0736 | 0,0886 | 0,0847 | 0,0726 | 0,1039 |
| 0,1039 | 0,0978 | 0,1048 | 0,0956 | 0,0776 | 0,0933 | 0,0929 | 0,0732 | 0,1152 |
| 0,1142 | 0,1041 | 0,1194 | 0,1042 | 0,0835 | 0,105 | 0,1013 | 0,0762 | 0,1281 |
| 0,1209 | 0,1128 | 0,1345 | 0,1146 | 0,09 | 0,1149 | 0,1109 | 0,0765 | 0,1451 |
| 0,1341 | 0,1225 | 0,1456 | 0,1274 | 0,0984 | 0,1269 | 0,1187 | 0,0778 | 0,159 |
| 0,1457 | 0,1348 | 0,1565 | 0,1356 | 0,1112 | 0,1344 | 0,125 | 0,0792 | 0,1691 |
| 0,1567 | 0,145 | 0,1656 | 0,1414 | 0,1247 | 0,1499 | 0,1327 | 0,0813 | 0,1689 |
| 0,1661 | 0,1573 | 0,1723 | 0,1504 | 0,1345 | 0,1622 | 0,1349 | 0,0829 | 0,1733 |
| 0,1615 | 0,1598 | 0,1829 | 0,152 | 0,1506 | 0,167 | 0,129 | 0,085 | 0,1774 |
| 0,1644 | 0,1717 | 0,1923 | 0,1569 | 0,1622 | 0,1756 | 0,1266 | 0,0878 | 0,18 |
| 0,1669 | 0,179 | 0,1978 | 0,1578 | 0,1713 | 0,1797 | 0,1248 | 0,0914 | 0,1882 |
| 0,1715 | 0,1873 | 0,2147 | 0,163 | 0,1781 | 0,1856 | 0,1247 | 0,0944 | 0,1898 |
| 0,1816 | 0,1998 | 0,225 | 0,1602 | 0,1861 | 0,1888 | 0,1248 | 0,1013 | 0,1896 |
| 0,1858 | 0,1995 | 0,2411 | 0,1668 | 0,1956 | 0,1931 | 0,1259 | 0,1074 | 0,1927 |
| 0,1804 | 0,2079 | 0,2478 | 0,1698 | 0,2058 | 0,1999 | 0,1252 | 0,1166 | 0,1985 |
| 0,1862 | 0,21 | 0,2675 | 0,1723 | 0,2291 | 0,1955 | 0,1241 | 0,1223 | 0,202 |
| 0,1871 | 0,217 | 0,2796 | 0,1756 | 0,2397 | 0,2011 | 0,1243 | 0,1319 | 0,2064 |
| 0,1828 | 0,2211 | 0,292 | 0,1784 | 0,2503 | 0,2005 | 0,1239 | 0,1425 | 0,2133 |
| 0,1766 | 0,2306 | 0,293 | 0,1821 | 0,2614 | 0,2011 | 0,1234 | 0,1444 | 0,217 |
| 0,1759 | 0,238 | 0,2945 | 0,1854 | 0,2773 | 0,2011 | 0,1223 | 0,154 | 0,2182 |
| 0,1758 | 0,2473 | 0,3019 | 0,1902 | 0,2823 | 0,2028 | 0,1218 | 0,1584 | 0,2164 |
| 0,1772 | 0,2476 | 0,3062 | 0,1934 | 0,2967 | 0,2011 | 0,1219 | 0,1568 | 0,2172 |
| 0,1806 | 0,2518 | 0,3088 | 0,1931 | 0,2988 | 0,2025 | 0,1204 | 0,1575 | 0,2148 |
| 0,1804 | 0,2512 | 0,3154 | 0,1934 | 0,3053 | 0,2029 | 0,1197 | 0,1606 | 0,2119 |
| 0,1823 | 0,2546 | 0,3143 | 0,1952 | 0,3144 | 0,2041 | 0,119 | 0,1605 | 0,2074 |
| 0,1833 | 0,2556 | 0,3149 | 0,2028 | 0,3195 | 0,2065 | 0,1187 | 0,1623 | 0,2036 |
| 0,1873 | 0,253 | 0,3139 | 0,1977 | 0,3229 | 0,208 | 0,1179 | 0,1643 | 0,2002 |
| 0,1879 | 0,2534 | 0,319 | 0,1992 | 0,3263 | 0,2094 | 0,1174 | 0,1616 | 0,1962 |

| Lactitol | N-AcetylNeura | ß-Methyl-DGa | D-Tartaric Aci | ß-D-Allose | Dihydroxy Ac | Capric Acid | 2-Deoxy-DRib | Gelatin |
|---|---|---|---|---|---|---|---|---|
| 155 | 148 | 153 | 178 | 160 | 2218 | 158 | 2072 | 227 |
| 149 | 146 | 155 | 177 | 155 | 2230 | 159 | 1893 | 215 |
| 152 | 152 | 161 | 179 | 160 | 2273 | 162 | 1839 | 208 |
| 167 | 161 | 174 | 188 | 161 | 2295 | 164 | 1806 | 220 |
| 173 | 163 | 177 | 195 | 159 | 2310 | 161 | 1766 | 202 |
| 175 | 176 | 197 | 203 | 169 | 2293 | 168 | 1743 | 219 |
| 170 | 191 | 207 | 216 | 171 | 2331 | 165 | 1751 | 243 |
| 174 | 208 | 223 | 224 | 175 | 2303 | 168 | 1768 | 264 |
| 201 | 235 | 254 | 248 | 188 | 2315 | 175 | 1771 | 295 |
| 210 | 248 | 284 | 276 | 197 | 2285 | 183 | 1758 | 328 |
| 222 | 275 | 320 | 296 | 207 | 2286 | 192 | 1758 | 375 |
| 234 | 302 | 365 | 329 | 222 | 2273 | 204 | 1774 | 434 |
| 275 | 372 | 416 | 374 | 246 | 2291 | 224 | 1757 | 507 |
| 302 | 429 | 494 | 445 | 271 | 2281 | 237 | 1761 | 590 |
| 334 | 485 | 562 | 492 | 299 | 2294 | 244 | 1742 | 672 |
| 356 | 550 | 614 | 561 | 317 | 2298 | 259 | 1757 | 764 |
| 379 | 593 | 670 | 590 | 355 | 2336 | 260 | 1748 | 828 |
| 402 | 629 | 725 | 633 | 403 | 2346 | 265 | 1728 | 886 |
| 422 | 671 | 778 | 664 | 440 | 2372 | 266 | 1757 | 942 |
| 442 | 714 | 849 | 671 | 508 | 2404 | 275 | 1773 | 1000 |
| 476 | 768 | 930 | 680 | 540 | 2419 | 282 | 1778 | 1059 |
| 478 | 815 | 1018 | 681 | 595 | 2407 | 283 | 1762 | 1102 |
| 511 | 824 | 1063 | 688 | 634 | 2457 | 287 | 1795 | 1138 |
| 521 | 861 | 1144 | 703 | 703 | 2471 | 293 | 1815 | 1181 |
| 527 | 879 | 1197 | 701 | 782 | 2502 | 286 | 1837 | 1195 |
| 560 | 910 | 1222 | 686 | 834 | 2533 | 293 | 1867 | 1197 |
| 578 | 942 | 1253 | 682 | 899 | 2547 | 300 | 1888 | 1183 |
| 596 | 956 | 1238 | 671 | 946 | 2558 | 296 | 1919 | 1173 |
| 627 | 963 | 1252 | 671 | 971 | 2574 | 303 | 1939 | 1152 |
| 644 | 970 | 1234 | 679 | 1018 | 2569 | 302 | 1986 | 1099 |
| 658 | 945 | 1204 | 669 | 1030 | 2572 | 294 | 2012 | 1067 |
| 698 | 936 | 1191 | 670 | 1018 | 2573 | 302 | 2009 | 1019 |
| 718 | 930 | 1163 | 661 | 1025 | 2587 | 297 | 2025 | 979 |
| 734 | 911 | 1127 | 665 | 1021 | 2595 | 294 | 2033 | 928 |
| 746 | 927 | 1104 | 662 | 1003 | 2587 | 300 | 2028 | 880 |
| 760 | 919 | 1107 | 654 | 972 | 2569 | 293 | 1996 | 850 |
| 4555,72289 | 3864,66165 | 3615,00568 | 3307,85268 | 2989,69072 | 2424,03315 | 2209,49264 | -742,913 | -1503,7413 |

182 **Plate PM3B, nucleosides and nucleotides:**

| Time (sec) | Uridine | Inosine | Guanosine | Cytidine | Adenosine | Thymidine | Xanthosine | Guanine | Xanthine | MOPS no sugar | Cytosine | Uracil | Thymine | Adenine | D-Glucose |
|---|---|---|---|---|---|---|---|---|---|---|---|---|---|---|---|
| 0s | 0,0896 | 0,0899 | 0,0921 | 0,0896 | 0,0914 | 0,091 | 0,0943 | 0,1393 | 0,1156 | 0,088 | 0,0908 | 0,096 | 0,0931 | 0,0902 | 0,0907 |
| 600s | 0,0898 | 0,0899 | 0,0906 | 0,0901 | 0,092 | 0,0912 | 0,0943 | 0,1402 | 0,1178 | 0,0879 | 0,0912 | 0,0938 | 0,0939 | 0,0908 | 0,0912 |
| 1200s | 0,0899 | 0,0903 | 0,0892 | 0,09 | 0,0924 | 0,091 | 0,0951 | 0,1245 | 0,1233 | 0,088 | 0,092 | 0,0912 | 0,0939 | 0,0907 | 0,0914 |
| 1800s | 0,0909 | 0,0915 | 0,09 | 0,0913 | 0,0928 | 0,0918 | 0,0952 | 0,1278 | 0,1171 | 0,0887 | 0,0926 | 0,0914 | 0,095 | 0,0912 | 0,0926 |
| 2400s | 0,091 | 0,092 | 0,091 | 0,0927 | 0,0936 | 0,0924 | 0,0963 | 0,1277 | 0,1228 | 0,0899 | 0,0931 | 0,0908 | 0,0962 | 0,0917 | 0,0937 |
| 3000s | 0,0935 | 0,0938 | 0,0928 | 0,0934 | 0,0948 | 0,0943 | 0,0975 | 0,1315 | 0,1206 | 0,0916 | 0,0944 | 0,0926 | 0,0976 | 0,092 | 0,096 |
| 3600s | 0,0957 | 0,0954 | 0,0944 | 0,0956 | 0,0964 | 0,0956 | 0,0981 | 0,1263 | 0,122 | 0,093 | 0,0953 | 0,0935 | 0,0987 | 0,0931 | 0,0977 |
| 4200s | 0,0994 | 0,098 | 0,0964 | 0,0976 | 0,098 | 0,0978 | 0,0992 | 0,1385 | 0,1267 | 0,0936 | 0,0958 | 0,0951 | 0,0993 | 0,0941 | 0,1018 |
| 4801s | 0,1027 | 0,1004 | 0,0998 | 0,1015 | 0,1003 | 0,1006 | 0,101 | 0,1334 | 0,1191 | 0,0955 | 0,0978 | 0,0966 | 0,1011 | 0,0957 | 0,107 |
| 5401s | 0,1039 | 0,1019 | 0,0998 | 0,1023 | 0,1009 | 0,1003 | 0,1016 | 0,1421 | 0,1178 | 0,0975 | 0,0983 | 0,0973 | 0,1013 | 0,0954 | 0,1105 |
| 6001s | 0,104 | 0,102 | 0,1015 | 0,1037 | 0,1026 | 0,1016 | 0,1018 | 0,1265 | 0,1162 | 0,0991 | 0,0997 | 0,0992 | 0,103 | 0,0968 | 0,113 |
| 6601s | 0,1046 | 0,104 | 0,1025 | 0,1047 | 0,104 | 0,1033 | 0,1039 | 0,1457 | 0,1205 | 0,0996 | 0,1009 | 0,0992 | 0,1039 | 0,0969 | 0,1182 |
| 7201s | 0,1051 | 0,1053 | 0,104 | 0,1081 | 0,1068 | 0,1045 | 0,1051 | 0,1301 | 0,1201 | 0,1031 | 0,1031 | 0,1013 | 0,1066 | 0,0987 | 0,1201 |
| 7801s | 0,1089 | 0,1077 | 0,1058 | 0,1102 | 0,1088 | 0,1072 | 0,1071 | 0,1345 | 0,1184 | 0,1054 | 0,1054 | 0,1021 | 0,1083 | 0,1012 | 0,1247 |
| 8401s | 0,1141 | 0,1108 | 0,1086 | 0,1161 | 0,1128 | 0,1101 | 0,1107 | 0,1481 | 0,1183 | 0,108 | 0,1081 | 0,1056 | 0,1106 | 0,1025 | 0,1285 |
| 9001s | 0,1221 | 0,115 | 0,1132 | 0,1226 | 0,1189 | 0,1153 | 0,1144 | 0,1399 | 0,1213 | 0,1117 | 0,1109 | 0,1084 | 0,1143 | 0,1039 | 0,1419 |
| 9601s | 0,1322 | 0,1245 | 0,1215 | 0,1328 | 0,1272 | 0,1224 | 0,1199 | 0,1449 | 0,1362 | 0,1157 | 0,117 | 0,1123 | 0,1184 | 0,1074 | 0,1553 |
| 10201s | 0,148 | 0,137 | 0,1333 | 0,1479 | 0,138 | 0,1336 | 0,125 | 0,1484 | 0,1395 | 0,1235 | 0,1226 | 0,1191 | 0,1235 | 0,1115 | 0,1721 |
| 10801s | 0,1616 | 0,1518 | 0,1445 | 0,1672 | 0,1527 | 0,1475 | 0,1342 | 0,1577 | 0,1404 | 0,1365 | 0,1319 | 0,127 | 0,1326 | 0,117 | 0,1894 |
| 11401s | 0,1701 | 0,1654 | 0,1664 | 0,1838 | 0,1689 | 0,1654 | 0,1497 | 0,1722 | 0,1481 | 0,1499 | 0,146 | 0,138 | 0,1464 | 0,1228 | 0,2053 |
| 12001s | 0,1777 | 0,1742 | 0,1676 | 0,2002 | 0,1799 | 0,1742 | 0,1689 | 0,1866 | 0,1713 | 0,15 | 0,1673 | 0,1552 | 0,1639 | 0,1301 | 0,2267 |
| 12601s | 0,1885 | 0,1862 | 0,1812 | 0,2213 | 0,1893 | 0,1791 | 0,1636 | 0,1889 | 0,1723 | 0,161 | 0,1605 | 0,1625 | 0,1622 | 0,1419 | 0,2464 |
| 13201s | 0,1934 | 0,1936 | 0,1892 | 0,231 | 0,2021 | 0,1951 | 0,1793 | 0,2047 | 0,1904 | 0,1714 | 0,1781 | 0,1699 | 0,1712 | 0,1472 | 0,2626 |
| 13802s | 0,1993 | 0,2001 | 0,1918 | 0,2459 | 0,2095 | 0,2023 | 0,1839 | 0,2183 | 0,1957 | 0,1765 | 0,1832 | 0,1794 | 0,1837 | 0,1655 | 0,2816 |
| 14402s | 0,2082 | 0,2089 | 0,2016 | 0,251 | 0,2172 | 0,2095 | 0,1856 | 0,2104 | 0,1944 | 0,1794 | 0,1855 | 0,1836 | 0,1858 | 0,1711 | 0,2983 |
| 15002s | 0,2111 | 0,2122 | 0,2091 | 0,262 | 0,2264 | 0,2168 | 0,1893 | 0,2158 | 0,198 | 0,183 | 0,1903 | 0,1863 | 0,189 | 0,175 | 0,3148 |
| 15602s | 0,2193 | 0,2202 | 0,2203 | 0,2725 | 0,2367 | 0,2259 | 0,1974 | 0,2399 | 0,2045 | 0,1901 | 0,1968 | 0,1888 | 0,1914 | 0,1796 | 0,3191 |
| 16202s | 0,228 | 0,2295 | 0,2273 | 0,2873 | 0,2454 | 0,2336 | 0,2035 | 0,2405 | 0,2101 | 0,1961 | 0,2024 | 0,1935 | 0,1974 | 0,184 | 0,3279 |
| 16802s | 0,2357 | 0,2336 | 0,2353 | 0,2957 | 0,2547 | 0,2432 | 0,21 | 0,2508 | 0,2148 | 0,2001 | 0,2088 | 0,1999 | 0,2034 | 0,1859 | 0,3328 |
| 17402s | 0,2404 | 0,2416 | 0,2365 | 0,2946 | 0,2608 | 0,246 | 0,2133 | 0,2492 | 0,2187 | 0,204 | 0,2106 | 0,205 | 0,2072 | 0,1878 | 0,3383 |
| 18002s | 0,2486 | 0,2473 | 0,2417 | 0,2995 | 0,2733 | 0,2565 | 0,2177 | 0,2602 | 0,2222 | 0,2077 | 0,2144 | 0,2075 | 0,2123 | 0,1942 | 0,3435 |
| 18602s | 0,2555 | 0,2568 | 0,2471 | 0,3031 | 0,2792 | 0,266 | 0,2246 | 0,2536 | 0,2245 | 0,2122 | 0,2194 | 0,2107 | 0,2136 | 0,2006 | 0,3425 |
| 19202s | 0,2612 | 0,2628 | 0,2489 | 0,3065 | 0,2834 | 0,2716 | 0,2289 | 0,2621 | 0,2305 | 0,2149 | 0,2237 | 0,216 | 0,2183 | 0,2037 | 0,3297 |
| 19802s | 0,2691 | 0,2702 | 0,2526 | 0,3094 | 0,2917 | 0,2802 | 0,2316 | 0,262 | 0,2327 | 0,2192 | 0,2279 | 0,2215 | 0,2237 | 0,2083 | 0,3368 |
| 20402s | 0,2744 | 0,2775 | 0,2569 | 0,3127 | 0,3009 | 0,2912 | 0,2378 | 0,2654 | 0,2393 | 0,22 | 0,2313 | 0,2258 | 0,2273 | 0,211 | 0,3379 |
| 21002s | 0,2811 | 0,2807 | 0,2624 | 0,3171 | 0,3069 | 0,2961 | 0,2401 | 0,2687 | 0,2443 | 0,2256 | 0,2381 | 0,2309 | 0,2336 | 0,2144 | 0,3342 |
| 21602s | 0,2854 | 0,2907 | 0,2629 | 0,3188 | 0,3182 | 0,2962 | 0,2463 | 0,2809 | 0,256 | 0,2288 | 0,2397 | 0,2322 | 0,2344 | 0,2195 | 0,3333 |
| 22202s | 0,2942 | 0,2996 | 0,2666 | 0,3227 | 0,3293 | 0,3009 | 0,2501 | 0,2959 | 0,2556 | 0,2329 | 0,2439 | 0,2393 | 0,2391 | 0,2231 | 0,3319 |
| 22802s | 0,2935 | 0,3021 | 0,2707 | 0,3259 | 0,3312 | 0,3021 | 0,2541 | 0,2848 | 0,2555 | 0,2353 | 0,2481 | 0,2406 | 0,2413 | 0,2287 | 0,3233 |
| 23403s | 0,2971 | 0,3047 | 0,2712 | 0,3299 | 0,3331 | 0,304 | 0,2584 | 0,3 | 0,2652 | 0,2417 | 0,2517 | 0,2424 | 0,245 | 0,2327 | 0,3243 |
| 24003s | 0,3058 | 0,3126 | 0,2767 | 0,334 | 0,3374 | 0,3081 | 0,265 | 0,2842 | 0,2723 | 0,2456 | 0,2554 | 0,2484 | 0,2499 | 0,236 | 0,3252 |
| 24603s | 0,3108 | 0,3161 | 0,2788 | 0,3377 | 0,3362 | 0,3119 | 0,266 | 0,2938 | 0,2804 | 0,2455 | 0,2581 | 0,2516 | 0,254 | 0,2386 | 0,3238 |
| 25203s | 0,3125 | 0,3225 | 0,2817 | 0,34 | 0,3388 | 0,3123 | 0,2691 | 0,2841 | 0,2762 | 0,2496 | 0,2618 | 0,2573 | 0,258 | 0,2424 | 0,3169 |
| 25803s | 0,3191 | 0,3279 | 0,2819 | 0,3448 | 0,3441 | 0,3149 | 0,2737 | 0,2888 | 0,2768 | 0,252 | 0,2643 | 0,2573 | 0,2588 | 0,2474 | 0,3191 |
| 26403s | 0,3246 | 0,3379 | 0,2849 | 0,3492 | 0,3476 | 0,3204 | 0,2752 | 0,3005 | 0,2738 | 0,2541 | 0,2668 | 0,2598 | 0,262 | 0,2516 | 0,3179 |
| 27003s | 0,3271 | 0,3391 | 0,2862 | 0,3468 | 0,348 | 0,318 | 0,2728 | 0,298 | 0,2816 | 0,2567 | 0,2679 | 0,2629 | 0,2641 | 0,2533 | 0,3132 |
| 27603s | 0,333 | 0,3433 | 0,2933 | 0,355 | 0,3551 | 0,3265 | 0,2844 | 0,3067 | 0,2889 | 0,2594 | 0,2743 | 0,2694 | 0,2701 | 0,2607 | 0,3168 |
| 28203s | 0,3338 | 0,3469 | 0,2926 | 0,3566 | 0,3535 | 0,3273 | 0,2862 | 0,2969 | 0,2901 | 0,2619 | 0,2741 | 0,2679 | 0,2693 | 0,2617 | 0,3142 |
| 28803s | 0,3405 | 0,3468 | 0,2926 | 0,3561 | 0,3548 | 0,3272 | 0,282 | 0,3106 | 0,2861 | 0,2632 | 0,2766 | 0,2721 | 0,2732 | 0,2638 | 0,3147 |
| 29403s | 0,3439 | 0,3507 | 0,2968 | 0,3608 | 0,3597 | 0,3294 | 0,2895 | 0,3129 | 0,2875 | 0,2654 | 0,28 | 0,2721 | 0,2744 | 0,2677 | 0,316 |
| 30003s | 0,3437 | 0,3533 | 0,2993 | 0,3617 | 0,3638 | 0,3313 | 0,2879 | 0,3017 | 0,2859 | 0,2656 | 0,2816 | 0,2765 | 0,2767 | 0,2701 | 0,3068 |
| 30603s | 0,3482 | 0,3539 | 0,3 | 0,3628 | 0,3643 | 0,3322 | 0,29 | 0,3059 | 0,2891 | 0,2668 | 0,2826 | 0,278 | 0,2801 | 0,2736 | 0,3147 |
| 31203s | 0,3532 | 0,3583 | 0,3025 | 0,3665 | 0,3682 | 0,3363 | 0,295 | 0,3248 | 0,2893 | 0,2703 | 0,2842 | 0,2821 | 0,2818 | 0,2772 | 0,3164 |
| 31803s | 0,3536 | 0,3596 | 0,3036 | 0,3646 | 0,3654 | 0,3359 | 0,2916 | 0,3203 | 0,2852 | 0,2701 | 0,2854 | 0,283 | 0,2838 | 0,2788 | 0,3108 |
| 32404s | 0,3598 | 0,3635 | 0,3064 | 0,3689 | 0,369 | 0,3417 | 0,2934 | 0,328 | 0,2878 | 0,271 | 0,2871 | 0,2862 | 0,287 | 0,2819 | 0,3112 |

| gfp | Uridine | Inosine | Guanosine | Cytidine | Adenosine | Thymidine | Xanthosine | Guanine | Xanthine | MOPS no substra | Cytosine | Uracil | Thymine | Adenine | Glucose |
|---|---|---|---|---|---|---|---|---|---|---|---|---|---|---|---|
| 0s | 91 | 86 | 85 | 89 | 87 | 88 | 82 | 91 | 87 | 86 | 84 | 100 | 87 | 88 | 87 |
| 600s | 88 | 87 | 87 | 89 | 85 | 85 | 79 | 99 | 85 | 86 | 87 | 88 | 86 | 83 | 86 |
| 1200s | 87 | 87 | 84 | 88 | 84 | 86 | 83 | 86 | 89 | 84 | 88 | 90 | 90 | 87 | 81 |
| 1800s | 86 | 87 | 86 | 88 | 84 | 88 | 82 | 88 | 87 | 87 | 88 | 90 | 85 | 86 | 85 |
| 2400s | 87 | 87 | 84 | 89 | 83 | 87 | 83 | 95 | 92 | 87 | 89 | 93 | 87 | 88 | 82 |
| 3000s | 91 | 90 | 88 | 86 | 85 | 89 | 83 | 93 | 89 | 92 | 88 | 91 | 89 | 87 | 87 |
| 3600s | 94 | 93 | 91 | 90 | 88 | 91 | 85 | 93 | 92 | 92 | 91 | 89 | 92 | 91 | 87 |
| 4200s | 95 | 93 | 93 | 91 | 89 | 90 | 87 | 97 | 95 | 92 | 96 | 94 | 90 | 93 | 87 |
| 4800s | 95 | 92 | 95 | 89 | 90 | 91 | 91 | 98 | 92 | 94 | 94 | 95 | 91 | 94 | 86 |
| 5401s | 95 | 88 | 97 | 94 | 91 | 94 | 88 | 100 | 91 | 92 | 94 | 95 | 97 | 93 | 89 |
| 6001s | 96 | 91 | 97 | 94 | 91 | 93 | 89 | 95 | 95 | 92 | 96 | 94 | 92 | 93 | 87 |
| 6601s | 98 | 92 | 97 | 93 | 88 | 90 | 89 | 114 | 97 | 97 | 97 | 98 | 95 | 92 | 86 |
| 7201s | 98 | 90 | 103 | 95 | 87 | 89 | 89 | 96 | 96 | 95 | 97 | 94 | 93 | 96 | 88 |
| 7801s | 98 | 90 | 103 | 95 | 93 | 94 | 92 | 102 | 93 | 96 | 98 | 94 | 96 | 94 | 87 |
| 8401s | 103 | 98 | 107 | 99 | 95 | 100 | 97 | 116 | 101 | 100 | 100 | 98 | 98 | 91 | 88 |
| 9001s | 117 | 105 | 121 | 109 | 107 | 110 | 104 | 110 | 110 | 109 | 112 | 109 | 109 | 105 | 99 |
| 9601s | 125 | 116 | 130 | 118 | 112 | 115 | 112 | 114 | 115 | 113 | 114 | 113 | 111 | 100 | 104 |
| 10201s | 136 | 123 | 138 | 128 | 117 | 116 | 116 | 117 | 122 | 114 | 112 | 110 | 111 | 101 | 100 |
| 10801s | 168 | 144 | 159 | 140 | 129 | 136 | 124 | 131 | 132 | 127 | 132 | 123 | 120 | 110 | 119 |
| 11401s | 184 | 159 | 171 | 149 | 133 | 143 | 140 | 146 | 134 | 131 | 138 | 124 | 133 | 105 | 124 |
| 12001s | 202 | 171 | 198 | 163 | 157 | 151 | 153 | 143 | 159 | 149 | 146 | 142 | 144 | 109 | 133 |
| 12601s | 223 | 198 | 239 | 192 | 182 | 174 | 160 | 172 | 174 | 168 | 160 | 162 | 163 | 121 | 152 |
| 13201s | 243 | 218 | 259 | 220 | 198 | 194 | 179 | 186 | 192 | 182 | 171 | 172 | 166 | 128 | 166 |
| 13801s | 286 | 256 | 302 | 260 | 230 | 224 | 212 | 218 | 232 | 209 | 193 | 188 | 183 | 141 | 196 |
| 14402s | 315 | 279 | 342 | 292 | 258 | 254 | 223 | 227 | 224 | 216 | 212 | 206 | 197 | 152 | 214 |
| 15002s | 329 | 313 | 380 | 319 | 286 | 275 | 242 | 244 | 226 | 226 | 219 | 208 | 197 | 146 | 217 |
| 15602s | 362 | 358 | 412 | 370 | 313 | 310 | 256 | 284 | 267 | 244 | 230 | 217 | 209 | 150 | 238 |
| 16202s | 407 | 387 | 462 | 418 | 354 | 347 | 280 | 302 | 276 | 261 | 259 | 238 | 236 | 160 | 264 |
| 16802s | 432 | 410 | 505 | 455 | 389 | 377 | 308 | 326 | 302 | 275 | 280 | 259 | 246 | 168 | 269 |
| 17402s | 473 | 457 | 537 | 499 | 413 | 399 | 310 | 360 | 308 | 273 | 283 | 271 | 260 | 165 | 267 |
| 18002s | 494 | 477 | 570 | 543 | 445 | 429 | 339 | 361 | 320 | 296 | 293 | 272 | 270 | 165 | 266 |
| 18602s | 552 | 516 | 612 | 583 | 487 | 457 | 363 | 424 | 348 | 323 | 314 | 292 | 281 | 183 | 280 |
| 19202s | 620 | 563 | 674 | 641 | 530 | 510 | 394 | 422 | 374 | 335 | 333 | 315 | 299 | 185 | 278 |
| 19802s | 651 | 583 | 711 | 668 | 559 | 525 | 408 | 447 | 389 | 352 | 351 | 325 | 316 | 186 | 281 |
| 20402s | 695 | 630 | 753 | 719 | 611 | 571 | 429 | 457 | 396 | 372 | 367 | 335 | 328 | 198 | 281 |
| 21002s | 761 | 674 | 801 | 757 | 645 | 595 | 451 | 486 | 443 | 391 | 382 | 353 | 350 | 204 | 277 |
| 21602s | 814 | 710 | 845 | 800 | 715 | 627 | 472 | 508 | 462 | 396 | 405 | 365 | 363 | 207 | 280 |
| 22202s | 849 | 755 | 872 | 851 | 769 | 660 | 497 | 542 | 464 | 424 | 409 | 380 | 371 | 212 | 287 |
| 22802s | 902 | 813 | 928 | 895 | 823 | 702 | 518 | 546 | 470 | 428 | 430 | 389 | 384 | 208 | 276 |
| 23402s | 969 | 848 | 938 | 937 | 885 | 724 | 541 | 562 | 497 | 459 | 448 | 415 | 409 | 221 | 282 |
| 24003s | 1024 | 896 | 975 | 986 | 929 | 771 | 568 | 580 | 523 | 465 | 454 | 415 | 408 | 220 | 277 |
| 24603s | 1078 | 961 | 1011 | 1042 | 992 | 796 | 583 | 591 | 531 | 482 | 452 | 440 | 416 | 224 | 272 |
| 25203s | 1156 | 1010 | 1050 | 1079 | 1030 | 835 | 622 | 617 | 548 | 498 | 491 | 448 | 428 | 231 | 273 |
| 25803s | 1247 | 1096 | 1066 | 1138 | 1097 | 860 | 645 | 611 | 572 | 516 | 501 | 475 | 447 | 245 | 284 |
| 26403s | 1301 | 1154 | 1104 | 1179 | 1143 | 873 | 659 | 634 | 571 | 527 | 525 | 477 | 455 | 243 | 272 |
| 27003s | 1390 | 1268 | 1128 | 1229 | 1207 | 937 | 702 | 643 | 600 | 533 | 530 | 492 | 477 | 250 | 279 |
| 27603s | 1463 | 1345 | 1154 | 1267 | 1236 | 942 | 711 | 682 | 615 | 551 | 531 | 507 | 488 | 252 | 280 |
| 28203s | 1541 | 1381 | 1164 | 1294 | 1275 | 970 | 752 | 720 | 612 | 559 | 540 | 500 | 490 | 263 | 274 |
| 28803s | 1616 | 1474 | 1203 | 1356 | 1338 | 1009 | 780 | 705 | 627 | 560 | 564 | 526 | 505 | 270 | 282 |
| 29403s | 1699 | 1547 | 1231 | 1390 | 1350 | 1034 | 786 | 750 | 623 | 576 | 578 | 527 | 519 | 277 | 276 |
| 30003s | 1793 | 1566 | 1241 | 1430 | 1398 | 1059 | 809 | 773 | 612 | 570 | 577 | 535 | 520 | 276 | 283 |
| 30603s | 1905 | 1637 | 1255 | 1456 | 1436 | 1065 | 822 | 768 | 638 | 589 | 596 | 556 | 542 | 291 | 280 |
| 31203s | 1964 | 1695 | 1268 | 1473 | 1453 | 1084 | 836 | 748 | 623 | 585 | 598 | 565 | 544 | 290 | 281 |
| 31803s | 2108 | 1746 | 1292 | 1506 | 1502 | 1109 | 839 | 758 | 629 | 590 | 609 | 579 | 563 | 295 | 282 |
| 32403s | 2145 | 1792 | 1300 | 1535 | 1518 | 1129 | 860 | 773 | 638 | 597 | 610 | 582 | 563 | 295 | 272 |
| ratio | 7601,77646 | 6235,38012 | 5669,62203 | 5177,22879 | 5154,89914 | 4152,37335 | 3907,58413 | 3614,20244 | 3199,76771 | 2792,349727 | 2679,57208 | 2534,17455 | 2454,87365 | 1079,81221 | 839,002268 |

183

**Table S3**: Strains, plasmids and primers

| Strains | Number | Genotype of interest | Origin |
|---|---|---|---|
| Escherichia coli K12 subs. MG1655 | C349 | WT | Lab collection |
| Vibrio cholerae N16961 | 7805 | WT | Lab collection |
| Pseudomonas aeruginosa PAO1 | N065 | WT | Lab collection |
| Klebsiella pneumoniae NTUH K2044 | N105 | WT | Liver abscess, [68] |
| Acinetobacter baumannii ATCC 19606 | I276 | WT | Lab collection |
| Deletions in E.coli : | | | |
| Δcrp | 6570 | | |
| Δcra | P940 | | This study |
| ΔcmtA | OO56 | | This study |
| ΔfruA | N812 | | This study |
| ΔchbC | OO66 | | This study |
| ΔlamB | N810 | | This study |
| ΔmalE | N811 | | This study |
| ΔfrwB | N813 | | This study |
| ΔfrvB | N814 | | This study |
| ΔptsG | N815 | | This study |
| ΔtreB | N816 | | This study |
| ΔmalX | N818 | | This study |
| ΔmtlA | N819 | | This study |
| ΔmglA | N820 | | This study |
| ΔmngA | N821 | | This study |
| ΔypdG | N822 | | This study |
| ΔsgcA | N823 | | This study |
| ΔsrlE | N824 | | This study |
| ΔascF | N825 | | This study |
| ΔagaW | OO64 | | This study |
| ΔbglF | OO67 | | This study |
| ΔsgcC | OO68 | | This study |
| ΔmanY | OO69 | | This study |
| ΔgalP | OO70 | | This study |
| ΔgatC | OO71 | | This study |
| ΔnagE | OO72 | | This study |
| ΔglvC | OO73 | | This study |
| Δudk | R060 | | This study |
| Δudp | R058 | | This study |
| GFP fusions: | | | |
| MG1655 + psc101 pcmtA-GFP | O537 | | This study |
| MG1655 Δcrp + psc101 pcmtA-GFP | O538 | | This study |

| | | | |
|---|---|---|---|
| MH1655 Δcra psc101 pcmtA-GFP | Q115 | | This study |
| MG1655 ΔcmtA + psc101 pcmtA-GFP | P315 |  | This study |
| MG1655 + psc101-btuBGFP | R051 | | This study |
| MG1655 + psc101 p1rrnB-GFP | R921 | | This study |
| MG1655 + psc101 pfruA-GFP | P115 | | This study |
| MG1655 big colonies | R796, R797, R798 | fusA 1779T>G , rplL 122_127delTAGCTG | This study |
| MG1655 small colony | R799 | fusA 2015C>T, rplL 122_127delTAGCTG | This study |
| MG1655 small colony | R800 | fusA 2011C>T | This study |
| Overexpressions P. aeruginosa: | | | |
| P. aeruginosa + p0 | P845 | | This study |
| P. aeruginosa pSEVA-238 mtlFGK+ | P847 | | This study |
| P. aeruginosa pSEVA-238 gtsB+ | P848 | | This study |
| P. aeruginosa pSEVA-238 oprB+ | P849 | | This study |
| P. aeruginosa pSEVA-238 PA2291 | R434 | | This study |
| P. aeruginosa pSEVA-238 fruA+ | S298 | | This study |
| Overexpression A. baumannii: | | | |
| A. baumannii pSEVA-238 p0 | R978 | | This study |
| A. baumannii pSEVA-238 fruA+ | R980 | | This study |
| Overexpression E. coli: | | | |
| MG1655 + pSEVA-238 p0 | O897 | | This study |
| pSEVA-238 cmtAB+ | P134 | | This study |
| pSEVA-238 fruBKA+ | P314 | | This study |
| pSEVA-238 malEFG+ | P132 | | This study |
| pSEVA-238 manXYZ+ | P313 | | This study |
| pSEVA-238 lamB+ | P311 | | This study |
| pSEVA-238 frwBC+ | P715 | | This study |
| pSEVA-238 mtlA+ | P716 | | This study |
| pSEVA-238 malX+ | P717 | | This study |
| pSEVA-238 mngA+ | P718 | | This study |
| pSEVA-238 glpTQ+ | P719 | | This study |
| pSEVA-238 bglF+ | P720 | | This study |
| pSEVA-238 ypdGH+ | P721 | | This study |
| pSEVA-238 bglH+ | P722 | | This study |
| pSEVA-238 treB+ | P723 | | This study |
| pSEVA-238 galP+ | Q144 | | This study |
| pSEVA-238 gatABC+ | Q145 | | This study |
| pSEVA-238 chiP+ | Q146 | | This study |
| pSEVA-238 chbCBA+ | Q147 | | This study |
| pSEVA-238 srlAEB+ | Q148 | | This study |
| pSEVA-238 acsF+ | Q149 | | This study |

| | | | |
|---|---|---|---|
| pSEVA-238 xylEFG+ | Q169 | | This study |
| pSEVA-238 nupC+ | Q872 | | This study |
| pSEVA-238 nupG+ | Q873 | | This study |
| pSEVA-238 frvAB+ | Q168 | | This study |
| pSEVA-238 ptsG+ | Q257 | | This study |
| ΔcmtA + pSEVA238 cmtA+ | Q322 | | This study |
| MG1655 ΔcmtA + pSEVA238 p0 | Q321 | | This study |

185

| **Clinical train number and stab** | Antibiotic resistance genes |
|---|---|
| 886 (anal) | - |
| 1193 (skin) | blaCTX-M-14_1, blaTEM-1B_1, blaOXA-1_1, ant(3'')-Ia_1, aac(3)-IId_1, aadA5_1, sul1_2, dfrA17_1, aph(3')-Ia_1, mph(A)_1, mph(A)_2, tet(A)_4, catA1_1 |
| 1195 (feces) | blaCTX-M-1_6, blaTEM-1B_1, aadA2_2, sul1_2, sul2_2, dfrA12_1, aph(3'')-Ib_5; aph(3')-Ia_1; aph(6)-Id_1, mph(A)_1; mph(A)_2, tet(A)_4 |
| 1215 (skin) | ant(3'')-Ia_1, aac(6')-IIc_1, sul1_2, tet(A)_4 |
| 1238 (feces) | ant(2'')-Ia_18; ant(3'')-Ia_1, sul1_2, sul2_2, aph(3'')-Ib_5; aph(6)-Id_1 |
| 1236 (skin) | blaTEM-1B_1, blaSHV-2_2, aadA5_1, sul1_2, sul2_2, sul2_2 |
| Ec019 (anal) | blaCTX-M-14, mdfA_(1) |
| Ec120 (anal) | mdfA_(1), ant(2'')-Ia_1, ant(3'')-Ia_1, sul1_5, sul2_2 |
| Ec068 (colostomy) | mdf(A)_1 |

186

| Plasmid | Number | Construction |
|---|---|---|
| pSEVA-238 | M027 | |
| psc101 | O896 | |
| Overexpression plasmids for E. coli: | | |
| pSEVA-238 cmtAB+ | P134 | PCR on gDNA using primers ML 290/291. Ligation into pSEVA-238 between EcorI and XbaI restriction sites. |
| pSEVA-238 fruBKA+ | P314 | PCR on gDNA using primers ML 295/296. Ligation into pSEVA-238 between EcorI and XbaI restriction sites. |
| pSEVA-238 malEFG+ | P132 | PCR on gDNA using primers ML 299/300. Ligation into pSEVA-238 between EcorI and XbaI restriction sites. |
| pSEVA-238 manXYZ+ | P313 | PCR on gDNA using primers ML 301/302. Ligation into pSEVA-238 between EcorI and XbaI restriction sites. |
| pSEVA-238 lamB+ | P311 | PCR on gDNA using primers ML 297/298. Ligation into pSEVA-238 between EcorI and XbaI restriction sites. |

| | | |
|---|---|---|
| pSEVA-238 frwBC+ | P715 | PCR on gDNA using primers ML 311/312. Ligation into pSEVA-238 between EcorI and XbaI restriction sites. |
| pSEVA-238 mtlA+ | P716 | PCR on gDNA using primers ML 303/304. Ligation into pSEVA-238 between EcorI and XbaI restriction sites. |
| pSEVA-238 malX+ | P717 | PCR on gDNA using primers ML 315/316. Ligation into pSEVA-238 between KpnI and XbaI restriction sites. |
| pSEVA-238 mngA+ | P718 | PCR on gDNA using primers ML 307/308. Ligation into pSEVA-238 between EcorI and XbaI restriction sites. |
| pSEVA-238 glpTQ+ | P719 | PCR on gDNA using primers ML 317/318. Ligation into pSEVA-238 between EcorI and XbaI restriction sites. |
| pSEVA-238 bglF+ | P720 | PCR on gDNA using primers ML 323/324. Ligation into pSEVA-238 between EcorI and XbaI restriction sites. |
| pSEVA-238 ypdGH+ | P721 | PCR on gDNA using primers ML 309/310. Ligation into pSEVA-238 between EcorI and XbaI restriction sites. |
| pSEVA-238 bglH+ | P722 | PCR on gDNA using primers ML 321/322. Ligation into pSEVA-238 between KpnI and XbaI restriction sites. |
| pSEVA-238 treB+ | P723 | PCR on gDNA using primers ML 313/314. Ligation into pSEVA-238 between KpnI and XbaI restriction sites. |
| pSEVA-238 galP+ | Q144 | PCR on gDNA using primers ML 339/349. Ligation into pSEVA-238 between XbaI and PtsI restriction sites. |
| pSEVA-238 gatABC+ | Q145 | PCR on gDNA using primers ML 343/344. Ligation into pSEVA-238 between EcorI and XbaI restriction sites. |
| pSEVA-238 chiP+ | Q146 | PCR on gDNA using primers ML 333/334. Ligation into pSEVA-238 between EcorI and XbaI restriction sites. |
| pSEVA-238 chbCBA+ | Q147 | PCR on gDNA using primers ML 331/332. Ligation into pSEVA-238 between EcorI and XbaI restriction sites. |
| pSEVA-238 srlAEB+ | Q148 | PCR on gDNA using primers ML 337/338. Ligation into pSEVA-238 between EcorI and XbaI restriction sites. |
| pSEVA-238 acsF+ | Q149 | PCR on gDNA using primers ML 335/336. Ligation into pSEVA-238 between EcorI and XbaI restriction sites. |
| pSEVA-238 xylEFG+ | Q169 | PCR on gDNA using primers ML 325/326. Ligation into pSEVA-238 between EcorI and XbaI restriction sites. |
| pSEVA-238 nupC+ | Q872 | PCR on gDNA using primers ML 349/350. Ligation into pSEVA-238 between EcorI and XbaI restriction sites. |
| pSEVA-238 nupG+ | Q873 | PCR on gDNA using primers ML 351/352. Ligation into pSEVA-238 between EcorI and XbaI restriction sites. |
| pSEVA-238 frvAB+ | Q168 | PCR on gDNA using primers ML 327/328. Ligation into pSEVA-238 between EcorI and XbaI restriction sites. |
| pSEVA-238 ptsG+ | Q257 | PCR on gDNA using primers ML 345/346. Ligation into pSEVA-238 between EcorI and XbaI restriction sites. |
| Overexpression plasmids for P. aeruginosa: | | |
| pSEVA-238 mtlFGK+ | P828 | PCR on gDNA using primers ML 365/366. Ligation into pSEVA-238 between BamHI and XbaI restriction sites. |
| pSEVA-238 gtsB+ | P829 | PCR on gDNA using primers ML 355/356. Ligation into pSEVA-238 between EcoRI and XbaI restriction sites. |
| pSEVA-238 oprB+ | P831 | PCR on gDNA using primers ML 359/360. Ligation into pSEVA-238 between EcoRI and XbaI restriction sites. |
| pSEVA-238 PA2291 | R433 | PCR on gDNA using primers ML 414/415. Ligation into pSEVA-238 between EcoRI and XbaI restriction sites. |

| | | |
|---|---|---|
| pSEVA-238 fruA+ | R963 | PCR on gDNA using primers ML 363/364. Ligation into pSEVA-238 between EcoRI and XbaI restriction sites. |
| Overexpression plasmids for A. baumannii | | |
| pSEVA-238 fruA+ | R962 | PCR on gDNA using primers ML 431/432. Ligation into pSEVA-238 between EcoRI and XbaI restriction sites. |
| Fusions with GFP: | | |
| psc101-pcmtAGFP | O501 | PCR on gDNA using primers ML 252/253. Ligation into pTOPO-TA. Digestion with EcoRI and subcloning into psc101. |
| psc101-p1rrnBGFP | R692 | Fruchard et al., 2022 (tgt) |
| psc101-pfruAGFP | P115 | PCR on gDNA using primers ML 261/262. Ligation into pTOPO-TA. Digestion with EcoRI and subcloning into psc101. |
| psc101-pbtuBGFP | R051 | PCR on gDNA using primers ML 411/412. Ligation into pTOPO-TA. Digestion with EcoRI and subcloning into psc101. |

| Number | Name | Sequence |
|---|---|---|
| ML 252 | FcmtA-5 | TCAATTATGTAATATGCATCACG |
| ML 253 | FcmtAGFP-3 | AGTTCTTCTCCTTTACGCATAAATTATCCTTATTTTATTT |
| ML 261 | FEcfruA-5 | GCAATTAGGAAAAATGGC |
| ML 262 | FEcfruA-3 | GTTCTTCTCCTTTACGCATAGTTCTCCTCTCTTGCTG |
| ML 290 | 5SEVAcmtAecoRI | GCGGAATTCATGCGGCTTAGTGATTATTT |
| ML 291 | 3SEVAcmtAxbaI | GCGTCTAGATCAGTGTTTATGTTCGGCGG |
| ML 296 | 5SEVAfruBKAecoRI | CGCGGAATTCATGTTCCAGTTATCCGTACAGGAC |
| ML 297 | 3SEVAfruBKAxbaI | CGCGTCTAGATTACGCTGCTTTCGCTACTGC |
| ML 297 | 5SEVAlamBecoRI | CGCGGAATTCATGATGATTACTCTGCGCAAAC |
| ML 298 | 3SEVAlamBxbaI | CGCGTCTAGATTACCACCAGATTTCCATCTGG |
| ML 299 | 5SEVAmalEFGecoRI | CGCGGAATTCATGAAAATAAAAACAGGTGCACG |
| ML 300 | 3SEVAmalEFGxbaI | CGCGTCTAGATTAACCTTTCACACCACCTGCCG |
| ML 301 | 5SEVAmanXYZecoRI | CGCGGAATTCGTGACCATTGCTATTGTTATAGG |
| ML 302 | 3SEVAmanXYZxbaI | CGCGTCTAGATTACAGTCCCAGCAGGCCGC |
| ML 303 | 5SEVAmtlAecoRI | CGCGGAATTCATGAATAAGAAGGTGTTAACCCTGTC |
| ML 304 | 3SEVAmtlAxbaI | CGCGTCTAGATACTTACGACCTGCCAGCAGTTGCAC |
| ML 307 | 5SEVAmngAecoRI | CGCGGAATTCATGGTATTGTTTTATCGGGCAC |
| ML 308 | 3SEVAmngAxbaI | CGCGTCTAGATTATGGCATTACGCCATCAG |

| | | |
|---|---|---|
| ML 309 | 5SEVAypdHGecoRI | CGCGGAATTCATGAGTAAGAAACTGATTGCC |
| ML 310 | 3SEVAypdHGxbaI | CGCGTCTAGATTACAGGCTATCGATTAACAATTTG |
| ML 311 | 5SEVAfrwBCecoRI | CGCGGAATTCATGAATGAGTTGGTGCAGATC |
| ML 312 | 3SEVAfrwBCxbaI | CGCGTCTAGATTAAGCGGTTTGCGCCAGGTG |
| ML 313 | 5SEVAtreBKpnI | CGCGGGTACCATGATGAGCAAAATAAACC |
| ML 314 | 3SEVAtreBxbaI | CGCGTCTAGATTAAACAATGTCCAGCGTGC |
| ML 315 | 5SEVAmalXKpnI | CGCGGGTACCATGACGGCGAAAACAGCACCG |
| ML 316 | 3SEVAmalXxbaI | CGCGTCTAGATTATGCCTGGACAGTATGCATCAGAC |
| ML 317 | 5SEVAglpTQecoRI | CGCGGAATTCATGTTGAGTATTTTTAAACCAG |
| ML 318 | 3SEVAglpTQXbaI | CGCGTCTAGATTACTCTTTATTAAGAAATTTTAC |
| ML 321 | 5SEVAbglHKpnI | CGCGGGTACCATGTTTAGACGAAATCTTATTAC |
| ML 322 | 3SEVAbglHxbaI | CGCGTCTAGATTACCACCAGATTTCAGCCTGGG |
| ML 323 | 5SEVAbglFecoRI | CGCGGAATTCATGACGGAGTTAGCCAGAAAAATAG |
| ML 324 | 3SEVAbglFxbaI | CGCGTCTAGATTAGCGAATGATGGATAACAGCG |
| ML 325 | 5SEVAxylFGHecoRI | GCGGAATTCATGAAAATAAAGAACATTCTAC |
| ML 326 | 3SEVAxylFGHxbaI | GCGTCTAGATCAAGAACGGCGTTTGGTTGC |
| ML 327 | 5SEVAfrvABecoRI | GCGGAATTCATGGCAGCTCTTACTGCAAGC |
| ML 328 | 3SEVAfrvABxbaI | GCGTCTAGAGACAAGCTCCTGTTGCGCGGCTTTC |
| ML 331 | 5SEVAchcBAecoRI | GCGGAATTCATGGAAAAGAAACACATTTATCTG |
| ML 332 | 3SEVAchCBAxbaI | GCGTCTAGATTATGCCTTCAGTTTTTCATGAAGC |
| ML 333 | 5SEVAchiPecoRI | GCGGAATTCATGCGTACGTTTAGTGGCAAACG |
| ML 334 | 3SEVAchiPxbaI | GCGTCTAGATCAGAAGATGGTGAATGGTGCG |
| ML 335 | 5SEVAascFecoRI | GCGGAATTCATGGCCAAAAATTATGCGGCGCTG |
| ML 336 | 3SEVAscFxbaI | GCGTCTAGATCAATTAAGACTTACTTCTTTGG |
| ML 337 | 5SEVAsrlAEBecoRI | GCGGAATTCATGATAGAAACCATTACTCATGG |
| ML 338 | 3SEVAsrlAEBxbaI | GCGTCTAGATTACTCCTTAACAGATTCAAACTTC |
| ML 339 | 5SEVAgalPxbaI | GCGTCTAGAATGCCTGACGCTAAAAAACAGG |
| ML 340 | 3SEVAgalPptsI | GCGCTGCAGTTAATCGTGAGCGCCTATTTCG |
| ML 341 | 5SEVAglvCBecoRI | GCGGAATTCATGCTCAGTCAAATTCAACGC |
| ML 342 | 3SEVAglvCBxbaI | GCGTCTAGATGCCTCCGTAATGGCAACATTTTCTG |

| | | |
|---|---|---|
| ML 343 | 5SEVAgatABCecoRI | GCGGAATTCATGACTAACCTGTTTGTTCG |
| ML 344 | 3SEVAgatABCxbaI | GCGTCTAGATTATTCTGCGAGAACGACTTTCTC |
| ML 345 | 5SEVApstGecoRI | GCGGAATTCATGTTTAAGAATGCATTTGCTAACC |
| ML 346 | 3SEVAptsGxbaI | CGCGTCTAGATTAGTGGTTACGGATGTACTCATCC |
| ML 349 | 5SEVAnupCEcoRI | GCGGAATTCATGGACCGCGTCCTTCATTTTGTAC |
| ML 350 | 3SEVAnupCXbaI | CGCGTCTAGATTACAGCACCAGTGCTGCGATTGAC |
| ML 351 | 5SEVAnupGEcoRI | GCGGAATTCATGAATCTTAAGCTGCAGCTGAAAATC |
| ML 352 | 3SEVAnupGXbaI | CGCGTCTAGATTAGTGGCTAACCGTCTGTGTGCCTG |
| ML 355 | 5SEVAPagtsBEcorI | CGCGGAATTCATGGCGACCAATTCCCC |
| ML 356 | 3SEVAPagtsBxbaI | CGCGTCTAGATCACGCATGGCGCTTGCC |
| ML 357 | 5SEVAPPalamBEcorI | CGCGGAATTCATGAACAACCTGTTGTTG |
| ML 358 | 3SEVAPalamBxbaI | CGCGTCTAGATCATGCCGCCTCCAGGGCG |
| ML 359 | 5SEVAPaoprBEcorI | CGCGGAATTCATGTACAAGAACAAGAAAACC |
| ML 360 | 3SEVAPoprBxbaI | CGCGTCTAGATCAGAACACCGTCTGGATC |
| ML 363 | 5SEVAPafruopEorI | GCGGAATTCATGCTCGAACTCGATACCC |
| ML 364 | 3SEVAPafruopxbaI | GCGTCTAGAGTTATCCGCCGGTTCTCCGGAGT |
| ML 365 | 5SEVAVamalopBamHI | CGCGGGATCCATGAACGACTCGATCAAGGC |
| ML 366 | 3SEVAPamalopXbaI | CGCGTCTAGATCAGGCCGCCTGTTGCAGTC |
| ML 411 | FEcbtuB-5 | GAGCTGACGCGCAGCGGTAAG |
| ML 412 | FEcbtuB-3 | GTTCTTCTCCTTTACGCATTGTAAAGCATCCACAATAG |
| ML 414 | 5SEVAPA2291ECorI | GCGGAATTCGTGAAATCCCATCTTCTCCG |
| ML 415 | 3SEVAPA2291xbaI | GCGTCTAGACTAGAACACCGTCTGGATCTTG |
| ML 431 | 5SEVAAbfruAecoRI | GCGGAATTCCGAATCCCTACTTCTGAGGTTC |
| ML 432 | 3SEVAAbfruAXbaI | GCGTCTAGACCTGAGCAAATGATGGCCCGTG |
